# A DNA nanoscope that identifies and precisely localizes over a hundred unique molecular features with nanometer accuracy

**DOI:** 10.1101/2020.08.27.271072

**Authors:** Nikhil Gopalkrishnan, Sukanya Punthambaker, Thomas E. Schaus, George M. Church, Peng Yin

**Affiliations:** Wyss Institute for Biologically Inspired Engineering, Harvard University, Boston, MA 02115, USA; Department of Systems Biology, Harvard Medical School, Boston, MA 02115, USA; Department of Genetics, Harvard Medical School, Boston, MA 02115, USA

## Abstract

Techniques that can both spatially map out molecular features and discriminate many targets would be highly valued for their utility in studying fundamental nanoscale processes. In spite of decades of development, no current technique can achieve both nanoscale resolution and discriminate hundreds of targets. Here, we report the development of a novel bottom-up technology that: (a) labels a sample with DNA barcodes, (b) measures pairwise-distances between labeled sites and writes them into DNA molecules, (c) reads the pairwise-distances by sequencing and (d) robustly integrates this noisy information to reveal the geometry of the underlying sample. We demonstrate our technology on DNA origami, which are complex synthetic nanostructures. We both spatially localized and uniquely identified over a hundred densely packed unique elements, some spaced just 6 nm apart, with an average spatial localization accuracy (RMS deviation) of ~2 nm. The bottom-up, sequencing-enabled mechanism of the DNA nanoscope is fundamentally different from top-down imaging, and hence offers unique advantages in precision, throughput and accessibility.

## Introduction

The study of complex materials with nanoscale features benefits from forming an image of it, if possible, with increasingly sophisticated instruments which provide molecular-level detail for further understanding or validation. Comprehensive visualization can be challenging for two reasons – size and molecular diversity.

The finest molecular details can only be resolved with nanoscale localization. At the same time there is a tremendous diversity of molecular targets, necessitating the ability to identify and discriminate between these functional components (Fig. 1a). We propose a novel technique, which we term a DNA nanoscope (Fig. 1b), that tags targets with synthetic DNA barcodes, measures distances between many target pairs using biochemical DNA reactions, and then reconstructs a detailed map of the underlying geometry that uniquely identifies every target. We developed and demonstrated the capabilities of the DNA nanoscope technique on ‘DNA origami’^1^, which are complex synthetic nanostructures composed of hundreds of unique components.

**Fig. 1.**
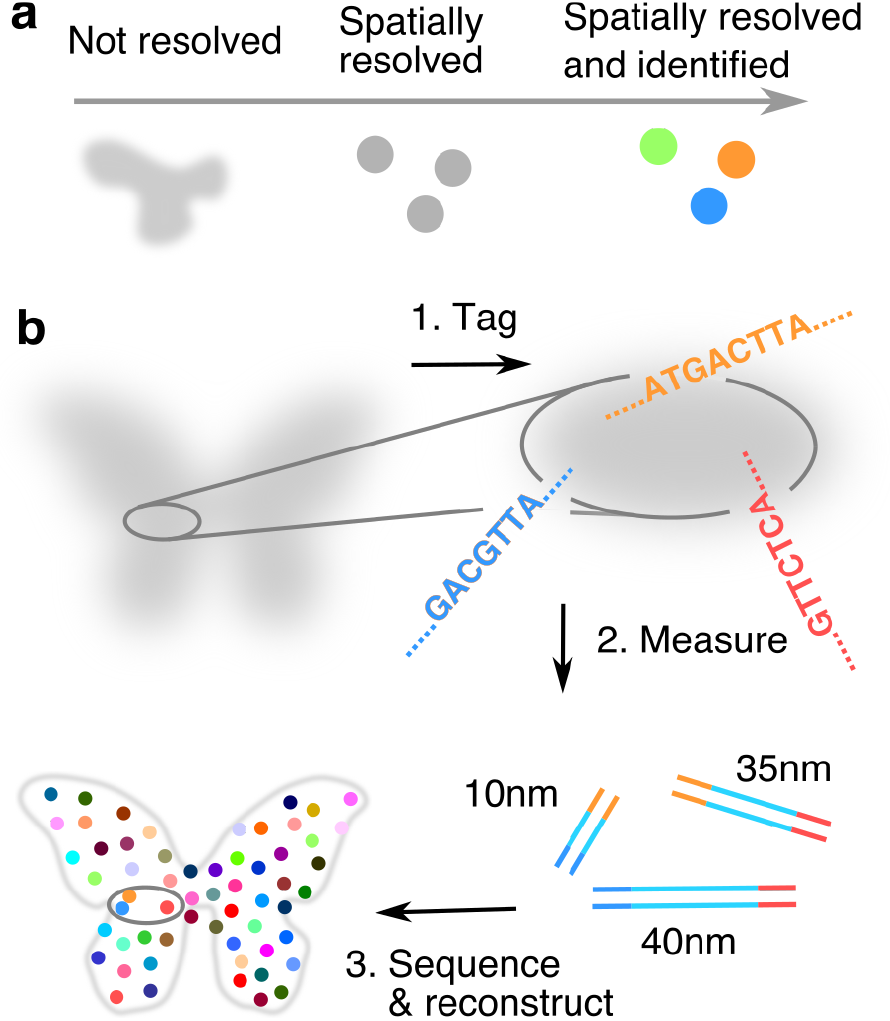
The DNA nanoscope ‘imaging-by-sequencing’ can both distinguish many targets and resolve features at the nanometer scale. **a.** A comprehensive visualization requires that we simultaneously resolve targets spatially and also determine their identity. **b.** Bird’s eye view of the DNA nanoscopy process. We tag targets with unique DNA barcodes, measure distances between many target pairs using DNA molecules, read the distances with massively parallel sequencing and integrate them into a molecular resolution spatial map that uniquely discriminates every target.

Like the DNA nanoscope, long-established and widely used class-average tools like X-ray crystallography and cryoEM also exploit sample periodicity or particle homogeneity to obtain nanometer or even angstrom resolution class-average reconstructions. However, they produce monochromatic images and can only discriminate molecular targets when they are resolved to near atomic precision, unachievable for many samples. At the other end of the spectrum, biochemical techniques like Hi-C can discriminate millions of DNA targets on chromosomes by sequencing them, however the contact densities currently obtainable from single nuclei Hi-C experiments preclude synthesizing this information into a structural model of the chromosome, while geometric models synthesized using data from ensemble Hi-C experiments have at best a local resolution of 5 kilobase pairs^6,7^.

This fledgling, ‘imaging-by-sequencing’ field^8–12^ has had two main experimental results. Our previous ‘auto-cycling proximity recording’ (APR)^8^ effort demonstrated seven-point reconstructions (spaced ~30 nm apart) from simple, binary proximity data. The subsequent ‘DNA microscope’, a reaction-diffusion scheme, demonstrated thousands of single particle localizations but only ~10 μm resolution^9^. In contrast to these previous attempts, our DNA nanoscope leverages the particle homogeneity of DNA origami to produce a nanoscale-resolution spatial map of a hundred or more points by making thousands of independent, pairwise distance measurements. This fine resolution is a direct consequence of two novel features of our molecular mechanism. First, our pairwise measurements report distance with high precision, as against previous efforts, which lacked precise distance reporting. Second, the DNA nanoscope measures and reports distances in the 10 nm to 100 nm range that is most relevant to molecular assemblies. This allowed us to resolve large gaps between components, situate otherwise disconnected clusters of points, and build a nanoscale precise global spatial map of the underlying geometry. Simulations showed (see Supplementary Fig. 1) that recording proximities with long reach but little distance precision resulted in maps in which points collapsed into unresolved clumps. Conversely, short reach but higher precision in distance measurements could resolve local geometry but not correctly situate distant points. However, when reach exceeded all gaps between adjacent points and the precision of distance measurement neared ~10%, reconstructions became surprisingly complete and accurate.

In this work, we pooled together information from many identical (up to manufacturing imperfections) copies of DNA origami to construct a class-average image. DNA origami structures have previously been characterized by AFMs^1^, EMs^2,3^ and super-resolution microscopes^4,5^. However, unlike the DNA nanoscope, none of these techniques can uniquely identify the over one hundred sequence-specific features that make up a typical DNA origami. In the final section we discuss how the DNA nanoscope can be extended from an ensemble technique to a general technique that does not rely on particle homogeneity.

## Results

### Encoding distances in DNA molecules

At the heart of the DNA nanoscope is a molecular ‘*ruler*’ mechanism that measures the distance between a pair of DNA-labeled targets and encodes it in a double stranded DNA molecule, which we call a ‘*distance record’*. The length of the distance record, in base pairs, directly corresponds to the physical distance measured. The molecular ruler mechanism consists of three stages – growth (Fig. 2a and Supplementary Fig. 2), connection and release (Fig. 2b and Supplementary Fig. 2). Given targets tagged with DNA handles, recording primers are introduced which bind to the handles via hybridization.

**Fig. 2.**
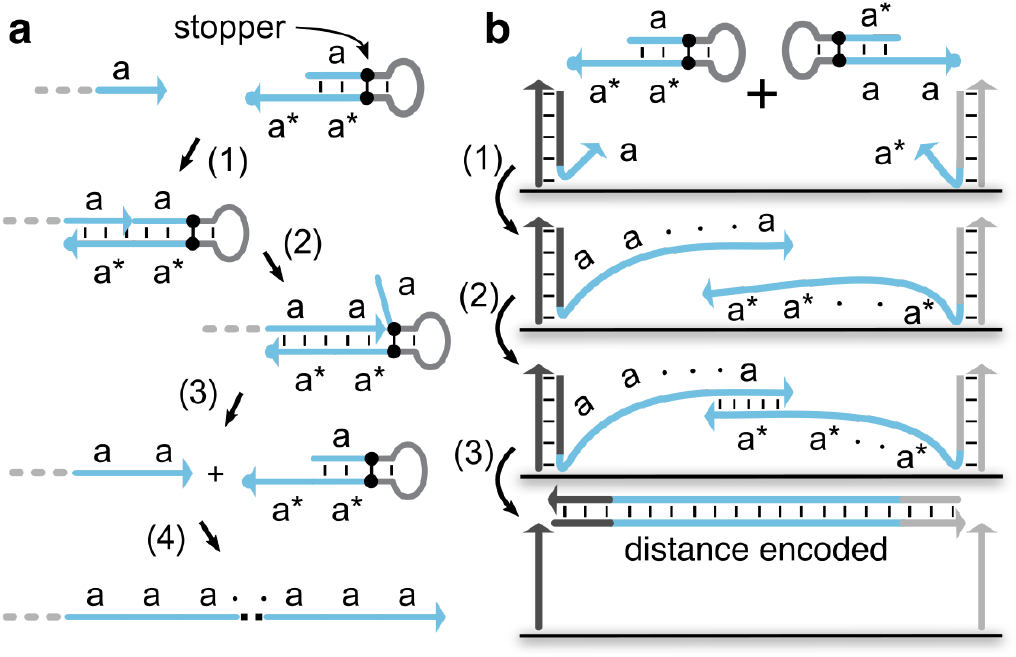
Molecular ruler mechanism (simplified). **a.** A primer exchange reaction (PER) cascade repeatedly adds the four base sequence domain ‘a’, as follows. (1) The recording primer hybridizes to a PER hairpin, (2) a strand displacing DNA polymerase (Bsm large fragment) extends the primer into the stem of the hairpin and in the process copies domain ‘a’. A ‘stopper’, a non-canonical base modification on the template that is not recognized by the DNA polymerase, blocks further extension. The polymerase dissociates from the hairpin. (3) The recording primer is only weakly bound to the hairpin and also dissociates. (4) The above sequence of reactions repeat, adding domain ‘a’ every time. In the same manner, a complementary PER cascade, shown in Supplementary Figure 2, repeatedly adds the four base sequence domain ‘a*’. **b.** A double-stranded DNA ‘distance record’ is generated as follows. Consider two DNA labeled targets with recording primers hybridized to them. (1) The primers take part in PER reaction cascades, as described in part A, adding sequence repeats of ‘a’ and ‘a*’ respectively. (2) The extended primers hybridize, (3) copy each other with the aid of the polymerase, are displaced from the targets and released into solution, making a distance record. The molecular ruler mechanism depicted here is a simplification. The full, actual mechanism is depicted in Supplementary Fig 2‥ See Supplementary Note 1 for the rationale for our molecular design choices.

Recording primers come in two complementary flavors, with either the sequence domain ‘a’ or the domain ‘a*’ (the reverse complement of ‘a’) at their 3’ ends. The recording primers, with the aid of a corresponding extension hairpin, a strand displacing DNA polymerase (Bsm large fragment) and dNTPs, undergo polymerase exchange reactions (PER)^13^ which repeatedly add the single-stranded domain sequence ‘a’ or ‘a*’ to their 3’ end (Fig. 2a and Fig. 2b(1)). Once complementary extended recording primers are long enough, they hybridize to each other via the domains ‘a’ and ‘a*’ (Fig. 2b(2)). At this point, again with the aid of a polymerase and dNTPs, the extended recording primers use each other as templates and polymerize to produce a double stranded DNA molecule that is displaced into solution (Fig. 2b(3)). The DNA molecule is a distance record, with the repeat domain ‘a a…a’ sandwiched between handle domains. The length of the repeat domain directly corresponds to the physical distance being measured. The sequence of the handle domain can serve as a DNA barcode to uniquely encode the molecular identity of the target. This process of growth, connection and release is isothermal and autonomous.

We wish to stress that each record molecule is a distance measurement, unlike the DNA microscope^9^ that encodes distance non-linearly in the *number* of identical proximity-dependent records produced. Of course, not every measurement of the same distance produces a distance record of the same length, because the growing single-stranded recording primers are entropic springs and their growth process is stochastic. However, repeated, independent measurements of the same distance can be aggregated to ultimately provide a ~1 nm distance measurement accuracy. In this work, our targets were positions on a DNA origami^1^, a well-characterized nanoscale breadboard. We made repeated measurements by having many identical to manufacturing imperfections) copies of DNA origami in the same reaction pot.

### Calibration of the molecular ruler

The molecular ruler mechanism produces distance records, whose length, in base pairs, must be related to the physical distance, in nanometers. We performed this translation by means of a calibration experiment. We placed molecular targets at known distances and performed DNA nanoscope recordings, which produced distance records that were then, in aggregate, related to the known distance, yielding a calibration function.

The calibration experiment was performed on a DNA origami adhered to a flat surface. The origami is composed of planar, parallel DNA double helices. The helices are held together by *staple* strands that cross over between neighboring helices at regular intervals along the helical axis. Each staple strand is uniquely addressable by way of its sequence and can be extended into handle domains that serve as handles for recruiting recording primers. We fixed the position of one of the targets near one end of one of the helices of the origami and offset the other target at regular intervals along this helix (Fig. 3a). The handle extended away from the surface and recording primers were bound to it by hybridization. The distance between targets can be calculated simply as the rise per base pair (= 0.34 nm) times the number of base pairs that separate them. The experimental workflow for a calibration experiment was as follows.

**Fig. 3.**
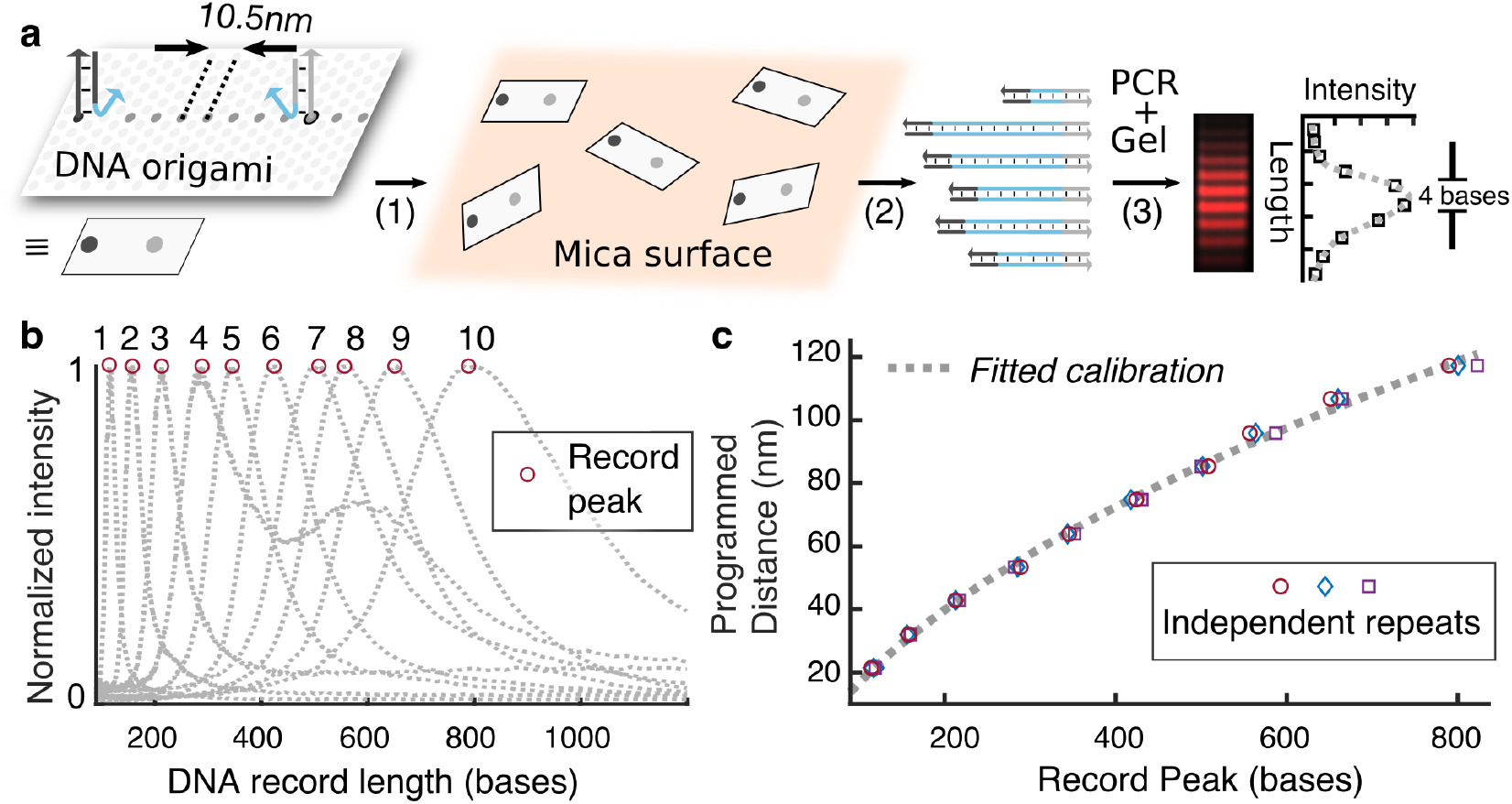
Molecular ruler calibration. **a.** DNA origami is used as a calibration standard. (1) DNA origami is deposited on a mica surface, (2) ruler recording generates distance records, (3) which are amplified by PCR and characterized by gel electrophoresis, which reveals a skew-normal distribution of record lengths. The discrete bands are 4 bases apart. See Materials and Methods for details on origami design and purification, ruler reaction conditions and PCR and gel protocols. **b.** Gel profiles of record lengths obtained from molecular recordings for various distances between target pairs. The programmed calibration distances are 1 = 21.4 nm, 2 = 32.0 nm, 3 = 42.8 nm, 4 = 53.4 nm, 5 = 63.9 nm, 6 = 74.8 nm, 7 = 85.3 nm, 8 = 95.9 nm, 9 = 106.8 nm and 10 = 117.3 nm. Each profile is normalized to its peak height. Bigger distances produce longer records that are more broadly distributed. The plotted DNA record lengths include primer regions of 32 bases each at either end. **c.** A calibration curve is fit to the peak of the distribution, giving us a function to transform a distance record, in base pairs, into a physical distance, in nanometers. The gel image and corresponding gel profiles for all three independent repeats can found in Supplementary Fig 3.).

The DNA origami was deposited (Fig. 3a(1)) onto a charged, atomically flat mica surface to minimize any flexibility due to thermal motion and reduce variability from one molecular measurement to the next. Reaction components (extension hairpins, strand displacing polymerase and dNTPs) were flown in and a DNA nanoscope recording was performed (Fig. 3a(2)). The distance records produced by this recording were collected, amplified by PCR and characterized by gel electrophoresis (Fig. 3a(3)). The distribution of lengths obtained reflects various independent measurements of the same distance. Our experiments showed that the distance records were skew-normal distributed (Fig. 3b). The greater the distance being measured, the longer, on average, were the distance records produced. The spread of the distribution also widened with increasing distance. We chose the location of the peak of the distribution, which is the most frequently produced distance record, as the representative for the increasing function (*a*√*x* + *b*) was fit to the peak data to generate a calibration function (Fig. 3c) that translates distance records into distance measurements. The lengths of the distance records, characterized by next-generation nanopore sequencing, were in excellent agreement with gel electrophoresis measurements (Supplementary Fig. 4).

### Full-color, molecular scale reconstruction of complex patterns

Armed with a molecular ruler mechanism to produce distance records and a calibration function to convert those records into physical distances, we applied the DNA nanoscope to reconstruct patterns on a DNA origami surface. We uniquely labeled each target feature of the pattern using DNA barcodes; recorded pairwise distances between labeled targets and reconstructed the pattern with molecular resolution.

### Pattern design

A DNA origami surface can be abstracted as a hexagonal grid, where each grid point corresponds to the 3’ end of a staple strand. A pattern is simply some subset of points chosen from this grid. Fig. 4a(1) shows a Smiley face pattern. All grid points have unique identities associated with them, furnished by the specific DNA sequence of the staple strand at that location.

**Fig. 4.**
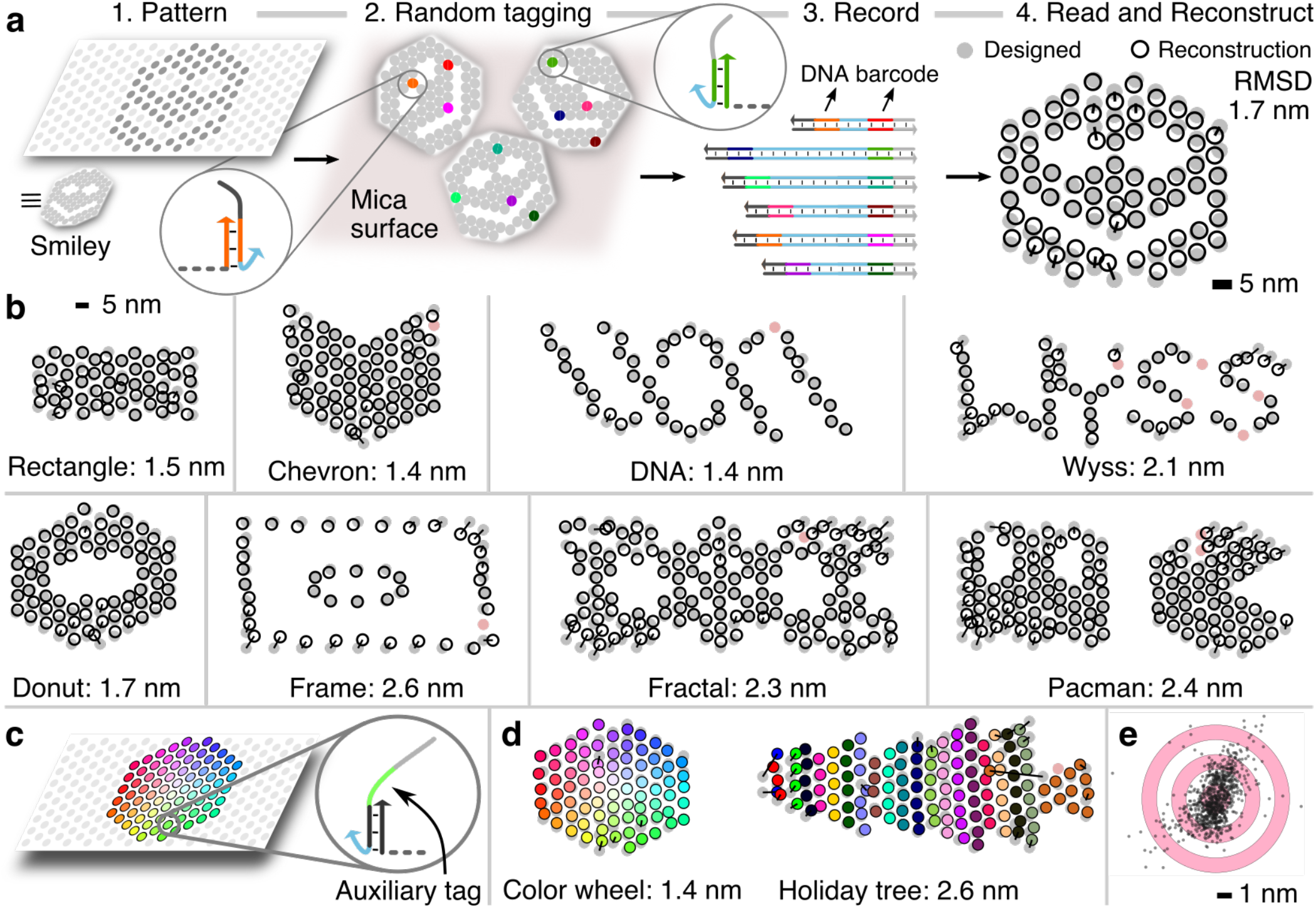
DNA nanoscope applied to various patterns **a. (**1) A pattern is some subset of positions on the DNA origami chosen from the underlying hexagonal grid. (2) A random subset of points is tagged with barcoded primers. While positions within each origami are only sparsely tagged, in aggregate each position is tagged many times over. (3) The identity of targets as well as the distance between targets is encoded inside barcoded distance records. (4) Distance records are read with next-generation sequencing to obtain length and barcode information, which is used to infer distances between points. An algorithm integrates pairwise distance measurements into a nanoscale precise map by embedding the points in a Euclidean plane. The reconstruction (hollow circles) is overlaid on the designed pattern (gray solid circles) for comparing the accuracy of the reconstruction. **b.** Many different patterns reconstructed with high accuracy. Each pattern is drawn to the same scale (scale bar = 5 nm). The numbers below the pattern are the RMS deviation between the designed and reconstructed pattern. Points missing from the reconstruction are indicated with red solid circles as opposed to gray solid circles. **c.** We encoded ‘color’ in auxiliary sequence tags that were then read out with the DNA nanoscope. **d.** Color wheel pattern with 77 distinct colors. Each auxiliary tag is unique. Holiday tree with 21 distinct colors. Each separate column of the pattern is a distinct auxiliary sequence, while points within the same column share the same sequence. All 13 points that make up the trunk share the same auxiliary sequence. **e.** An aggregate view of the accuracy of all the reconstructions from **b** and **d**. Each dot corresponds to the offset error vector between the reconstructed and the designed point. Each offset vector is translated to the center of the bulls-eye, whose each ring is 1 nm wide.

### Tagging

The DNA origami pattern is first prepared for recording by a tagging strategy that associates a barcode sequence with a staple strand. This barcode sequence, synthesized as a 3’ appendage on the corresponding staple strand, is used as a ‘handle’ to specifically recruit, via hybridization, a barcoded recording primer for a ruler measurement. We did not attempt to tag every feature of the pattern in every copy of the DNA origami. Instead we pursued a sparse tagging strategy, where every feature was randomly labeled with some probability and otherwise left unlabeled (Fig. 4a(2)). See Supplementary Materials and Methods for details on how this was achieved. In aggregate, across all the copies of the few hundred thousand DNA origami that were part of an experiment, we expect each feature of the pattern, on average, was tagged thousands of times.

### Recording

Once tagged, the DNA origami were deposited on a mica surface, as before, to reduce thermal molecular fluctuations and variations between origami copies. A molecular ruler recording was performed and distance records generated (Fig. 4a(3)). The distance records contained DNA barcodes at either end, corresponding to the underlying targets from which the measurement was produced.

### Reading distance records

Finally, both the lengths and barcode sequences of the distance records were read with next generation sequencing. Each distance record was parsed to identify its barcode sequences and then assigned to the target pair from which it was likely generated. The length distribution of distance records for a target pair reflects several independent measurements of the distance between them. The distribution was smoothed and the location of the most prominent peak extracted. The calibration function was used to translate this peak location into a physical distance measurement.

### Inferring geometry from distance data

The question of integrating noisy, pairwise distance measurements into an embedding in Euclidean space, referred to variously as the distance geometry problem^14^, global positioning problem^15^, localization problem^16^ etc., is well studied and has applications in sensor network localization, manifold learning and reconstruction of protein conformation from NMR data. Noisy distance measurements tend to end up in conflict with each other. The problem of producing an accurate embedding is thus a problem of balancing conflicting measurements.

The accuracy of a distance measurement between any two points depends on the number of corresponding distance records read. In fact, we found that the height of the most prominent peak, in units of number of reads, serves as an effective proxy for accuracy. In a calibration experiment we aggregated many thousands (sequencing) to millions (gel electrophoresis) of distance records from a single target pair, allowing us to precisely and accurately pinpoint the peak of the distribution. In contrast, a typical pattern reconstruction experiment only aggregates tens to hundreds of distance records per target pair, increasing uncertainty and producing less accurate distance measurements. Additionally, the multiplexed recording, amplification and sequencing process results in spurious reads, i.e. reads that likely come from unwanted side reactions. In cases where very few distance records are read from a target pair, these spurious reads exacerbate uncertainty and lead to highly inaccurate measurements (Supplementary Fig. 5). However, even with inaccurate data we managed to produce surprisingly accurate reconstructions by identifying less accurate measurements using peak heights and discounting them using weights.

Briefly, if the height of the most prominent peak is below a threshold parameter, the measurement is assigned a weight of 0. As the height exceeds the threshold, the assigned weight asymptotically approaches 1. A smaller weight reduces the influence of the corresponding measurement, allowing us to resolve conflicts in favor of more accurate measurements.

The value of the threshold parameter controls the relative influence of the measurements and changes the produced embedding. A threshold that is too low risks allowing too many inaccurate measurements to influence the embedding while a too high threshold risks discounting too many accurate measurements. The appropriate balance for the threshold parameter is auto-set by trying a series of values, each of which produces an embedding. Each embedding is evaluated for internal consistency, i.e. how well the embedding agrees with the measured distances. The threshold that produces the most internally consistent embedding is chosen as the optimal. Note that this auto-set procedure is without reference to any knowledge of the answer (see Supplementary Note 2C for details on how weights are calculated and Supplementary Note 2D for details on how the threshold parameter is auto-set).

In formal terms, we modeled the question as a global nonlinear optimization problem in two-dimensional Euclidean space. The objective function, which we seek to minimize, is defined as the weighted mean-squared-error between the measured and the embedded Euclidean distance. Simply attempting to minimize the objective function by starting from an arbitrary initial embedding was not robust. The optimization solution space is high-dimensional (2*n* where *n* is the number of points in the pattern) and highly non-convex, which resulted in the algorithm being trapped at local minima or saddle points. One solution to this issue would be to do repeated random initializations and pick the best instance. However, this is a computationally expensive approach.

Instead, we solved this optimization problem in three stages. First, points with very few (less than three) reliable (i.e. zero weight) measurements were dropped from the reconstruction, along with all their associated measurements (see Supplementary Note 2A). In the second stage, we used a robust facial reduction algorithm^17^ to obtain an initial solution that gave equal weight to all remaining measurements (see Supplementary Note 2B). Weights were then introduced and this initial solution was refined using a quasi-Newton algorithm to find the minimum of the objective function and thus obtain the final reconstructed pattern (see Supplementary Note 2C). The obtained embedding of the points was superimposed on the designed pattern using the Kabsch rigid transformation, which minimizes the root-mean-square deviation, a measure of the average distance between two paired sets of points. We emphasize that the patterns were reconstructed only using information from the pairwise distance records. No *a priori* knowledge of the geometry of the points was used to arrive at the final answer.

### Molecular resolution reconstructions

We successfully applied the DNA nanoscope technique to nine different patterns (Fig. 4) and obtained molecular resolution reconstructions. The root-mean-square deviation (RMSD) was used to quantify the average error between the designed and the reconstructed pattern. The RMSDs for the various patterns range from 1.4 nm to 2.6 nm. The points in the most densely packed patterns (Fig. 4b, Rectangle, Chevron, Donut and Pacman) are merely 6 nm apart and yet were clearly spatially resolved in the respective reconstructions. We successfully resolved negative space (Fig. 4a Smiley; Fig. 4b Donut, Frame, Fractal and Pacman), clusters of segregated points (Fig. 4b Frame, Wyss and Pacman) and sparse patterns (Fig. 4b Frame, DNA and Wyss). The variety of patterns reconstructed demonstrates the robustness of our approach. The biggest patterns were approximately 100 nm wide and 50 nm tall. The highest number of points localized was 135, in the Pacman pattern. Apart from spatially localizing the various points of a dense nanoscale pattern, the DNA nanoscope also uniquely distinguishes them by means of their barcode sequence, something that has proven unfeasible for microscopy techniques, which suffer from low-multiplexing capabilities.

### Full-color reconstructions

DNA origami has found wide use as a nanoscale breadboard and been decorated with receptor ligands^18^, gold nanoparticles^19–21^, quantum dots^22^, fluorescent dyes^23–25^ and carbon nanotubes^26^.

The attachment is usually mediated by using an auxiliary sequence tag. These auxiliary tags are independent of, and in addition to, the barcode tags associated with staple strands. Auxiliary tags allow us a programmable way to specify the geometry and absolute valency of objects decorated on DNA origami. Auxiliary tags could also be used to encode UMIs (unique molecular identities) that might help distinguish a particular DNA origami from its cohorts. We show that the DNA nanoscope can be used to natively read auxiliary tags in a multiplexed manner, demonstrating its power to discriminate molecular identity. We used a 12 base auxiliary sequence to encode ‘color’ information and reconstructed two patterns (Fig. 4c and Fig. 4d) that showcase our ability to read many ‘colors’ in arbitrary conformations.

### Robustness of reconstructions

As remarked earlier, the accuracy of our reconstructions exceed what one may naively expect from looking at the quality of the obtained raw data. For example, many distance measurements are significantly inaccurate (Supplementary Fig. 5) but the resulting reconstructions (Fig. 4e) are very accurate. In fact, we can tolerate further deterioration in data quality without suffering a severe loss of reconstruction accuracy. We again reconstructed patterns from the same experimental data as in Fig. 4, this time first deteriorating the data by one of four distinct methods to test how limited data quality is tolerated by the DNA nanoscope (Fig. 5).

**Fig. 5.**
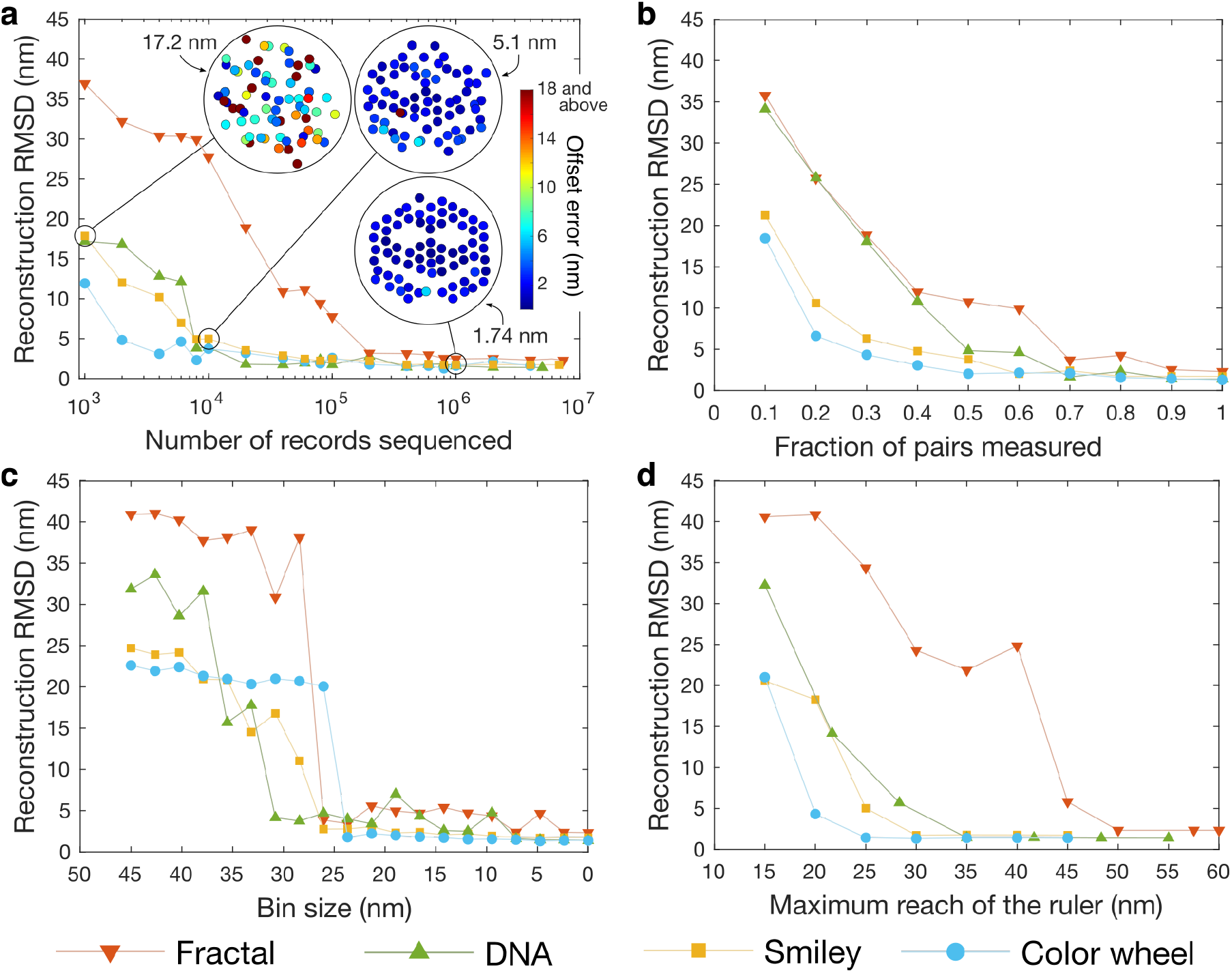
Reconstruction accuracy with deteriorating data quality for four representative patterns (Fractal, DNA, Smiley and Color wheel). **A.** We reconstructed patterns by successively sampling fewer and fewer sequencing reads. Even 10,000 sequence reads are sometimes sufficient to obtain ~5nm or better accuracy. Each plotted RMSD is an average of 10 independent samples. The inset shows example reconstructions. Each point is shaded by its error, which is defined as its offset distance from its designed position. **B.** The loss of some fraction of pairwise distance measurements, chosen at random from all possible pairs, is well tolerated by the DNA nanoscope. **C.** Distance measurements are binned to reduce precision. Some precision is necessary to reconstruct patterns with high accuracy, but the accuracy of the reconstruction does not significantly deteriorate with some loss of precision. On way of understanding the quantitative effect of bin size is to note that a bin size of *l* introduces an average error of *l* / 4 in a measurement, assuming a uniform distribution of distances within a bin. Thus, a bin size of 20nm would introduce an average error of 5nm in the measurements. **D.** All measurements between points farther apart than a maximum reach are discarded to demonstrate the effect of limited ruler reach. The closest spaced points in our patterns are 6 nm apart. We observe that a maximum reach limited to immediate neighbors fails to produce high-quality reconstructions. When reach extends beyond immediate neighbors, construction quality significantly improves. A reach extending to span the diameter of the pattern did not significantly improve reconstruction accuracy.

First, we reduced the number of records that were used to reconstruct a pattern by randomly sampling fewer and fewer DNA sequencing reads (Fig. 5a), consequently progressively deteriorating the accuracy of distance measurements. We found that 1 million total reads per pattern are sufficient to obtain ~2 nm RMSD for almost every pattern, and further sequencing did not significantly improve accuracy. Almost half the patterns achieve their optimal accuracy with as few as 100,000 reads (e.g. DNA, Smiley, Color wheel, Chevron and Rectangle). An RMSD of ~5 nm was obtained in some cases with as few as 10,000 reads (e.g. DNA, Smiley and Rectangle). A mere 2,000 reads sufficed to reconstruct the 77 point Color wheel with ~5nm RMSD.

We also tested the effects of an uneven deterioration in data by disregarding all distance reads between some fraction of pairs, resulting in the complete absence of respective distance measurements. We found that the random loss of up to 30% of the distance measurements is well tolerated (Fig. 5b) by most patterns and dense, compact patterns could tolerate the loss of almost 50% of measurements.

Third, as opposed to accuracy, we degraded the precision of the distance measurements by binning them. That is, we created equal sized distance bins (For instance [0 nm, 10 nm], [10 nm, 20 nm], and so on) and measurements that lay within each bin were approximated to the mid-point of that bin (5 nm, 15 nm and so on). Note that a bin size of *l* leads to an average perturbation of *l* / 4 in the distance measurements, assuming a uniform distribution of distances in a bin. A larger bin size corresponds to lower precision. We found that precision deteriorations corresponding to bin sizes of up to ~25 nm were well tolerated by the DNA nanoscope (Fig. 5c). In the limit of large bin sizes, we are effectively simulating a proximity-only measurement. Reconstructions fail to be accurate in these cases, demonstrating that in general a degree of precision, i.e. measuring distances and not just recording binary proximity, is critical for accurate reconstructions.

Finally, we confirmed our hypothesis that recording short-range distances that only span immediate neighbors does not produce accurate reconstructions. We simulated this limited ‘reach’ by discarding all measurements greater than a certain maximum and attempted to reconstruct patterns. We found that when reach extends beyond immediate neighbors (often sufficient to span the larger gaps), reconstruction accuracy improves significantly (Fig. 5d). This suggests that while individual distance measurements may mislead, there is “wisdom in the crowd”.

## Discussion

We have devised a DNA nanoscope, a tool that records molecular identities and spatial organization in DNA molecules with nanoscale localization accuracy. This DNA nanoscope was used to record dense, nanoscale patterns on homogenous DNA origami particles containing over a 100 unique features. Features spaced just 6 nm apart were clearly resolved with an average spatial localization accuracy of ~2 nm. Each feature was uniquely identified. This combination of spatial resolution and unique molecular identification is unprecedented even for homogenous particles, and has not been achieved by any other technology.

Bottom-up ‘imaging by sequencing’ technologies, like our DNA nanoscope, stand in contrast to top-down microscopy methods, and confer unique performance and operational advantages. The molecular recording processes that generate spatial data are isotropic and hence we expect that our 3D spatial resolution will match our demonstrated ~5 nm 2D spatial resolution as long as appropriate calibration standards are used. This is in contrast to microscopy methods that have worse 3D resolution in comparison to their 2D resolution. The recording ‘instruments’ of the DNA nanoscope are a swarm of molecules, diffusing throughout and inspecting a large population of molecular targets in parallel. This eliminates the need to correct any sample drift with respect to the instrument, which imposes practical and fundamental limits on the resolution of microscopy techniques. This parallelism also means that the recording throughput of a large sample is similar to that of smaller samples. The throughput is limited only by our ability to quickly sequence the records. Sequencing throughput is constantly improving and has seen Moore’s law like improvements in the past few years.

Apart from these performance advantages, the DNA nanoscope protocol has several operational advantages. First, there is no requirement that the sample be accessible to electromagnetic radiation, only to tiny diffusing DNA molecules that can likely penetrate to otherwise inaccessible locations. Second, the recording interactions with the sample are via gentle biochemical reactions (DNA hybridization and polymerization) in contrast to high-energy lasers, electron beams or physical probes used in super-resolution microscopy, electron microscopy and scanning probe microscopy respectively. Third, the sample does not need any special preparation, like adhering it to a surface, or freezing it in vitreous ice, that hold it immobile with respect to macro-scale recording instruments. The recording process is setup simply like a PCR reaction, except without any temperature cycling. Fourth, no capital-intensive, complex and hard to maintain instruments that are periodically rendered obsolete need to be purchased. A $1000 start-up kit available from a commercial source was used in this work. The per assay cost is currently high, costing about $500 per structure mapped in this work, but is seeing rapid drops in price as the technology continues to mature.

The bottom-up ‘imaging by sequencing’ field is nascent, and many challenges and opportunities remain. In this work, we exploited the homogeneity of DNA origami to reconstruct class average structures. The technique can potentially be extended to acquire images of the structure of single particles. Currently, our ruler recording is pairwise destructive and only one copy of a distance record can be generated from each labeled target. This precludes single particle reconstructions, as the resulting disjoint pairwise distances cannot be integrated into a spatial map. One solution is to combine pairwise non-destructive recording, as described in our previous ‘APR’^8^ work, with the ‘molecular ruler’ mechanism demonstrated here.

As we scale down to single particle reconstructions we scale up in the number of molecular features that we must resolve. In class average experiments, distinct physical copies of molecular targets are superimposed and treated as one target. In contrast, in single particle experiments, each physical copy will have to be treated separately. Thus, an experiment may have millions of unique targets. We argue that the DNA nanoscope technique will scale up to these numbers. Consider the case of a typical DNA origami experiment, with 50,000 DNA origami structures, each consisting of 50 points. These 2.5 million (50,000 times 50) targets can be labeled with unique DNA barcodes (there are a possible ~1 billion DNA sequences of length 20). Ruler recording reactions occur asynchronously and in parallel. Each ruler reaction at least a minute to produce a distance record^8^. Thus, a DNA origami can produce, on average, a few thousand distance records in a matter of hours. We have shown that 10,000 distance records per DNA origami proved sufficient to reconstruct them with sub-5nm accuracy. Improvements in the ruler mechanism that narrow the distribution of record lengths produced for each distance will further reduce the sequencing requirements. The total number of records that would need to be sequenced would be on the order of 500 million, already in reach of short-read sequencing technology and only an order of magnitude away from what long-read nanopore sequencing can currently achieve. The sequencing library could also be split, with shorter records sequenced on short-read high volume sequencers and the long-read nanopore devices focused on the longer reads.

We predict that the DNA nanoscope and related ‘imaging by sequencing’ techniques will gain widespread adoption over the next few years and drive fundamental nanoscale discoveries.

## Materials and Methods

A brief summary of the methods is provided here. Additional details may be found in the Supplementary Materials and Methods section.

### DNA origami manufacture and purification

The scaffold strand (M13mp18 single stranded DNA, 5 nM final concentration) was combined with: (i) all 216 ‘blunt’ staple oligos (50 nM final concentration of each oligo, see Supplementary Table 2 for sequences), the appropriate subset (depending of the pattern being tagged, see Supplementary Fig. 7, 8 and 9 and Supplementary Table 3) of barcoded ‘handle’ staple oligos (5 nM final concentration of each oligo) and corresponding appropriate subset of barcoded primers of type a and a* (5 nM final concentration of each oligo, see Supplementary Table 4 for sequences) in 1X TE Mg buffer (pH 7.4, 10 mM Tris-HCl, 0.1 mM EDTA, 10 mM MgSO_4_). The mixture was then cooled from 90°C to 60°C at the rate of 1 min/°C and then from 60°C to 50°C at the rate 10 min/°C and finally from 50°C to 25°C at the rate of 1 min/°C. Folded origami was stored at 4°C for up to one week prior to purification. DNA origami were purified by agarose gel electrophoresis to eliminate misfolded and aggregated origami as well as to remove excess staple and primer oligos.

### DNA nanoscope recording

A thin layer of mica was peeled from a mica sheet using sticky tape and then affixed to a sticky bottomless six-channel slide to assemble fluid-exchange reaction chambers for recording experiments. Purified DNA origami (50 μL at 50 pM) was added to the reaction chamber and allowed to bind to the mica surface for 10 min. The chamber was then washed twice with 50 μL of 1X TE Mg to remove unbound origami. The exposed, unbound mica surface is then passivated with a BSA solution (50 μg/mL) for 5 min and further washed with 1X TE Mg and a magnesium-supplemented 1X Thermopol buffer (20 mM Tris-HCl, 10 mM (NH_4_)_2_SO_4_, 10 mM KCl, 7 mM MgSO_4_, 0.1% Triton®-X-100, pH 8.8 @ 25°C). 50 μL of the recording mix, consisting of 100 nM extension hairpin type ‘a’, 100 nM extension hairpin type ‘a*’, 0.08 U/μL Bsm DNA polymerase LF, 100 μM dNTP solution mix, 1X Thermopol buffer and 5 mM MgSO_4_, is added to the reaction chamber and the slide kept at 37°C for 3 hours. After 3 hours, the supernatant containing distance records was aspirated and PCR amplified for further characterization.

### PAGE characterization

The length distribution of the distance records for each calibration distance was characterized by running PCR amplified distance records on a denaturing PAGE gel (180 V for 30 min at 50°C). Gel images were analyzed with the Fiji image processing software package.

### Next-generation sequencing and analysis

PCR amplified distance records were purified by denaturing polyacrylamide gel electrophoresis (see Supplementary Methods for details) to remove short-length spurious distance records. Purified distance records were prepared for next-gen sequencing using the Oxford Nanopore SQK-LSK109 ligation sequencing kit and sequenced to produce 10 to 15 million raw reads. We used Oxford Nanopore’s Guppy basecalling software (v3.2.1) to (1) read sequence information from raw sequencing data and then use MATLAB scripts to (2) demultiplex reads from different experiments, (3) extract the lengths of the distance records and assign them to their appropriate target-pair, (4) infer the distance for each target-pair from all assigned distance records, and finally (5) reconstruct the underlying geometry from pairwise distance measurements. The MATLAB scripts can be found at github.com/nikhil314/DNA-Nanoscope.

## Acknowledgments

We thank Prof. William M. Shih for useful discussion and comments on this work.

## Funding

This study is supported by grants to P.Y. by the Office of Naval Research (under grants N00014-16-1-2410 and N00014-18-1-2549), the National Institutes of Health (under grants 1R21CA235421-01, 5DP1GM133052-02 and 5R01GM124401-02), the National Science Foundation (under grants CBET-1729397 and MCB-1540214), the Defense Advanced Research Projects Agency (under grant W911NF-17-1-0075), and the Molecular Robotics Initiative at Wyss Institute

## Author contributions

N.G. conceived and designed the study, performed the experiments, developed the software, collected and analyzed the data, and wrote the manuscript. T.S. conceived the study, analyzed the data, and wrote the manuscript. S.P. performed nanopore sequencing experiments and collected data. G.M.C. provided scientific guidance and contributed to study supervision. P.Y. conceived and supervised the study and wrote the manuscript. All authors edited and approved the manuscript

## Competing interests

N.G., T.S. and P.Y. have filed a provisional patent covering aspects of this work. P.Y. is co-founder and director of Ultivue Global, NuProbe Inc. and Torus Biosystems; and

## Data and materials availability

The raw sequencing data files are many GB in size and are available on reasonable request. The code for performing the analysis and reconstructions is available on GitHub and a link to it can be found in the Materials and Methods section.

## Supplementary Materials available online

Supplementary Materials and Methods

Supplementary Notes 1 and 2

Supplementary Figures 1 to 10

Supplementary Tables 1 to 5

References (27-28)

## Notes

https://github.com/nikhil314/DNA-Nanoscope

## References

1. Rothemund, P. W. K. Folding DNA to create nanoscale shapes and patterns. Nature 440, 297–302 (2006).

2. Douglas, S. M. et al. Self-assembly of DNA into nanoscale three-dimensional shapes. Nature 459, 414–418 (2009).

3. Bai, X. C., Martin, T. G., Scheres, S. H. W. & Dietz, H. Cryo-EM structure of a 3D DNA-origami object. Proc. Natl. Acad. Sci. U. S. A. 109, 20012–20017 (2012).

4. Dai, M., Jungmann, R. & Yin, P. Optical imaging of individual biomolecules in densely packed clusters. Nat. Nanotechnol. 11, 798–807 (2016).

5. Iinuma, R. et al. Polyhedra self-assembled from DNA tripods and characterized with 3D DNA-PAINT. Science 344, 65–9 (2014).

6. Le Treut, G., Képès, F. & Orland, H. A Polymer Model for the Quantitative Reconstruction of Chromosome Architecture from HiC and GAM Data. Biophys. J. 115, 2286–2294 (2018).

7. Lesne, A., Riposo, J., Roger, P., Cournac, A. & Mozziconacci, J. 3D genome reconstruction from chromosomal contacts. Nat. Methods 11, 1141–1143 (2014).

8. Schaus, T. E., Woo, S., Xuan, F., Chen, X. & Yin, P. A DNA nanoscope via auto-cycling proximity recording. Nat. Commun. 8, 696 (2017).

9. Weinstein, J. A., Regev, A. & Zhang, F. DNA Microscopy: Optics-free Spatio-genetic Imaging by a Stand-Alone Chemical Reaction. Cell 178, 229–241.e16 (2019).

10. Boulgakov, A. A., Ellington, A. D. & Marcotte, E. M. Bringing Microscopy-By-Sequencing into View. Trends in Biotechnology 38, 154–162 (2020).

11. Boulgakov, A. A., Xiong, E., Bhadra, S., Ellington, A. D. & Marcotte, E. M. From Space to Sequence and Back Again: Iterative DNA Proximity Ligation and its Applications to DNA-Based Imaging. bioRxiv 470211 (2018). doi:10.1101/470211

12. Hoffecker, I. T., Yang, Y., Bernardinelli, G., Orponen, P. & Högberg, B. A computational framework for DNA sequencing microscopy. Proc. Natl. Acad. Sci. U. S. A. 116, 19282–19287 (2019).

13. Kishi, J. Y., Schaus, T. E., Gopalkrishnan, N., Xuan, F. & Yin, P. Programmable autonomous synthesis of single-stranded DNA. Nat. Chem. 10, 155–164 (2018).

14. Souza, M., Lavor, C., Muritiba, A. & Maculan, N. Solving the molecular distance geometry problem with inaccurate distance data. BMC Bioinformatics 14, (2013).

15. Singer, A. A remark on global positioning from local distances. Proc. Natl. Acad. Sci. U. S. A. 105, 9507–9511 (2008).

16. Javanmard, A. & Montanari, A. Localization from Incomplete Noisy Distance Measurements. Found. Comput. Math. 13, 297–345 (2013).

17. Drusvyatskiy, D., Krislock, N., Voronin, Y. L. & Wolkowicz, H. Noisy euclidean distance realization: Robust facial reduction and the pareto frontier. SIAM J. Optim. 27, 2301–2331 (2017).

18. Hawkes, W. et al. Probing the nanoscale organisation and multivalency of cell surface receptors: DNA origami nanoarrays for cellular studies with single-molecule control. Faraday Discuss. 219, 203–219 (2019).

19. Liu, W., Halverson, J., Tian, Y., Tkachenko, A. V. & Gang, O. Self-organized architectures from assorted DNA-framed nanoparticles. Nat. Chem. 8, 867–873 (2016).

20. Kuzyk, A. et al. DNA-based self-assembly of chiral plasmonic nanostructures with tailored optical response. Nature 483, 311–314 (2012).

21. Acuna, G. P. et al. Fluorescence enhancement at ocking sites of DNA-directed self-assembled nanoantennas. Science 338, 506–10 (2012).

22. Samanta, A., Zhou, Y., Zou, S., Yan, H. & Liu, Y. Fluorescence quenching of quantum dots by gold nanoparticles: A potential long range spectroscopic ruler. Nano Lett. 14, 5052–5057 (2014).

23. Gopinath, A., Miyazono, E., Faraon, A. & Rothemund, P. W. K. Engineering and mapping nanocavity emission via precision placement of DNA origami. Nature 535, 401–405 (2016).

24. Lin, C. et al. Submicrometre geometrically encoded fluorescent barcodes self-assembled from DNA. Nat. Chem. 4, 832–839 (2012).

25. Hemmig, E. A. et al. Programming Light-Harvesting Efficiency Using DNA Origami. Nano Lett. 16, 2369–2374 (2016).

26. Maune, H. T. et al. Self-assembly of carbon nanotubes into two-dimensional geometries using DNA origami templates. Nat. Nanotechnol. 5, 61–66 (2010).

