## Supplementary Materials for "A DNA nanoscope that identifies and precisely localizes over a hundred unique molecular features with nanometer accuracy"

**Supplementary Materials and Methods**

DNA origami design, manufacture and purification

**Design**: The rectangular DNA origami nanostructure used in all experiments was designed using the cadnano software (*27*), according to principles laid out in (*28*). The origami is 18 helices tall and 376 base pairs wide, and is 'twist corrected' (*28*) to promote flatness and minimize stress. Assuming a helix width of 2 nm and a helix to helix spacing of 1 nm, the structure's height is calculated to be 18*2 nm + 17*1 nm = 53 nm. Assuming a rise of 0.34 nm per base pair, the structure's width is calculated to be 376*0.34 nm = 127.84 nm. The origami is composed of a scaffold strand (M13mp18 single-stranded DNA, New England Biolabs (NEB), catalog no. N4040S) and 216 'blunt' staple oligos (Integrated DNA Technologies (IDT). See Supplementary Fig. 6 for the scaffold routing and Supplementary Table 2 for 'blunt' staple oligo sequences. Extended staple oligos (IDT) composed of a 'blunt' staple followed by a barcoded handle domain are used to recruit barcoded molecular ruler primers (IDT) via hybridization. The sequences for extended staple oligos are listed in Supplementary Table 3 and the sequences for barcoded molecular ruler primers are listed in Supplementary Table 4.

**Manufacture of randomly tagged DNA origami patterns:** Random sparse tagging was achieved by setting up a winner take all competition for staple incorporation at every target site, by supplying both the barcoded 'handle' staple and the corresponding barcode-less 'blunt' staple. At every target site, either the handle staple or the corresponding blunt staple can incorporate into the DNA origami, but not both. In any particular copy of the DNA origami, a site was successfully tagged when the handle staple won the competition, allowing recording primer recruitment at that site. Conversely, the site was not tagged when the blunt staple won the competition. The average density of tagged points on a DNA origami depends on the relative propensity of incorporation of these competing staples. The relative propensity, and hence the average density of tagged points on a DNA origami, was tuned by varying the relative concentrations of competing staples. The locations of the barcoded handle staples for each pattern are presented in Supplementary Fig. 7, 8 and 9.

The scaffold strand (5 nM final concentration) was combined with: (i) all 216 'blunt' staple oligos (50 nM final concentration of each oligo, see Table S2 for sequences), (ii) the appropriate subset (depending of the pattern being tagged, see Supplementary Fig. 7, 8 and 9 and Supplementary Table 3) of barcoded 'handle' staple oligos (5 nM final concentration of each oligo) and (iii) corresponding appropriate subset of barcoded primers of type a and a* (5 nM final concentration of each oligo, see Supplementary Table 4 for sequences) in 1X TE Mg buffer (pH 7.4, 10 mM Tris-HCl, 0.1 mM EDTA, 10 mM $MgSO_4$). The mixture was then cooled from 90 °C to 60 °C at the rate of 1 min/°C and then from 60 °C to 50 °C at the rate 10 min/°C and finally from 50 °C to 25 °C at the rate of 1 min/°C. Folded origami was stored at 4 °C for up to one week prior to purification.

**Purification:** DNA origami patterns were purified by agarose gel electrophoresis to eliminate aggregated structures and unbound oligos. 20 µL of folded DNA origami was mixed with 4 µL of 6X loading dye and loaded per lane in a 8 cm tall and 6 cm wide 1 % agarose gel (UltraPure agarose, Thermo Fisher Scientific, catalog no. 16500100) pre-stained with 1X SYBR Gold. The gel was run at 80 V for 2 hours on an Amersham instrument to separate well-folded DNA origami from excess oligos and aggregates. The band corresponding to well-folded DNA origami was visualized under a blue-light transilluminator (Safe Imager, Thermo Fisher Scientific) and excised with a clean razor blade. The gel slice was transferred to a freeze-n-squeeze DNA gel-extraction spin column (Bio-Rad, catalog no. 7326165), crushed with a clean pestle, kept at -20 °C for 10 min and then centrifuged at 2,000 g for 4 min. The flow through containing purified DNA origami was diluted 10-fold in 1X TE Mg buffer and 50 µL aliquots were stored at -20 °C for recording experiments.

<u>DNA nanoscope recording experiment</u>

**1. Prepare fluid-exchange reaction chamber:** A thin layer of mica of dimensions 25 mm X 75 mm is peeled from a mica sheet (Muscovite Mica V-1 Quality, Electron Microscopy Sciences, catalog no. 71855-05-10) using sticky tape (3M Scotch Clear Magic Tape, 25 mm wide). The mica sheet stuck to the tape is then affixed to a sticky bottomless six-channel slide (sticky-slide VI 0.4, ibidi, catalog no. 80608) to assemble fluid-exchange reaction chambers for recording experiments. Each channel is used to perform an independent DNA nanoscope recording experiment.

**2. Deposit origami on surface:** 50 µL of frozen DNA origami aliquot is thawed and heated to 42 °C for 2 min, to dissociate any aggregates. It is then added to a reaction chamber and allowed to bind to the mica surface for 10 min. The chamber is then washed twice with 50 µL of 1X TE Mg to remove unbound origami. The exposed, unbound mica surface is then passivated with a BSA solution (50 µg/mL, NEB) for 5 min and further washed with 1X TE Mg and a magnesiumsupplemented 1X Thermopol buffer (20 mM Tris-HCl, 10 mM $(NH_4)_2SO_4$, 10 mM KCl, 7 mM $MgSO_4$, 0.1% Triton®-X-100, pH 8.8 @ 25 °C, NEB, catalog no. B9004S).

**3. Perform molecular ruler reaction:** 50 µL of the recording mix, consisting of 100 nM extension hairpin type 'a', 100 nM extension hairpin type 'a*', 0.08 U/µL Bsm DNA polymerase LF (*ThermoFisher Scientific,* catalog no. EP0691), 100 µM dNTP solution mix (NEB, catalog no. N0447S), 1X Thermopol buffer and 5 mM $MgSO_4$, is added to the reaction chamber and the slide kept at 37 ºC for 3 hours. After 3 hours, the supernatant containing distance records is collected for further processing.

PCR amplification of distance records

Distance records were amplified by PCR. For PAGE analysis, the PCR reaction mix contained distance records (1 µL per 20 µL reaction mix), fluorescent PCR primer 1 (0.3 µM, IDT. See Supplementary Table 1 for sequence), fluorescent PCR primer 2 (0.3 µM, IDT. See Supplementary Table 1 for sequence), AccuPrime *Pfx* Reaction Mix (1X), AccuPrime *Pfx* DNA polymerase (0.025 U/µL, *ThermoFisher Scientific*, catalog no. 12344024) and EvaGreen fluorescent nucleic acid dye (1X, Biotium, catalog no. 31000-T). For next-gen sequencing analysis, the PCR mix contained inner primers (10 nM each, IDT), outer primers (0.3 µM each, IDT) and the other PCR mix components listed above. The outer primers are sequencing barcodes used to multiplex multiple libraries (i.e. distinct DNA nanoscope experiments) on the same sequencing chip. The inner primers contain sequences common to the ends of all barcoded recording primers. The sequence information for these can be found in table S5.

Temperature cycling for the PCR mix had (i) an initial denaturation step of 95 °C for 2 min, followed by (ii) 25 to 35 cycles of denaturation at 95 °C for 15s and primer binding and extension at 60 °C for 45s and (iii) a final extension at 65 °C for 2 min. The progress of the PCR amplification was monitored on a real time PCR machine. The number of temperature cycles was chosen to allow the signal to plateau, indicating completion of the PCR reaction.

PAGE characterization of distance records for calibration experiments

The length distribution of the distance records for each calibration distance was characterized using PAGE. 10 µL of PCR-amplified distance records were mixed with 10 µL of 2X denaturing loading dye (95 % formamide, 10 mM NaOH, 0.025 % bromophenol blue, 1 mM EDTA), heated to 95 °C for 2 min and then loaded into individual lanes of an 8 cm X 8 cm 4 % denaturing polyacrylamide gel (4 % v/v acrylamide/bis-acrylamide (19:1), 7.4 M Urea, 0.5X TBE). The gel was electrophoresed at a voltage of 180 V for 30 min at 50 °C. The gel was then stained with a SYBR Gold (1:10,000 v/v) solution for 10 min and then imaged on a Typhoon FLA 9500 gel scanner (General Electric) in the red (635 nm emission laser and 665 nm low-pass red filter) and blue channels (473 nm emission laser and 510 nm low-pass blue filter). Gel images were analyzed with the Fiji image processing software package.

Purification of distance records for next-gen sequencing

Distance records were purified to remove short-length spurious distance records prior to sample preparation for next-gen sequencing. The purification steps were as follows:

1. **Column concentration**: The post-PCR distance records were cleaned and concentrated using a bind-wash-elute procedure on a silica membrane column using the MinElute PCR purification kit (Qiagen, catalog no. 28004) and it's associated standard protocol. The input was 100 µL of post-PCR mix and the elution volume was 10 µL in buffer EB (10 mM Tris-Cl, pH 8.5).

2. **PAGE purification:** Concentrated samples were mixed with 10 µL of 2X denaturing loading dye (95 % formamide, 10 mM NaOH, 0.025% bromophenol blue, 1 mM EDTA), heated to 95 °C for 2 min and then loaded into individual lanes of an 8 cm X 8 cm 4 % denaturing polyacrylamide gel (4 % v/v acrylamide/bis-acrylamide (19:1), 7.4 M Urea, 0.5X TBE). The gel was electrophoresed at a voltage of 180 V for 60 min to 80 min at 50 °C to allow the short-length spurious distance records to diffuse out of the gel into the surrounding running buffer. The appropriate time to run the gel to allow this to happen was determined by monitoring, in adjacent lanes, the migration of fluorescent fiduciary bands that co-migrate with short-length spurious distance records.

3. **PAGE concentration:** After the short-length spurious distance records were discharged into the surrounding running buffer, the desired distance records, which range in size from ~150 b to ~1 kb and were distributed over a large gel volume, were concentrated by reversing the current flow, as follows. First, the running buffer was replaced to prevent the spurious records from re-entering the gel. Next, 50 µL of a dense 15 % denaturing polyacrylamide gel (15 % v/v acrylamide/bis-acrylamide (19:1), 7.4 M Urea, 0.5X TBE) was pipetted into each lane of the gel and allowed to polymerize. The voltage across the gel terminals was now reversed and the gel run at 180 V for 60 min to 80 min at 50 °C to allow the desired distance records to collect in the dense gel, concentrating them. The dense gel was excised with a razor blade. A clean 0.5 mL tube was pierced at the bottom with a 20-gauge syringe needle and placed into a 1.5 mL tube,

setting up a gel-shredder tube-in-tube widget. The excised gel from each lane was placed in the gel-shredder tube-in-tube widget and centrifuged at 14,000 g for 2 min. The disintegrated gel collects in the 1.5 mL tube, and the 0.5 mL tube is discarded. 50 µL of buffer EB is added to the 1.5 mL tube, which is then heated to 65 °C for 5 min followed by freezing at –20 °C for 10 min. The frozen pellet is transferred to a Freeze 'N Squeeze gel extraction spin column (Biorad, catalog no. 7326165) and centrifuged at 14,000 g for 3 min to collect ~ 50 µL of solution containing desired distance records. The solution is finally cleaned and concentrated using a bind-wash-elute procedure on a silica membrane column using the MinElute PCR purification kit (Qiagen, catalog no. 28004) and it's associated standard protocol. The final elution volume is 10 µL in buffer EB (10 mM Tris-Cl, pH 8.5). The concentration is measured on a NanoDrop 2000 (ThermoFisher) using OD260 absorption.

Library preparation for next-gen sequencing

Purified distance records were prepared for next-gen sequencing using the Oxford Nanopore SQK-LSK109 ligation sequencing kit. The protocol titled '*1D Genomic DNA by Ligation'* (see Oxford nanopore website) was used to prepare samples. A total of 0.5 pmole of purified distance records were used in the library preparation. Sometimes samples from two different experiments (i.e. distinct patterns) were combined, in which case we used 0.25 pmole of sample from each experiment. Briefly, the protocol involves a combined step of DNA repair and end-prep with the NEBNext FFPE DNA Repair Mix (New England Biolabs, catalog no. M6630S) and the NEBNext Ultra II End Repair / dA-Tailing Module (New England Biolabs, catalog no. E7546) respectively. This is followed by cleanup with the Agencourt AMPure XP beads (2X bead to sample ratio). Then, sequencing adaptors are ligated using the NEBNext Quick T4 DNA Ligase (New England Biolabs, catalog no. E6056) followed by a final round of bead clean-up. The sample is then mixed with loading beads, added to a 'primed' flow cell (FLO-MIN106) (see protocol for details) and sequenced for between 12 hours to 24 hours, producing around 10 million to 15 million raw reads. The sample can be split across multiple flow cells if more reads are required.

Data analysis workflow

We use Oxford Nanopore's Guppy basecalling software (v3.2.1) to (1) read sequence information from raw sequencing data and then use MATLAB scripts to (2) demultiplex reads from different experiments, (3) extract the lengths of the distance records and assign them to their appropriate target-pair, (4) infer the distance for each target-pair from all assigned distance records, and finally (5) reconstruct the underlying geometry from pairwise distance measurements. The MATLAB scripts referenced below can be found at github.com/nikhil314/DNA-Nanoscope. Details of the process are as follows:

1. **Basecalling using Guppy (v3.2.1):** Guppy is a base-calling software provided by Oxford Nanopore that can be downloaded and run on local machines. It takes raw sequencing data in the FAST5 file format and produces sequences in the FASTQ file format. Guppy was run on Windows Server 2008 machines with either 12 or 16 CPU cores. The process took about 40min to base-call 1 million reads on our computing setup. The command used to perform base-calling was:

```
guppy_basecaller.exe --input_path <input_path_fast5> --save_path
<output_path_fastq> --num_callers 12 --cpu_threads_per_caller 4 --
qscore_filtering -q 50000 -c dna_r9.4.1_450bps_fast.cfg
```

where:

`<input_path_fast5>` is the path of the raw reads in FAST5 format,

`<output_path_fastq>` is the path where base-called reads are stored in FASTQ format,

`--num_callers` specifies the number of parallel base-calling instances to run,

`--cpu_threads_per_caller` specifies the number of threads used by each instance of the base-calling software,

`--qscore_filtering` specifies that the reads be filtered by quality (default is to pass all reads with q_score above 7),

`-q` specifies number of reads to write per fastq file, and

`-c` specifies the configuration file.

2. **Demultiplex reads from different experiments:** The ends of the reads are scanned for sequencing barcodes (using local sequence alignment) and sorted into subdirectories based on barcodes identified. The MATLAB command used was:

```
sort_barcoded_reads(fastq_dir,ONT_barcodes)
```

where:

`fastq_dir` is the path of the fastq sequence files and

`ONT_barcodes` specifies the sequencing barcodes and their reverse complements.

The output is written into fastq files with reads that are sorted into sub-directories corresponding to the identity of the sequencing barcodes.

3. **Extract record lengths and assign to correct target pair:** Reads are scanned to identify the unique staple barcode sequences associated with each distance record. The length of the repeat region between the barcode sequences is extracted and assigned to the appropriate target pair. The following MATLAB command was executed for each directory containing reads from the same experiment:

```
pairwise_record_list = extract_pairwise_record_lengths(target_barcodes,path,
library_size, color_length)
```

where:

`target_barcodes` specifies the staple barcode sequences,

`path` specifies the path of the fastq reads,

`library_size` specifies the number of sequencing libraries that were combined for a single run,

`color_length` specifies the length of the auxiliary tag sequence, and

`pairwise_record_list` is the output, a matrix of size (n, n, 2001) where n is the number of target points and cell (i, j, k) holds the number of distance records of length k bases (only counting the repeat region) between points i and j. All distance records of length > 2000 are stored in the slice (:, :, 2001).

`pairwise_record_list` variables from different sequencing runs of the same experiment were combined by simply adding them.

4. **Infer the measured distance between every target pair:** The distribution of distance record lengths between every pair of points was examined to identify the major peak (in base pairs), which was then converted into a distance (in nanometers) by applying the calibration function. The MATLAB function used was:

```
[pairwise_distances, pairwise_peak_heights] =
finddist_geometry(pairwise_records_list,calibration_fun)
```

where:

`pairwise_records_list` is the output of extract_pairwise_record_lengths (see previous step),

`calibration_fun` is a cfit object that holds the calibration function of the ruler, which maps bases to nanometers,

`pairwise_distances` is one output, an array of measured distances for each pair of points, and `pairwise_peak_heights` is the peak height (in bases) corresponding to the distances measured. It is a measure of the confidence in the measurement.

5. **Reconstruct geometry from pairwise distance measurements:** The distance measurements are integrated into a coherent embedding of the targets in the 2D Euclidean plane, using the following MATLAB function:

```
[theta, prune, score] = solveDGP(pairwise_distances, pairwise_peak_heights,
opt_threshold)
```

where:

`pairwise_distances` and `pairwise_peak_heights` are the outputs from `finddist_geometry` (see previous step),

`opt_threshold` is a parameter used to prune unreliable measurements and to generate weights for measurements reflecting their reliability (see supplementary text for details on how `opt_threshold` is auto-set),

`theta` is a list of coordinates, specifying the final embedding,

`prune` is a logical-valued array indicating which target points were dropped from the final reconstruction, and

score is a measure of the internal consistency of the embedding. See supplementary text S2D.

The final embedding is compared to the designed pattern by superimposing them to minimize the RMSD (root-mean-square deviation). The MATLAB script used is:

```
[theta_translated, lrms] = superimpose(theta, theta_designed, prune)
```

where:

`theta` and `prune` are the outputs from `solveDGP`,

`theta_designed` contains a list of coordinates specifying the designed pattern,

`theta_translated` is the superimposition of the final embedding that minimizes the RMSD between the designed and reconstructed pattern, and

`lrms` is the corresponding RMSD.

**Supplementary Notes**

1. Design of molecular ruler primers and extension hairpins

Two main considerations went into the design of molecular ruler primers and extension hairpins. First, we reasoned that even weak secondary structure in the growing primers (‘a a …a’ and ‘a* a* … a*’) would result in a high propensity intra-molecular reaction where the primer folds back on itself. Normally such a state is transient and would resolve itself. However, in the presence of DNA polymerase in the molecular ruler reaction mixture, the primer could extend back on itself, proving fatal to the ruler reaction. This led us towards adopting a two letter code for the sequence domains ‘a’ and ‘a*’. The second consideration was the number of bases added to the primer at every growth step. Here, we reasoned that the fewer the number of bases added at each step, the more gradual the growth of the primer and consequently more precise the ruler. These two considerations, unfortunately, are in tension with each other. A two-letter code results in a relatively weak binding interaction between a primer and an extension hairpin. Experiments showed that a toehold binding interaction of at least 12 bases (of an {A, T} alphabet) is necessary for the extension reaction to proceed with reasonable efficiency, under standard reaction conditions. In conventional PER repeat extension reactions, the extension sequence at each step is identical to the toehold sequence (for example, ‘s’ is extended to ‘s s’). Thus, a conventional PER implementation would result in the addition of 12 bases at every step. However, we engineered a system where a 12 base toehold only adds a 4 base repeat at every step (Supplementary Fig. 2), by making the toehold sequence itself a repeat. In particular we chose ‘a’ = ‘r r r’ where ‘r’ = ‘AAAT’. Thus, a toehold sequence of a (= ‘r r r’) was extended by the sequence ‘r’ at every step of the extension reaction. Correspondingly, we chose ‘a*’ = ‘r* r* r*’ and extended it by ‘r*’ at every step. The complete sequences for the primers and extension hairpins are listed in Supplementary Table 1.

2. Inferring geometry from pairwise distance measurements

The question of inferring geometry from pairwise distance measurements is modeled as an embedding problem in two-dimensional Euclidean space. Each target point $i$ of the pattern is parameterized with X and Y co-ordinates as $(x_i, y_i)$ . We look for an embedding that minimizes the error between the experimentally measured distances and the Euclidean embedded distances. Not every measurement made by the ruler is equally reliable. We observed a clear positive correlation between the height of the major peak (in units of number of reads) and the accuracy of the measurement. Therefore, we developed an algorithm that assigns weights to the various measurements according to the height of the corresponding major peak. The embedding algorithm proceeds in three stages – pruning, producing an initial embedding and finally, refining the embedding.

**A. Pruning:** A threshold parameter (auto-tuned, as explained below) was used to filter measurements. All measurements corresponding to peak heights less than the threshold were marked unreliable and pruned. All measurements corresponding to peak heights greater than the threshold were marked equally reliable and retained. In rare cases, some points were left with very few (less than three) associated reliable distance measurements as a result of this pruning. Such points were dropped from the reconstruction by deleting all associated distance measurements.

**B. Initial embedding:** We used a robust facial reduction algorithm (*18*) to obtain an initial embedding. This initial embedding has been shown to work well as an initial solution for generating embeddings using nonlinear optimization approaches, making it less likely that the optimization process is trapped in local minima or at saddle points.

**C. Refining the embedding:** In refining the embedding, we assigned each measurement a weight in the range $[0,1]$ to denote its reliability. Measurements with corresponding peak heights less than the threshold parameter are assigned a weight of zero (i.e. pruned). Measurements with corresponding peak heights greater than the threshold parameter were assigned a positive weight $w_{ij}$ as described in the below equations. In particular the objective function $J$, which we seek to minimize, is defined as:

$$J = \sum_{i=1}^{n}\sum_{j=1}^{i-1}\frac{w_{ij}}{W}\left(d_{ij}^{measured} - d_{ij}^{embedded}\right)^2$$

where:

$\{1,2,\cdots,n\}$ are the uniquely identified points that make up the pattern,

$w_{ij} = \max\left(\frac{2}{1-e^{-k\left(p_{ij}-t\right)}} - 1, 0\right)$ is the weight assigned to the measured distance between points $i$ and $j$. The smoothness parameter $k = 0.8$, $p_{ij}$ is the peak height and $t$ is the threshold parameter. Note that $w_{ij} \in [0,1]$,

$W = \sum_{i=1}^{n} \sum_{j=1}^{i-1} w_{ij}$ is the sum of all weights, used to normalize the weights,

$d_{ij}^{measured}$ is the experimentally measured distance between points $i$ and $j$ and,

$d_{ij}^{embedded} = \sqrt{(x_i - x_j)^2 + (y_i - y_j)^2}$ is the embedded Euclidean distance.

The optimization is performed using the fminunc MATLAB function, which implements a quasi-Newton algorithm to find the minimum of an unconstrained multivariable function.

**D. Choosing the threshold parameter:** The threshold parameter was auto-set, without any *a priori* knowledge of the geometry of the designed pattern, as follows. The embedding was performed for all integer values of the threshold parameter $t$ from 0 to 100 and an embedding score was calculated as $score = J + \alpha r$

where:

$J$ is the objective function, defined above,

$\alpha$ is the penalty applied for pruning points. We empirically set $\alpha = 0.5$ for all patterns except for the Wyss pattern, for which $\alpha = 0.25$ and,

$r$ is the number of points pruned.

The optimum threshold was chosen to be the one that minimized $score$, reflecting the best agreement between the measured and embedded distances. Note that the embedding score cannot take negative values. The term $J$ captures the agreement between the measured distances and the embedded distances for all non-pruned points. The penalty term for pruned points, $\alpha r$, is necessary to prevent the algorithm from trivially achieving $J = 0$ by setting the threshold to a very high value that prunes all points because all weights $w_{ij}$ are set to zero.

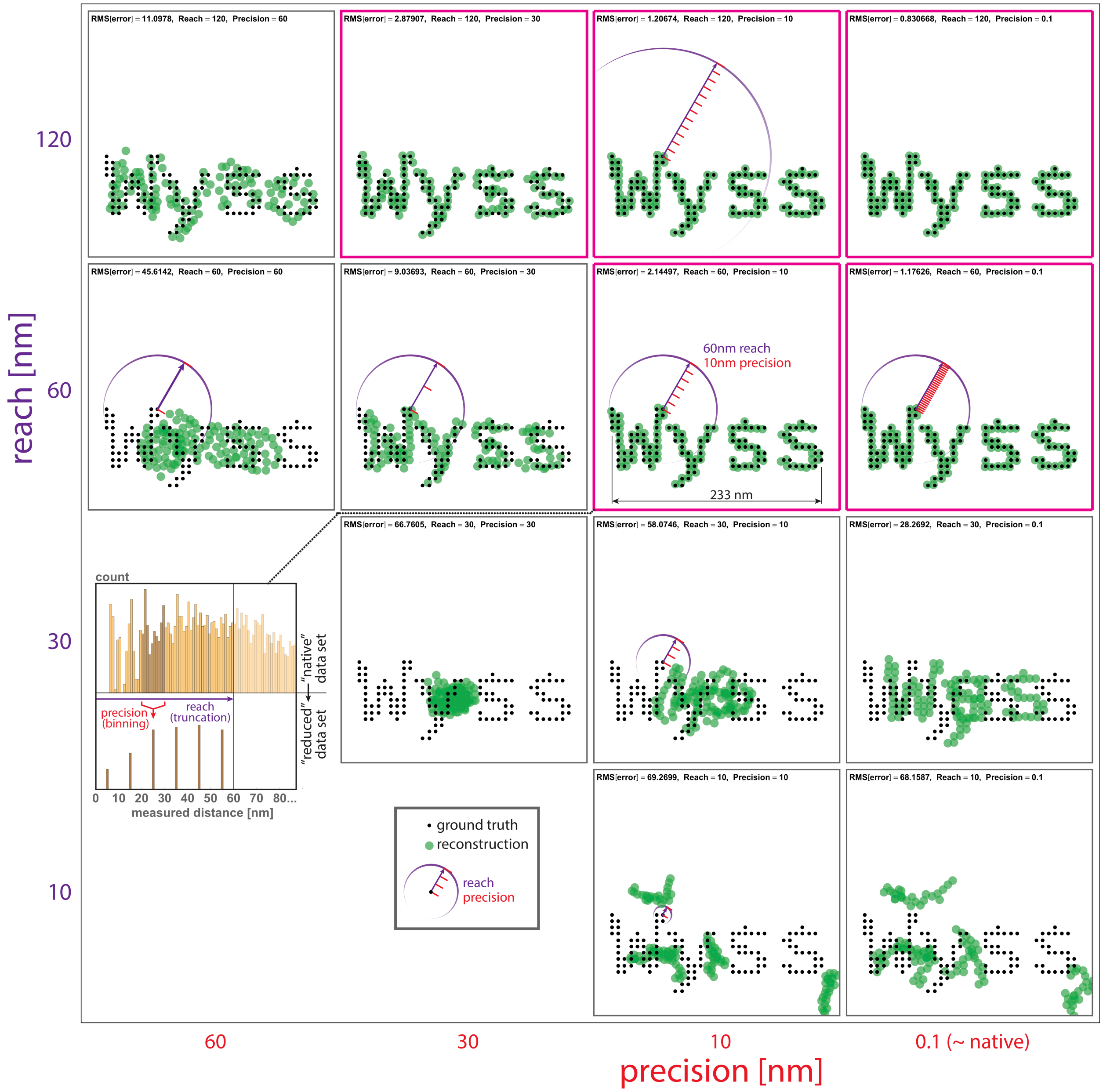

**Supplementary Fig. 1.** Effect of ruler precision and reach in reconstruction. Shown are results of a completely *in silico* study simulating diminished ruler precision and reach. A base "ground truth" geometry ("Wyss") comprising 121 points and spanning 233 nm was designed. Distance measurements for each pair (i.e., post sequencing and processing) were generated by modifying known (ground truth) values about a normal distribution of error with a fixed coefficient of variation (CV) of 5%. Measurements were then degraded by deteriorating precision and/or limiting reach. "Precision" is effectively the width of bins used to group similar measurements (histogram inset), while "reach" is the length of the farthest distance that results in a measurement. Reconstructions follow the same computational process (a reconstruction minimizing total discrepancy^2 between reads) as in the main text. Results show that limiting precision (60 nm), even at high (120 nm) reach, results in high RMSD. Similarly, high precision (0.1 nm, virtually without binning) yielded poor results if reach was limited to nearest neighbors

(10 nm). Only with reasonable precision of 10 nm and reach of 60 nm (1/4 of total pattern span, enough to cross gaps within and between letter components) did reconstructions fall near RMSD = 2 nm. Further precision or reach improvements helped, but minimally. Pink outlines denote reconstructions near or below RMSD = 2nm.

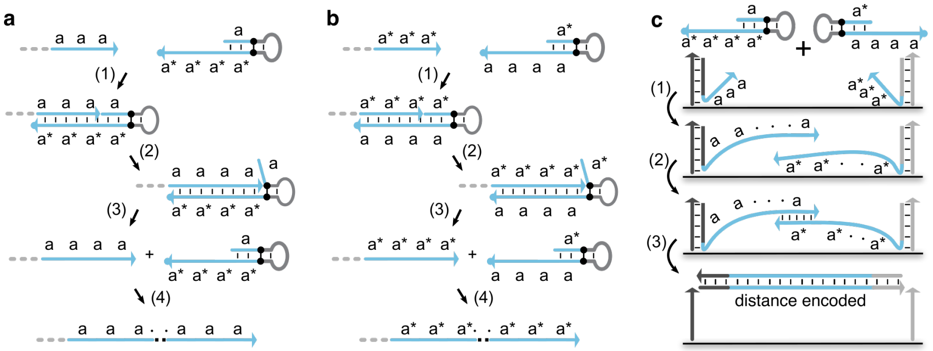

**Supplementary Fig. 2.** The full molecular ruler mechanism. Note that unlike the simplified version depicted in Fig. 2, the primer actually binds to the hairpin using a 12 base domain ‘a a a’. The rest of the details of the mechanism are identical. **a.** A primer exchange reaction (PER) cascade repeatedly adds the four base sequence domain ‘a’, as follows. (1) The recording primer hybridizes to a PER hairpin, (2) a strand displacing DNA polymerase (Bsm large fragment) extends the primer into the stem of the hairpin and in the process copies domain ‘a’. A ‘stopper’, a non-canonical base pair modification (isoG-isoC) on the template that is not recognized by the DNA polymerase, blocks further extension. The polymerase dissociates from the hairpin. (3) The recording primer is only weakly bound to the hairpin and also dissociates. (4) The above sequence of reactions repeat, adding domain ‘a’ every time. **b.** In the same manner, a complementary PER cascade repeatedly adds the four base sequence domain ‘a*’. **c.** A double-stranded DNA ‘distance record’ is generated as follows. Consider two DNA labeled targets with recording primers hybridized to them. (1) The primers take part in PER reaction cascades, as described in parts A and B, adding sequence repeats of ‘a’ and ‘a*’ respectively. (2) The extended primers hybridize, (3) copy each other with the aid of the polymerase, are displaced from the targets and released into solution, making a distance record. The whole process is isothermal and autonomous.

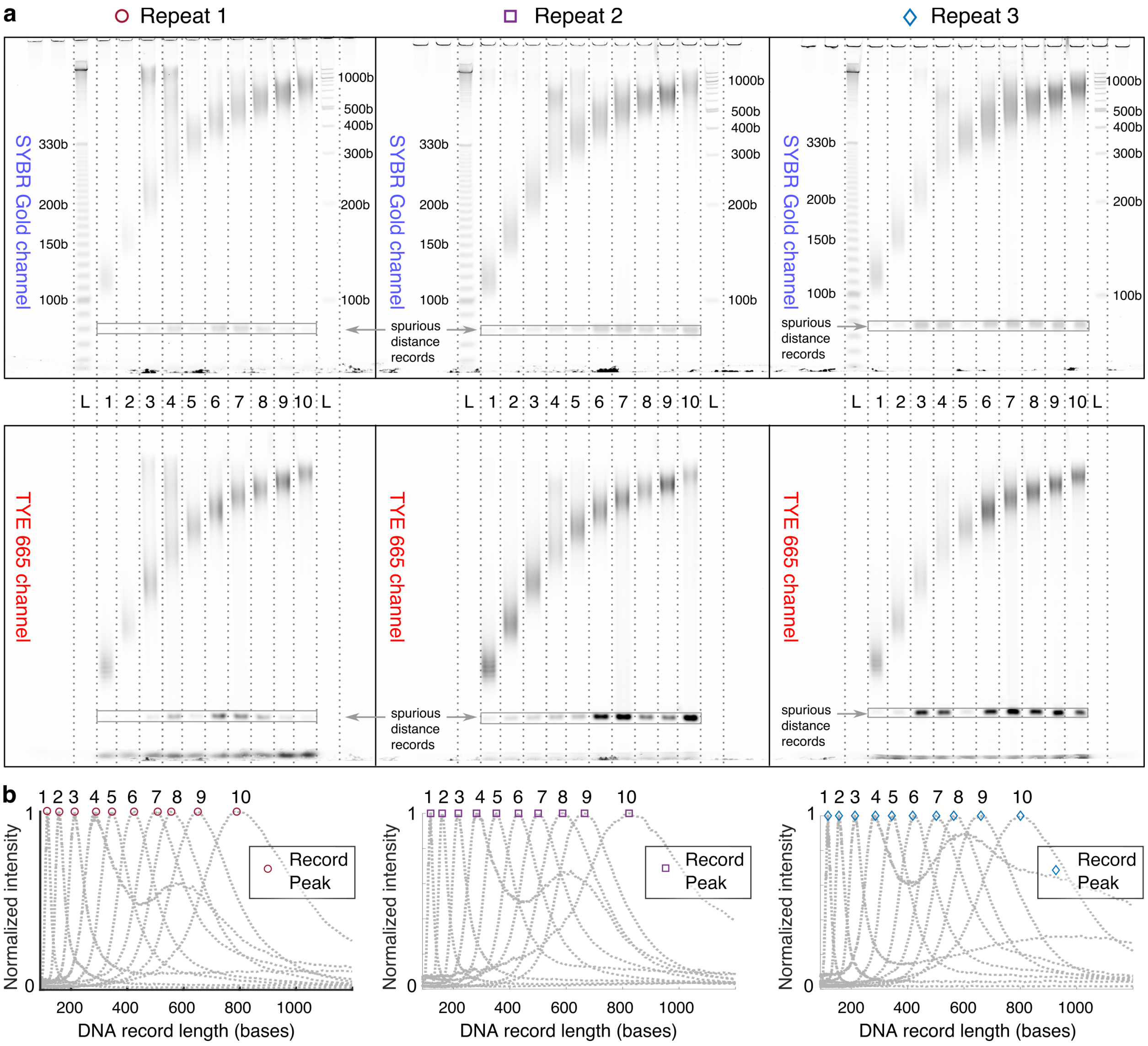

**Supplementary Fig. 3.** Molecular ruler calibration experiment. The experimental setup is described in Fig. 3. **a.** PAGE characterization of distance records. After PCR, distance records were size separated on a denaturing PAGE (see Materials and Methods section for details). The gels were imaged in two non-overlapping optical channels. Solid lines indicate gel boundaries and dotted lines are lane guides. SYBR Gold is known to stain DNA in a sequence and length dependent manner, confounding estimation of relative quantities of distance records of various lengths. Instead, the relative molar ratio of distance records of various lengths was estimated using the TYE665 channel. The TYE665 intensity should be proportional to the molar amount of the distance record, since it is introduced in an equimolar ratio by conjugation of the dye to the PCR primers. The SYBR Gold channel is used to track the ladder and hence estimate the absolute length of the distance records. **b.** Length distribution of distance records for various calibration distances. Each distance measurement reveals a distinct length distribution of distance records. The distance records are skew-normal distributed, with longer distances resulting in longer records that are more broadly distributed. The peak of the distribution was used an archetype to generate a calibration curve. Gel profiles corresponding to lane 4 show a minor peak, which we believe is due to either a defect in manufacturing DNA origami or contamination. This peak is ignored.

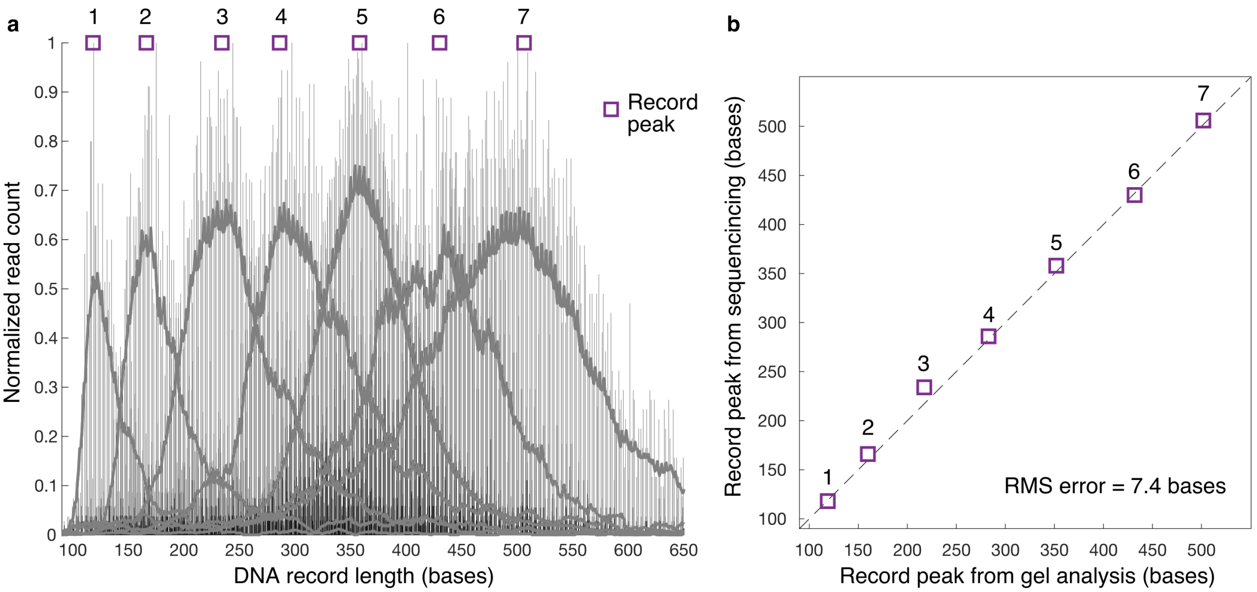

**Supplementary Fig. 4.** Characterization of the lengths of distance records by sequencing. Records from ruler experiments corresponding to seven different calibration distances were barcoded using barcoded PCR primers, pooled and then read using next-generation DNA sequencing. The programmed calibration distances are 1 = 21.4 nm, 2 = 32.0 nm, 3 = 42.8 nm, 4 = 53.4 nm, 5 = 63.9 nm, 6 = 74.8 nm and 7 = 85.3 nm **a.** The height of the bars equals the number of reads, normalized relative to the count of the most read record. The length includes primer regions, 32 bases each, flanking the variable length repeat region. The gray 'outline' curves are the moving average traces (span = 8) of the record lengths, shown here to allow the reader to discern records belonging to different calibration distances and their corresponding peaks. Next-gen sequencing sampled a few thousand reads for every pair of distances, while gel electrophoresis looks at billions of records in aggregate and hence results in smoother profiles (Fig. S3). **b.** A comparison of the peak locations characterized by gel electrophoresis (X-axis) versus peak locations obtained from next-generation DNA sequencing shows no significant systematic biases in sampling distance records with next-gen sequencing. The typical sequencing depth used in this work (about a few hundred sequences per pairwise distance) is inadequate to accurately sample record distributions corresponding to longer calibration distances (8, 9 and 10 in Fig. 3) and hence these are not included in this figure. The absence of accurate measurements of these longer distances from our sequencing data did not preclude us from successfully reconstructing patterns with points that span longer distances.

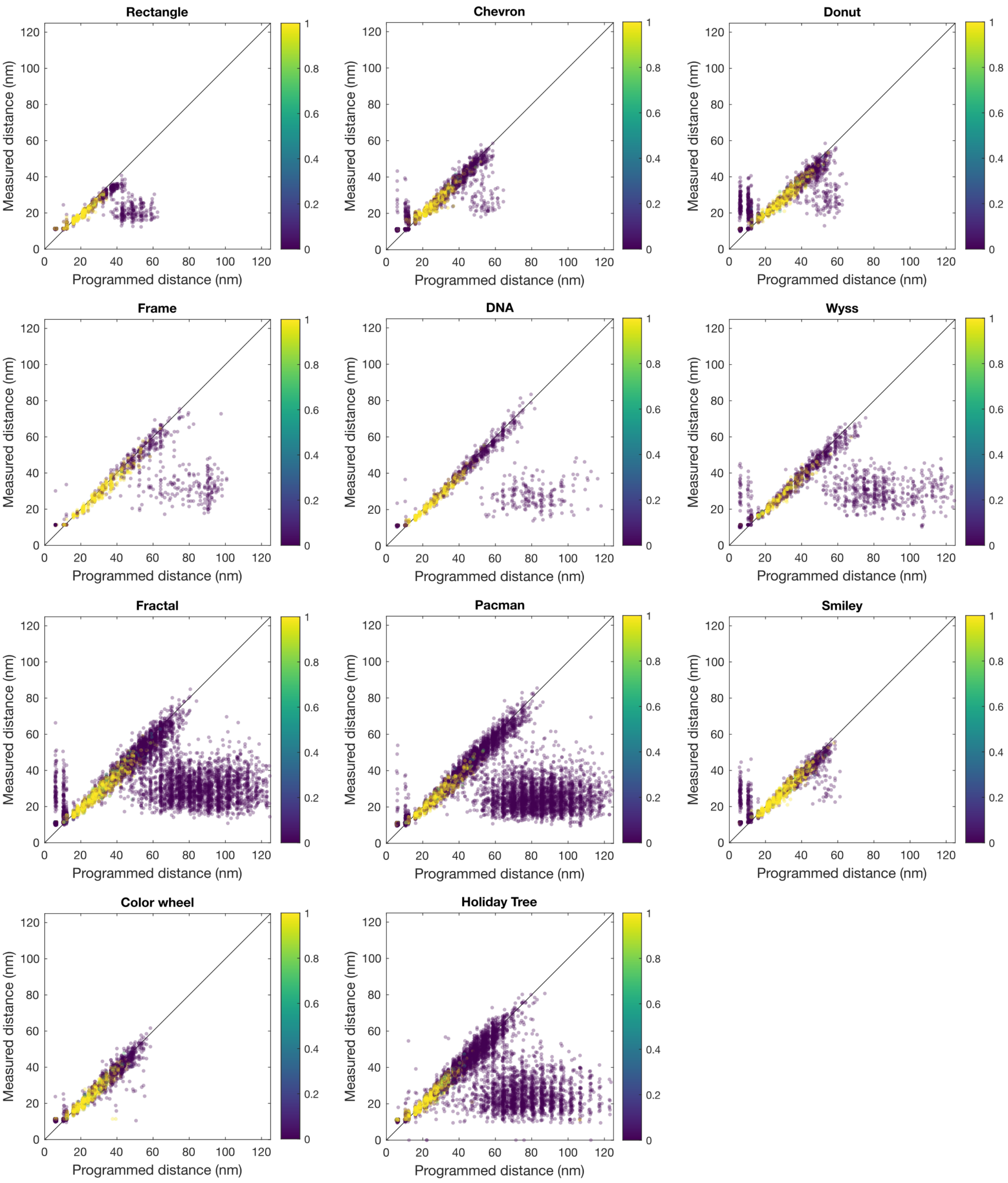

**Supplementary Fig. 5.** Programmed distances versus measured distance for each pattern from Fig. 4. Each dot within one plot corresponds to a unique pair of points. The color of the dot indicates the weight assigned to that measurement by the reconstruction algorithm (see Supplementary Note 2 for details). The overall quality of the reconstructions exceeds the average accuracy of the measurements made because of network effects. The plots indicate that points that are farther apart from each other are more likely to have an unreliable ruler measurement.

This is a result of two factors. One, fewer records are generated for longer distances because the efficiency of successfully creating records drops with increasing distance, likely limited by insufficient primer extensions due to either deleterious reactions and/or insufficient reaction time. Two, the longer the distance being measured, the more the number of sequence reads required to accurately sample the distribution of distance records. This is because the lengths of distance records are more broadly distributed as the distance being measured increases. We expect deeper sequencing to yield accurate distance measurements over longer distances.

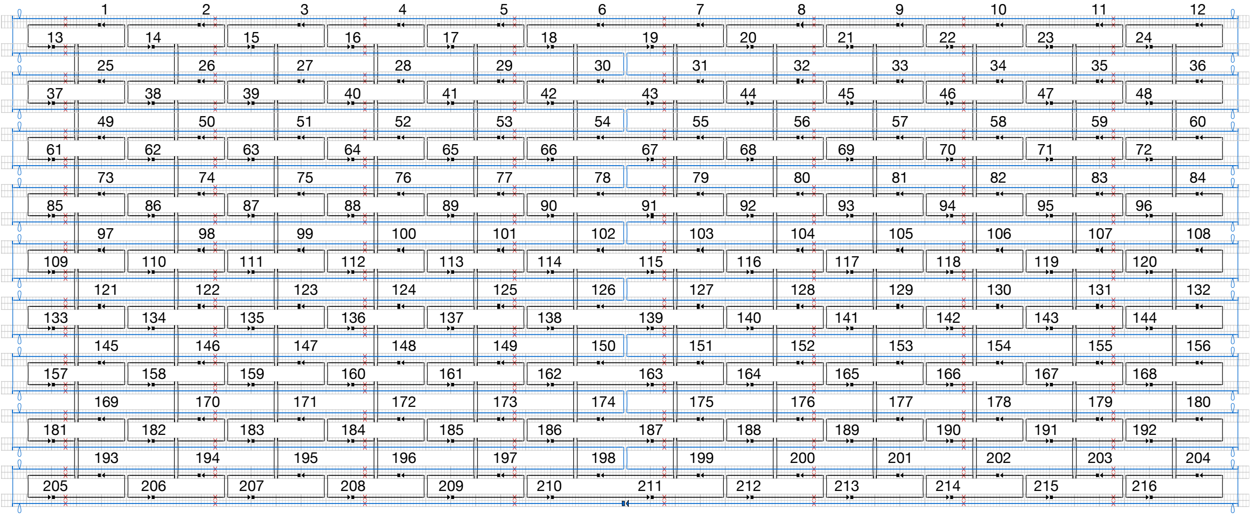

**Supplementary Fig. 6.** Scaffold and staple diagram for the DNA origami used in this work. The scaffold strand (M13mp18) is depicted in blue and the staple strands in black. Arrows at the end of strands represent 3 prime ends and the blocks at the other end represent 5 prime ends. The red crosses represent 'skipped' bases in the cadnano design software, corresponding to twist corrections (see Materials and Methods) made to minimize strain and promote flatness. The numeric labels correspond to the three prime ends of the staple strands. They are also the sequence IDs for blunt staples, extended staples and recording primers in the corresponding supplementary tables. The 'blunt' staple sequences are listed in Table S2, extended staple sequences in Table S3 and recording primer sequences in Table S4.

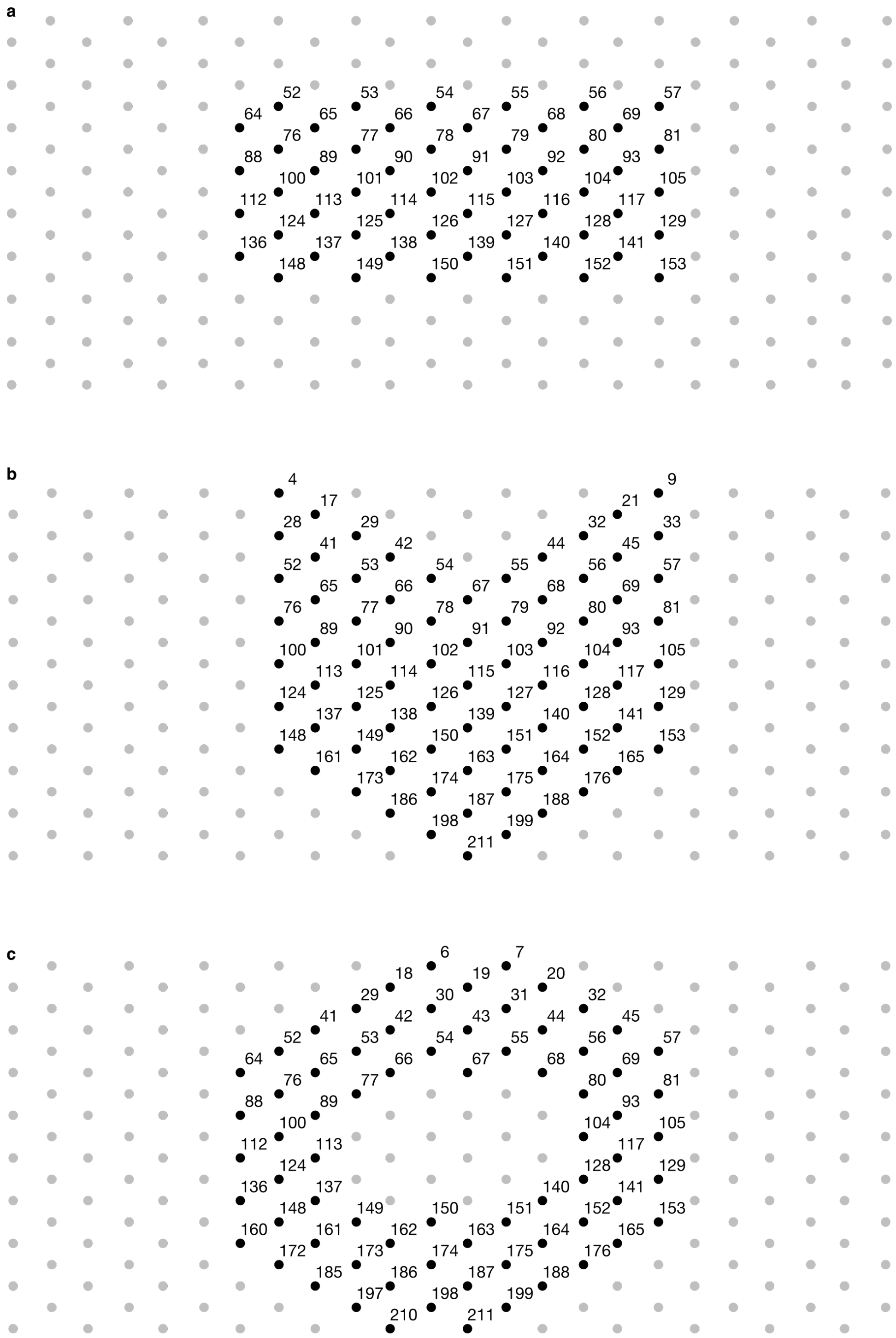

**Supplementary Fig. 7.** Pattern diagrams. Grey dots are 3' ends of blunt staples and black dots are 3' ends where either a blunt or extended staples may be incorporated. Numeric labels indicate the unique identity of every position in an origami. **a.** Rectangle. **b.** Chevron. **c.** Donut.

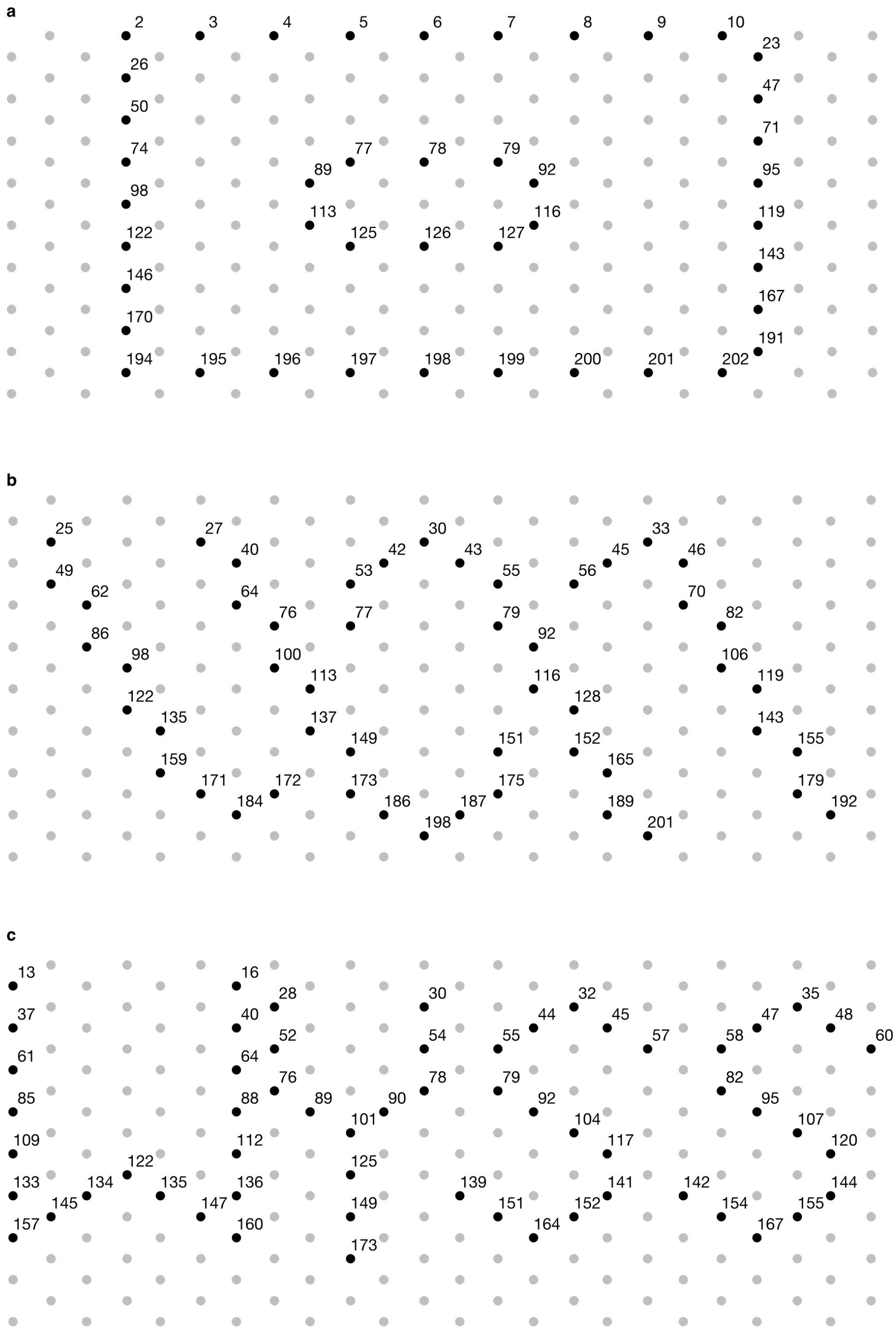

**Supplementary Figure 8.** Pattern diagrams. Grey dots are 3' ends of blunt staples and black dots are 3' ends where either a blunt or extended staples may be incorporated. Numeric labels indicate the unique identity of every position in an origami. **a.** Frame. **b.** DNA. **c.** Wyss.

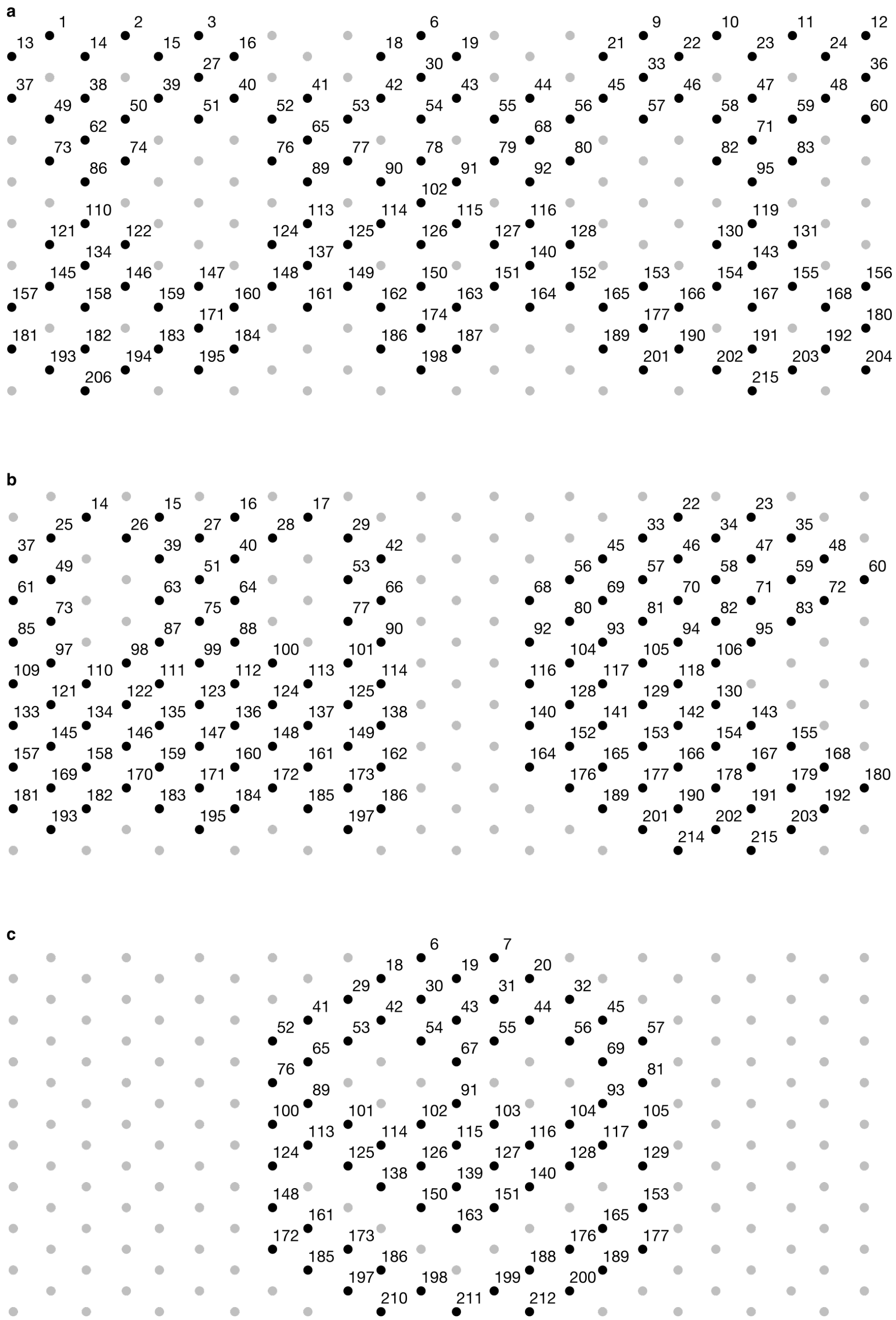

**Supplementary Figure 9.** Pattern diagrams. Grey dots are 3’ ends of blunt staples and black dots are 3’ ends where either a blunt or extended staples may be incorporated. Numeric labels indicate the unique identity of every position in an origami. **a.** Fractal. **b.** Pacman. **c.** Smiley.

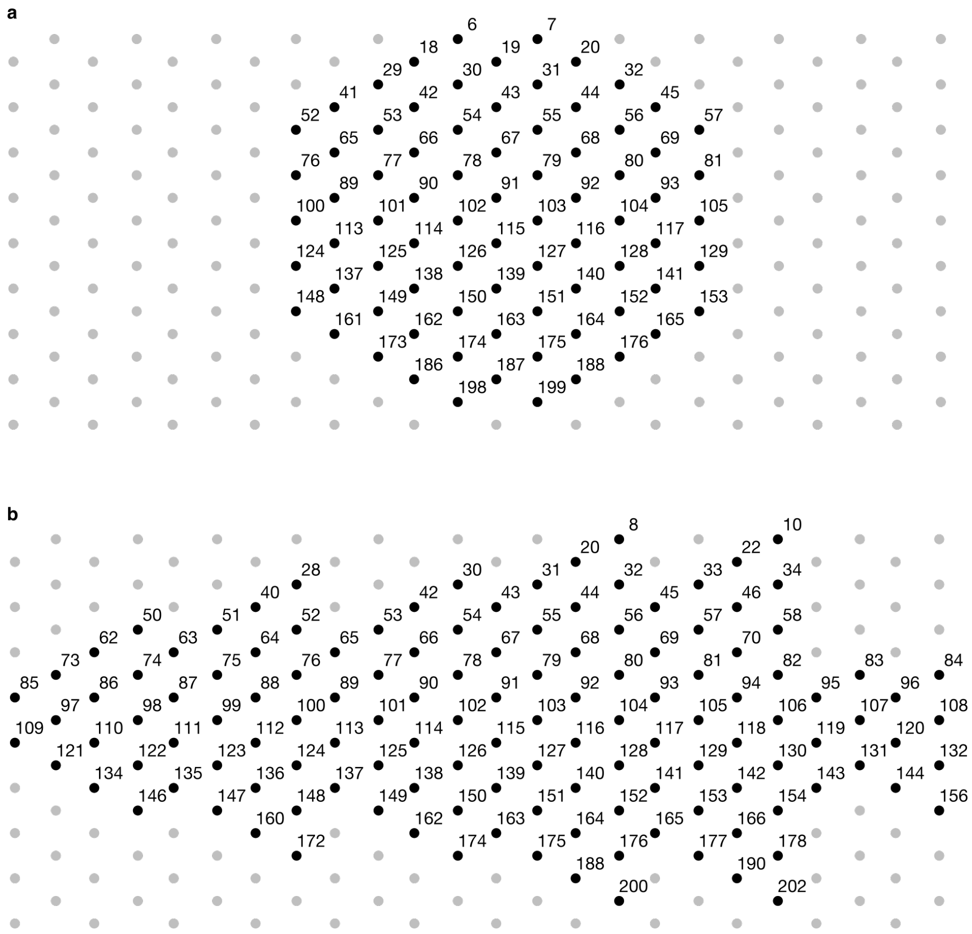

**Supplementary Figure 10.** Pattern diagrams. Grey dots are 3' ends of blunt staples and black dots are 3' ends where either a blunt or extended staples may be incorporated. Numeric labels indicate the unique identity of every position in an origami. **a.** Color wheel. **b.** Holiday tree.

| Strand | Sequence (5' to 3') |
|---|---|
| Recording primer 'a' | TCGTGCGAGTATAGAAAGTGAGGGATTAATGGAAATAAATAAAT |
| Recording primer 'a*' | TCTACCCCATGAAGAGTAAATAGGTTGTGGGAATTTATTTATTT |
| PER growth hairpin 'a' | AAAT/iisodG/mCGCCCGCCACTAGCGGGCmG/iMe-isodC/ATTTATTTATTTATTT/3InvdT/ |
| PER growth hairpin 'a*' | ATTT/iisodG/mCGCCCGCCACTAGCGGGCmG/iMe-isodC/AAATAAATAAATAAAT/3InvdT/ |
| PCR Primer 1 | /TYE665/TCGTGCGAGTATAGAAAGTGAGGGATTAATGG |
| PCR Primer 2 | /TYE665/TCTACCCCATGAAGAGTAAATAGGTTGTGGGA |

**Supplementary Table 1.** Growth hairpins, recording primers and fluorescent PCR primers for the molecular ruler calibration experiments described in Fig. 3 and Fig. S3.

| Location ID | Blunt origami staple sequences, 5’ to 3’ |
|---|---|
| 1 | AACCATCGCGTAACACTGAGTTTCGCCACCCT |
| 2 | TTGATACCACAACGCCTGTAGCACGCCACC |
| 3 | GCTTGCTTCATAGTTAGCGTAACGAGGTGTAT |
| 4 | AAAAGGAGGTCTTTCCAGACGTTACGAGAGGG |
| 5 | TTCACGTTTATGGGATTTTGCTATTTTGCT |
| 6 | CATGAAAGTATTAAGAGGCTGAGACTCCTCA |
| 7 | GCGTTTGCAGAGCCACCACCGGAATTCTGAAA |
| 8 | CATCGGCACGCCACCCTCAGAACTAATGCC |
| 9 | TTGCCTTTACCACCCTCAGAGCCGGTGCCTTG |
| 10 | AGCACCGTCCAGAGCCGCCGCCAGAGTGTACT |
| 11 | CCGGAAACAGGCAGGTCAGACGAAAGCGTC |
| 12 | ATCACCAGCAAACAAATAAATCCTGAAAGCGC |
| 13 | CAGAGCCACCACCCTCATTTTCATAGGAACC |
| 14 | CTCAGAACCGCCACCCTCAGAACCGTCACCAG |
| 15 | CACCGTACTCAGGAGGTTTAGTACTTCCACAG |
| 16 | TTGATATAAGTATAGCCCGGAATATCTAAAG |
| 17 | CAGTACCAGGCGGATAAGTGCCGTGTAAATGA |
| 18 | AGAGAAGGATTAGGATTAGCGGGGAACAACTT |
| 19 | AACAACTATTCAGCGGAGTGAGAAAAAATCAC |
| 20 | CCCTGCCTATTTCGGAACCTATTACCGCCTCC |
| 21 | AGTAACAGTGCCCGTATAAACAGTCGCCACCC |
| 22 | GGTAATAAGTTTTAACGGGGTCACCACCAGA |
| 23 | ATACATGGCTTTTGATGATACAGGCATTGACA |
| 24 | AGTCTCTGAATTTACCGTTCCAGTTTGGCCTT |
| 25 | ACCGAACTAAAGGCCGCTTTTGCGATGACAAC |
| 26 | GGAACGAGCAGCGAAAGACAGCAAAACAGC |
| 27 | AAATCCGCAACGGCTACAGAGGCTGTTTATCA |
| 28 | GATTTGTATTTTTCATGAGGAAGTAAGGCTCC |
| 29 | ATTATACCAAATACGTAATGCCATAATTTT |
| 30 | CGGAACCCATCTTTTCATAATCTAGAAAGG |
| 31 | ACGTAGAAAGAAACGCAAAGACACCCTTATTA |
| 32 | TTAAGACTTTTGTCACAATCAATGCGTTTT |
| 33 | AATAACGGTTACCAGCGCCAAAGAAATCAAGT |
| 34 | AGTTACCACAACCGATTGAGGGAGTCGATAGC |
| 35 | TTTTTAAGCGGAAATTATTCATTAGCAAGG |
| 36 | TGAAATAGCGTCACCGACTTGAGCCCAGCAAA |
| 37 | CATGTACCCCACGCATAACCGAGGCTTGCA |
| 38 | TACAAACTGATAGTTGCGCCGACAGGATCGTC |

| Location ID | Blunt origami staple sequences, 5' to 3' |
|---|---|
| 39 | ACAGCCCTTCGAGGTGAATTTCTTTCGGAACG |
| 40 | TTTTGTCCCTTTAATTGTATCGTTGAGGAC |
| 41 | ATTTTCTGGAAAATCTCCAAAAAATTCCATTA |
| 42 | TCAACAGTAAGGAATTGCGAATAACTACGAAG |
| 43 | ACACTAAACTAAAACGAAAGAGGCGTGGCAAC |
| 44 | CTCAGAGCTTTTCGGTCATAGCCCCACGGAAT |
| 45 | TCAGAGCCAGCGTCAGACTGTAGCAGAAAATT |
| 46 | ACCACCAAATCAGTAGCGACAGCAAAAGGG |
| 47 | GGAGGTTGGTCACCAATGAAACCAGGAAGGTA |
| 48 | GATATTCATAGCACCATTACCATTAAAGGTGA |
| 49 | CCAAAAGGATAGGCTGGCTGACCTATAAGGGA |
| 50 | ATACCACATGACAAGAACCGGATCTTAGCC |
| 51 | GGTAGAAAAACGTAACAAAGCTGCTTGTGTCG |
| 52 | TAAAACGAGGCTTGCCCTGACGAGACAACGGA |
| 53 | GTCAGGACAGTAAATTGGGCTTGCCCAGCG |
| 54 | ATATAAAAATACATACATAAAGAAAAGAAT |
| 55 | TACAAAATTAAGAAACGATTTTTTGTTAGCAA |
| 56 | ACGAGCGTGAAAATAGCAGCCTTGGCATGA |
| 57 | TACAATTTTAAAAACAGGGAAGCGAACGCAAT |
| 58 | AATCAAGATTAACTGAACACCCTGCCGAACAA |
| 59 | CCTCCCGAAATTGAGCGCTAATAGAAGCCC |
| 60 | GTATTCTACAAGAATTGAGTTAAGAGAAACAA |
| 61 | GGGAGTTGACCAACTTTGAAAGGTACAGAC |
| 62 | ACCCTCAGGCGCAGACGGTCAATCTCATCAAG |
| 63 | AGGGTAGCGACCTGCTCCATGTTAATTCATTA |
| 64 | TAAAGACTCATCGCCTGATAAATCATTCAG |
| 65 | AACGGGTAAAGCGCGAAACAAAGTAAACACCA |
| 66 | GCACCAACACACTCATCTTTGACCAGATGGTT |
| 67 | ATGCGATTACTTTAATCATTGTGATTATCCCA |
| 68 | AAGTTTATCCTTATTACGCAGTATGTTTAACG |
| 69 | CATATGGTAATACCCAAAAGAACTTACAGAGA |
| 70 | CGACATTGAAGGAAACCGAGGACATTAGAC |
| 71 | AATATTGAAAAAGTAAGCAGATAGAACAAAGT |
| 72 | ATTATCACCAATAGCTATCTTACCTCAGAGAG |
| 73 | CCTTTAATCAGACGACGATAAAAAACATAACG |
| 74 | AGCAAACTCTTTTGCAAAAGAAGTTTAGGA |
| 75 | TAATTCGAAATAGTAAAATGTTTATTATTACA |
| 76 | GGAAGCCCAATACTGCGGAATCGTTACGTTAA |

| Location ID | Blunt origami staple sequences, 5' to 3' |
|---|---|
| 77 | AAGCAAAGATCCCCCTCAAATGCTATACCA |
| 78 | ATCCAAAAAACAGCCATATTATATTACCTT |
| 79 | ACAATAAAGTTTATCAACAATAGATTGCCAGT |
| 80 | ACAAAAGGAAAATAATATCCCATAACGCTA |
| 81 | CGAGCCAGGTAGAAACCAATCAATCACCCAGC |
| 82 | CGCCAACACTTATCATTCCAAGAAAAGCCTTA |
| 83 | AGGGCTTATACCGCACTCATCGATAGCGAA |
| 84 | TCTTACCATTTATTTTCATCGTAGCTTATCCG |
| 85 | CAGGCGCAATTACGAGGCATAGATAACCCT |
| 86 | AGTAATCTTTCAACTAATGCAGATCCAAAATA |
| 87 | CCCAAATCGATTCATCAGTTGAGATTTTGCCA |
| 88 | TGAATAAACTAACGGAACAACAGACTGGAT |
| 89 | GAACGAGTGTTGGGAAGAAAAATCCATAAATA |
| 90 | TAATTTCATTAAGAACTGGCTCATTTTAAACA |
| 91 | ATCAGGTCAACGAGAATGACCATAATGCAGAA |
| 92 | TCAAAAATCTTTCCAGAGCCTAATTAAGTCCT |
| 93 | GAATAACATATCCTGAATCTTACCCCTAATTT |
| 94 | GGGAGAATTAGTTGCTATTTTGAATCGGCT |
| 95 | CAGAGGGTCTTGCGGGAGGTTTTGCGGGTATT |
| 96 | ATAACCCAAGAACGCGAGGCGTTTGAACAAGC |
| 97 | TTTGCGGGATTGCTGAATATAATGAGAGAGTA |
| 98 | TTGTACCATTTAAATATGCAACTGACCGGA |
| 99 | AATAAAGCAAGTTTCATTCCATATTCGCGTTT |
| 100 | CAGGCAAGATTCTGCGAACGAGTAGATTAAGA |
| 101 | ATAGTAGTTAGATACATTTCGCATAGTCAG |
| 102 | CGCGCCTCAACATGTTCAGCTAAATCAAAA |
| 103 | ACCTCCGGTGCTGATGCAAATCCAAGACGACG |
| 104 | CAAAATCACGCGAGAAAACTTTTAGTACCG |
| 105 | AGACGCTGTAATTTCATCTTCTGAGGCATTTT |
| 106 | ATCCTTGATTGAAATACCGACCGTTTTAACAA |
| 107 | AAATCGTCTAAATAAGAATAAACCAACAGT |
| 108 | AATATATGCTAGAAAAAGCCTGTTATACAAAT |
| 109 | CGTTTACTGCTCCTTTTGATAAATGGCTTA |
| 110 | GCGAGAGGCCAACAGGTCAGGATTCTGTAGCT |
| 111 | GAGGGGGTGCTTCAAAGCGAACCAAAAGTACG |
| 112 | AGCGTCCGAAAGACTTCAAATAAACAGTTG |
| 113 | TTCATTGACGGATTGCATCAAAAAGATTTAGT |
| 114 | GTTCAGAATTTACCCTGACTATTAAATGGTCA |

| Location ID | Blunt origami staple sequences, 5' to 3' |
|---|---|
| 115 | GCGAGCTGTTTAGCTATATTTTCAATAACTAT |
| 116 | GAACAAGATAAAGTAATTCTGTCCATCGCAAG |
| 117 | ACGAGCATTAATAAGAGAATATAATCAAATAT |
| 118 | GTCTTTCTGTAATTTAGGCAGACCTAAATT |
| 119 | AAACCAAGATTGAGAATCGCCATAGTGATAAA |
| 120 | AAGCCGTTGTATAAAGCCAACGCTACCGGAAT |
| 121 | TTTTTGTTTATATTTTAAATGCAAGTAATACT |
| 122 | AACGTTAAAGGTAAAGATTCAAAAAATCGG |
| 123 | GAAGATTGGAGACAGTCAAATCACAATTAAGC |
| 124 | CGGTTGATCAACCGTTCTAGCTGAAAATCATA |
| 125 | TCGTAAAAAGGGTAGCTATTTTTTCTACTA |
| 126 | ATGTAAACTTAGGTTGGGTTATTTTGGGGC |
| 127 | TTCAGGTTCTTTTACATCGGGAGACCTTTTTA |
| 128 | CGTAAAACCCTGATTGCTTTGAAAATTTAT |
| 129 | GTTAGAACCGCGCAGAGGCGAATTTTAGATTA |
| 130 | TGTTTGGACTGAGCAAAAGAAGATCCCTTAGA |
| 131 | AGATGATGAGAAAACAAAATTAACTTCTGT |
| 132 | GAGCGGAACATTTGAATTACCTTTCATAAATC |
| 133 | GAGCTTAAGAAGCCTTTATTTCTTTTTAGA |
| 134 | CAACATGTAAAACATTATGACCCTTGCCTGAG |
| 135 | GTGTCTGGCTCAGAGCATAAAGCTAGGGTGAG |
| 136 | ATTCCCAGCAAAGAATTAGCAACATCAATA |
| 137 | TTGACCATAGCATTAACATCCAATTAAATTAA |
| 138 | ATAACCTGAAAAGGTGGCATCAATGAGAGATC |
| 139 | TGGAGCAACTATCAGGTCATTGCCATACAGTA |
| 140 | ACAAAGAATAGGTCTGAGAGACTAAACAATAA |
| 141 | ATTTTAGTAGAAGAGTCAATAGTGTACCAAGT |
| 142 | TAATGGTAAACATAGCGATAGCATTCATTT |
| 143 | TAAGGCGTGCTATTAATTAATTTTGATGAAAC |
| 144 | CATAATTATGAGTGAATAACCTTGTTACATTT |
| 145 | AAGGCGATCGCGTCTGGCCTTCCTGCATTAAA |
| 146 | TTACGCCAAACATTAAATGTGAGAATTGTA |
| 147 | GGAAGGGCGGATTCTCCGTGGGAAAAAAACAG |
| 148 | GCCATTCGCCGTAATGGGATAGGTATGTACCC |
| 149 | CACCGCTTGGCGCATCGTAACCGACGGTAA |
| 150 | ACAGTACTAACGTCAGATGAATTGAGAGTC |
| 151 | TGGTCAGTAAGGTTATCTAAAATACGTAGATT |
| 152 | GAACCTCACAACTAATAGATTAGATTTGCA |

| Location ID | Blunt origami staple sequences, 5' to 3' |
|---|---|
| 153 | GCAAATGATACATTTGAGGATTTAATGGAAGG |
| 154 | GCCTGCAAACAAACAATTCGACAAATCCTGAT |
| 155 | AAAACAGATGCCCGAACGTTATTGATTATC |
| 156 | AAAAATACGTAACATTATCATTTTACCAGAAG |
| 157 | ACCCTCAAAATCAGCTCATTTTCCATCAAA |
| 158 | TAATGTGTTATTTTGTTAAAATTCGTAGCCAG |
| 159 | AAAGGCCGTATAAGCAAATATTTACGAGTAAC |
| 160 | TGATATTAATCAGAAAAGCCCCCAAACGGC |
| 161 | TGCCGGAGCTAGCATGTCAATCATCACGTTGG |
| 162 | TACAAAGGACAAGAGAATCGATGATGCATCTG |
| 163 | CAGGAAGAAGGGGACGACGACAGTTGAAAGGA |
| 164 | CGGATTCGAGAAATAAAGAAATTGTCTTTAGG |
| 165 | TACAAAATCTACCATATCAAAATTAGCCGTCA |
| 166 | CAATTACTTATACTTCTGAATAGAAGTATT |
| 167 | AAACATCAGCAATTCATCAATATACTCGTATT |
| 168 | AACAATTTTTATCATCATATTCCTAATTTTAA |
| 169 | CTGATTGCGACGGCCAGTGCCAAGTGTGCTGC |
| 170 | GTTTTTCTTCGACTCTAGAGGATTTCGCTA |
| 171 | AGAGGCGGCGAATTCGTAATCATGAACTGTTG |
| 172 | GCTGCATTTGTGTGAAATTGTTATGGCAAAGC |
| 173 | CCCGCTTTACAACATACGAGCCGTTTCCGG |
| 174 | ATTGAGGTGGCAAATCAACAGTATCGGCCT |
| 175 | TTGCAACATGACGCTCAATCGTCTTCAATATC |
| 176 | GGTAATATATTGGCAGATTCACCCCTTGCT |
| 177 | GAGTAGAAATAAAAGGGACATTCTAGCCAGCA |
| 178 | CTTCTTTGAACCCTTCTGACCTGATTAACACC |
| 179 | GTCCATCAGGCACAGACAATATTAGAAGAT |
| 180 | TAATCAGTTCTTTAATGCGCGAACTCGCCATT |
| 181 | AATAATTTAAGTTGGGTAACGCACGACGTT |
| 182 | CTTTCATCGCTGGCGAAAGGGGGACTTGCATG |
| 183 | AACCCGTCGATCGGTGCGGGCCTCCCCCGGGT |
| 184 | GGATTGACCATTCAGGCTGCGCGTCATAGC |
| 185 | TGTAGATGCTGGTGCCGGAAACCACCGCTCAC |
| 186 | CCAGTTTGTCGCACTCCAGCCAGCGAAGCATA |
| 187 | CTAACTCAAGCCTGGGGTGCCTAAGAAATACC |
| 188 | AGCACTAAAATATCAAACCCTCAAGAAATGGA |
| 189 | ATAGATAAAAAATCTAAAGCATCAAGTCACAC |
| 190 | AGACTTTCAGTGCCACGCTGAGGGCCAACA |

| Location ID | Blunt origami staple sequences, 5’ to 3’ |
|---|---|
| 191 | AAATCCTTGGTGAGGCGGTCAGTAAAGCGTAA |
| 192 | AAGTTTGACGAACGAACCACCAGCTTTGAATG |
| 193 | CCACGCTGGTTTGCCCCAGCAGGGGCAACAG |
| 194 | CTGTTTGATGGTGGTTCCGAAATCAGGGTG |
| 195 | TCCCTTATAAATCAAAAGAATAGCGCGCGGGG |
| 196 | GGGTTGAGTGTTGTTCCAGTTTGTCGTGCCA |
| 197 | GTCCACTATTAAAGAACGTGGACCTCACTG |
| 198 | TACATTTGGAAAAACGCTCATGTGAGTGAG |
| 199 | CGCTGGCAAGTGTAGCGGTCACGGCCAGCCA |
| 200 | AACCACCACACCCGCCGCGCTTACCTTGCT |
| 201 | CTACAGGGCGCGTACTATGGTTGCACTTGCCT |
| 202 | GCACGTATAACGTGCTTTCCTCGTAGCAATA |
| 203 | AGAGCGGGAGCTAAACAGGAGGCAGAGTCT |
| 204 | GGGATTTTAGACAGGAACGGTACGTGTTTTTA |
| 205 | GTAAAACCCTTCACCGCCTGGCCAAGCGGT |
| 206 | CCTGCAGGTTTCACCAGTGAGACGCGAAAATC |
| 207 | ACCGAGCTTTTGCGTATTGGGCGCCGGCAAAA |
| 208 | TGTTTCCAATGAATCGGCCAACCCGAGATA |
| 209 | AATTCCACCCAGTCGGGAAACCTGGAACAAGA |
| 210 | AAGTGTAACATTAATTGCGTTGCGTCCAACGT |
| 211 | CAAAGGGCGAAAAACCGTCTATCACGCTAGGG |
| 212 | TTATTTACCCAGAACAATATTACCCTGCGCGT |
| 213 | GACCAGTAGAACTCAAACTATCGGATGCGCCG |
| 214 | GAGATAGATTAGTAATAACATCTTTGACGA |
| 215 | GAATACGTCGCAAATTAACCGTTGTTAGAATC |
| 216 | GCTATTAGGAGGCCACCGAGTAAACGATTAAA |

**Supplementary Table 2.** Blunt staple sequences for DNA origami used in all the experiments.

| Location ID | Extended staple sequence with barcoded handle, 5' to 3' |
|---|---|
| 1 | AACCATCGCGTAACACTGAGTTTCGCCACCCTCTCCTACGCTGGTATAGACT |
| 2 | TTGATACCACAACGCCTGTAGCACGCCACCCTATTGCACTCCTCCAGTAA |
| 3 | GCTTGCTTCATAGTTAGCGTAACGAGGTGTATTTAGACCACTAGCGTTAGAC |
| 4 | AAAAGGAGGTCTTTCCAGACGTTACGAGAGGGAGCTACAAGAATGACTGACT |
| 5 | TTCACGTTTATGGGATTTTGCTATTTTGCTAGGCTTATTGACGATGGATA |
| 6 | CATGAAAGTATTAAGAGGCTGAGACTCCTCACGCAAGGTTACGTGTAATGA |
| 7 | GCGTTTGCAGAGCCACCACCGGAATTCTGAAACTCTTAGGTGGACAACTACC |
| 8 | CATCGGCACGCCACCCTCAGAACTAATGCCCTCATCTCATCCTGCATTGT |
| 9 | TTGCCTTTACCACCCTCAGAGCCGGTGCCTTGATCCAGAGGTATATCAGTCC |
| 10 | AGCACCGTCCAGAGCCGCCGCCAGAGTGTACTCTCTCTTCGTAGGTGACATC |
| 11 | CCGGAAACAGGCAGGTCAGACGAAAGCGTCCGATAGCGTTCGAACAAGAA |
| 12 | ATCACCAGCAAACAAATAAATCCTGAAAGCGCGGTACTTCTGCAGGATCATA |
| 13 | CAGAGCCACCACCCTCATTTTCATAGGAACCGGCTTGAATTCGCTTGATAG |
| 14 | CTCAGAACCGCCACCCTCAGAACCGTCACCAGGAATTAGCGTATTCCGCTTA |
| 15 | CACCGTACTCAGGAGGTTTAGTACTTCCACAGTGAGACTAACAAGATGAGGT |
| 16 | TTGATATAAGTATAGCCCGGAATATCTAAAGCCAGATATTGTTCTCCGGTT |
| 17 | CAGTACCAGGCGGATAAGTGCCGTGTAAATGACCTAGTTGCATAATGTCCTC |
| 18 | AGAGAAGGATTAGGATTAGCGGGGAACAACTTGTTACACCGTTAGAGGTTCA |
| 19 | AACAACTATTCAGCGGAGTGAGAAAAAATCACCACAATCCTATCAGTTGGTT |
| 20 | CCCTGCCTATTTCGGAACCTATTACCGCCTCCAACTAATGGCATTCAGTTCG |
| 21 | AGTAACAGTGCCCGTATAAACAGTCGCCACCCGAATAGAGACTTACGTGGCA |
| 22 | GGTAATAAGTTTTAACGGGGTCACCACCAGAAGGTGAAGTTTGTGCATAGT |
| 23 | ATACATGGCTTTTGATGATACAGGCATTGACATAGTCTCGGAGCGTATAGTG |
| 24 | AGTCTCTGAATTTACCGTTCCAGTTTGGCCTTAATGGTTCACAAGGTAGTTC |
| 25 | ACCGAACTAAAGGCCGCTTTTGCGATGACAACGAGTCTAGGTAGACCATTGT |
| 26 | GGAACGAGCAGCGAAAGACAGCAAAACAGCTTCAACCTGTTACGAAGCAA |
| 27 | AAATCCGCAACGGCTACAGAGGCTGTTTATCAGAACTCTGTCGTTCCAATTC |
| 28 | GATTTGTATTTTTCATGAGGAAGTAAGGCTCCGTAGTGCATTTTGAAGCTGC |
| 29 | ATTATACCAAATACGTAATGCCATAATTTTATAACCTCACGACTCACTAA |
| 30 | CGGAACCCATCTTTTCATAATCTAGAAAGGACAACCGGATAACAAGGATG |
| 31 | ACGTAGAAAGAAACGCAAAGACACCCTTATTAGGACCGTAAGTAACCATTCG |
| 32 | TTAAGACTTTTGTCACAATCAATGCGTTTTGTGTGACGAGTACCATCTAG |
| 33 | AATAACGGTTACCAGCGCCAAAGAAATCAAGTGTAGAGTCATTGCACGTACC |
| 34 | AGTTACCACAACCGATTGAGGGAGTCGATAGCTACGTGTTACTTCTTGCGAT |
| 35 | TTTTTAAGCGGAAATTATTCATTAGCAAGGCAGGCGGATAGTACAGTTAG |
| 36 | TGAAATAGCGTCACCGACTTGAGCCCAGCAAAAATAAGGCACTCCTCTTACT |
| 37 | CATGTACCCCACGCATAACCGAGGCTTGCAATATCCTCCAGGTCACTTAA |
| 38 | TACAAACTGATAGTTGCGCCGACAGGATCGTCTGAGACACTTTACAATCCGG |

| Location ID | Extended staple sequence with barcoded handle, 5’ to 3’ |
|---|---|
| 39 | ACAGCCCTTCGAGGTGAATTTCTTTCGGAACGAGAGGCATATGAGGTAATCG |
| 40 | TTTTGTCCCTTTAATTGTATCGTTGAGGACTAGATCACCACTAGCAACTT |
| 41 | ATTTTCTGGAAAATCTCCAAAAAATTCCATTACCTCTCGGATCAATAGGAAG |
| 42 | TCAACAGTAAGGAATTGCGAATAACTACGAAGGAAGTGTGTTGGCAAGTATT |
| 43 | ACACTAAACTAAAACGAAAGAGGCGTGGCAACTTAACGGTGTGTTGATAGGT |
| 44 | CTCAGAGCTTTTCGGTCATAGCCCCACGGAATCTCATTGTCTGGACACTAGG |
| 45 | TCAGAGCCAGCGTCAGACTGTAGCAGAAAATTAATTCATCGCATCCACTGAG |
| 46 | ACCACCAAATCAGTAGCGACAGCAAAAGGGTTGGCATCTTAAGAGACTGG |
| 47 | GGAGGTTGGTCACCAATGAAACCAGGAAGGTAGGTTCCATGTTGATACTCGA |
| 48 | GATATTCATAGCACCATTACCATTAAAGGTGACGCACATAAGTCCTTATCCT |
| 49 | CCAAAAGGATAGGCTGGCTGACCTATAAGGGAGTCGTCATTAACCTGGATGA |
| 50 | ATACCACATGACAAGAACCGGATCTTAGCCTTCCACAGTAGCGATAACTA |
| 51 | GGTAGAAAAACGTAACAAAGCTGCTTGTGTCGCTTACAGCACGTTGGTGTAA |
| 52 | TAAAACGAGGCTTGCCCTGACGAGACAACGGACTTATGATCGTAGACCGTGG |
| 53 | GTCAGGACAGTAAATTGGGCTTGCCCAGCGGAGTATTCCACACGATTGTT |
| 54 | ATATAAAAATACATACATAAAGAAAAGAATTAGGAACGACTCTCTTCTCG |
| 55 | TACAAAATTAAGAAACGATTTTTTGTTAGCAACATGCAATAGAGTTGTCGAT |
| 56 | ACGAGCGTGAAAATAGCAGCCTTGGCATGAACTACCTTGTGTAATTGGCT |
| 57 | TACAATTTTAAAAACAGGGAAGCGAACGCAATCTAATGTCGACAACGACGAC |
| 58 | AATCAAGATTAACTGAACACCCTGCCGAACAATCCTAACCTACTCCTAGTCG |
| 59 | CCTCCCGAAATTGAGCGCTAATAGAAGCCCTGAGTCGATTTGCGATTCAA |
| 60 | GTATTCTACAAGAATTGAGTTAAGAGAAACAAACGAATACCATACTGGTTGT |
| 61 | GGGAGTTGACCAACTTTGAAAGGTACAGACTGTACTATGCCTTGAATCCA |
| 62 | ACCCTCAGGCGCAGACGGTCAATCTCATCAAGGGTAACTAAGCCGTGAGATG |
| 63 | AGGGTAGCGACCTGCTCCATGTTAATTCATTATTCCTTGGCTCATTCCTAAC |
| 64 | TAAAGACTCATCGCCTGATAAATCATTCAGGCTCTATCTTACATCCGACG |
| 65 | AACGGGTAAAGCGCGAAACAAAGTAAACACCACACAGTTCACGTTATTGGTG |
| 66 | GCACCAACACACTCATCTTTGACCAGATGGTTGAGAGAATGTTCTGAACGTG |
| 67 | ATGCGATTACTTTAATCATTGTGATTATCCCAGGTTCACAATAGAGCGACTA |
| 68 | AAGTTTATCCTTATTACGCAGTATGTTTAACGTGTCGAGGTATCTCAACAAG |
| 69 | CATATGGTAATACCCAAAAGAACTTACAGAGAGCTTGTTGAACATACCAGAA |
| 70 | CGACATTGAAGGAAACCGAGGACATTAGACCCTCTGAACAAACTGAGCTT |
| 71 | AATATTGAAAAAGTAAGCAGATAGAACAAAGTAACAGGTAGGTAATAACCGG |
| 72 | ATTATCACCAATAGCTATCTTACCTCAGAGAGGGCAATGACTCAATAAGTCG |
| 73 | CCTTTAATCAGACGACGATAAAAAACATAACGCGGTGTTCAATAGACGTATC |
| 74 | AGCAAACTCTTTTGCAAAAGAAGTTTAGGAGGACTATTCGGTACTCAGAT |
| 75 | TAATTCGAAATAGTAAAATGTTTATTATTACATCAACTACGTCCATCAACAC |
| 76 | GGAAGCCCAATACTGCGGAATCGTTACGTTAAGAACTGACAATCACTCTGTT |

| Location ID | Extended staple sequence with barcoded handle, 5’ to 3’ |
|---|---|
| 77 | AAGCAAAGATCCCCCTCAAATGCTATACCAGAGAGCAACCCCTCATATAG |
| 78 | ATCCAAAAAACAGCCATATTATATTACCTTGCATAAGCAGGGTTCTAGTG |
| 79 | ACAATAAAGTTTATCAACAATAGATTGCCAGTGAAGACCATCTAGAACCTGA |
| 80 | ACAAAAGGAAAATAATATCCCATAACGCTAGAACTAGGCGATAGTCTTGC |
| 81 | CGAGCCAGGTAGAAACCAATCAATCACCCAGCTGGCTAATGTAATCACCACT |
| 82 | CGCCAACACTTATCATTCCAAGAAAAGCCTTATAACAGCTTCGTTCAATCCT |
| 83 | AGGGCTTATACCGCACTCATCGATAGCGAATATCGTACTCGCAACACAAT |
| 84 | TCTTACCATTTATTTTCATCGTAGCTTATCCGCAGCGAGTGTCTATTCGTAC |
| 85 | CAGGCGCAATTACGAGGCATAGATAACCCTTCAATCCGAACCAATGTTCT |
| 86 | AGTAATCTTTCAACTAATGCAGATCCAAAATACACAACGTAAAGCGTACAAC |
| 87 | CCCAAATCGATTCATCAGTTGAGATTTTGCCAATTCCGATCTAATCGTGCTA |
| 88 | TGAATAAACTAACGGAACAACAGACTGGATTCAGGTACTATGAGCTTGAG |
| 89 | GAACGAGTGTTGGGAAGAAAAATCCATAAATACACGTAAGTCGTCTTACATG |
| 90 | TAATTTCATTAAGAACTGGCTCATTTTAAACAGGAGACTTATCCATTGCGAA |
| 91 | ATCAGGTCAACGAGAATGACCATAATGCAGAATGAGTGGTGTGTATTGTGTC |
| 92 | TCAAAAATCTTTCCAGAGCCTAATTAAGTCCTAGGCGTACTTTTCTCGATAG |
| 93 | GAATAACATATCCTGAATCTTACCCCTAATTTTGATTGAAGGTAACCAGTGA |
| 94 | GGGAGAATTAGTTGCTATTTTGAATCGGCTTAGTGAATCCACATGCAACT |
| 95 | CAGAGGGTCTTGCGGGAGGTTTTGCGGGTATTTGTACTGGAGCAATCTAGTG |
| 96 | ATAACCCAAGAACGCGAGGCGTTTGAACAAGCCTCAAGCTATCCACATAACC |
| 97 | TTTGCGGGATTGCTGAATATAATGAGAGAGTAACTTACGTCTTTCGAGACAA |
| 98 | TTGTACCATTTAAATATGCAACTGACCGGACTCAATCATCAATTGGTGGT |
| 99 | AATAAAGCAAGTTTCATTCCATATTCGCGTTTTTACTTGGACGGAGTGTAAT |
| 100 | CAGGCAAGATTCTGCGAACGAGTAGATTAAGAGCAATCTTAGACAGTACCGT |
| 101 | ATAGTAGTTAGATACATTTCGCATAGTCAGCCATCAGAATTGTGAGTTCT |
| 102 | CGCGCCTCAACATGTTCAGCTAAATCAAAATAGTCCTCATGGTGCTTATA |
| 103 | ACCTCCGGTGCTGATGCAAATCCAAGACGACGTGGAGGTATAGGACGTAGTG |
| 104 | CAAAATCACGCGAGAAAACTTTTAGTACCGCTTCGGCTTCTAGTGTAAGC |
| 105 | AGACGCTGTAATTTCATCTTCTGAGGCATTTTTGGTACAATGCTCTCAATGT |
| 106 | ATCCTTGATTGAAATACCGACCGTTTTAACAACTATAACGCACTACGAGGTG |
| 107 | AAATCGTCTAAATAAGAATAAACCAACAGTGATGATTCGTTGCTACATCG |
| 108 | AATATATGCTAGAAAAAGCCTGTTATACAAATTGCGAACTTATTACACCTCG |
| 109 | CGTTTACTGCTCCTTTTGATAAATGGCTTACCGATAACCGTCTTGATGAC |
| 110 | GCGAGAGGCCAACAGGTCAGGATTCTGTAGCTCCTACGTTATCTGCAAGGAT |
| 111 | GAGGGGGTGCTTCAAAGCGAACCAAAAGTACGTTCATTAGCGCAGATCAGTT |
| 112 | AGCGTCCGAAAGACTTCAAATAAACAGTTGGAAGGTTACTTCCAGACCTC |
| 113 | TTCATTGACGGATTGCATCAAAAAGATTTAGTTATGTCAGTGGATACTTGGT |
| 114 | GTTCAGAATTTACCCTGACTATTAAATGGTCATTAACACGCTTTAGGTTCGT |

| Location ID | Extended staple sequence with barcoded handle, 5’ to 3’ |
|---|---|
| 115 | GCGAGCTGTTTAGCTATATTTTCAATAACTATCTCGGAAGTTCTCGTAGTAT |
| 116 | GAACAAGATAAAGTAATTCTGTCCATCGCAAGAGTCAGAACATTCGATCCAC |
| 117 | ACGAGCATTAATAAGAGAATATAATCAAATATCGAGTCACATCAAGGACATT |
| 118 | GTCTTTCTGTAATTTAGGCAGACCTAAATTGTACAAGTTCGAACCTACCG |
| 119 | AAACCAAGATTGAGAATCGCCATAGTGATAAAGCTACGATACTGCTGAACTC |
| 120 | AAGCCGTTGTATAAAGCCAACGCTACCGGAATCGACACGTATAGTCTGTCTA |
| 121 | TTTTTGTTTATATTTTAAATGCAAGTAATACTGACATCCGTTTTGAGGTCTT |
| 122 | AACGTTAAAGGTAAAGATTCAAAAAATCGGAGCGTATTAGGGATCTTACT |
| 123 | GAAGATTGGAGACAGTCAAATCACAATTAAGCTCTAAGCGTACTTGGCTAGA |
| 124 | CGGTTGATCAACCGTTCTAGCTGAAAATCATAGACTGTTCCTGGACTGATAT |
| 125 | TCGTAAAAAGGGTAGCTATTTTTTCTACTACGGATAGAAGGAACACTACA |
| 126 | ATGTAAACTTAGGTTGGGTTATTTTGGGGCGAGCTAGTCCTAGTGGTTAG |
| 127 | TTCAGGTTCTTTTACATCGGGAGACCTTTTTATGTAGCGGATTATCAATGCG |
| 128 | CGTAAAACCCTGATTGCTTTGAAAATTTATTTAGCTGTTCGTCGATGATT |
| 129 | GTTAGAACCGCGCAGAGGCGAATTTTAGATTAACCATTCTTCGTAGTATCCT |
| 130 | TGTTTGGACTGAGCAAAAGAAGATCCCTTAGATTCAGTCTACTTCCTTACGA |
| 131 | AGATGATGAGAAAACAAAATTAACTTCTGTCTGTCACCATCTAGTACGTT |
| 132 | GAGCGGAACATTTGAATTACCTTTCATAAATCTTGTAACCTGGAATACGGTT |
| 133 | GAGCTTAAGAAGCCTTTATTTCTTTTTAGATGCGTATTCTCTCTTGTGTT |
| 134 | CAACATGTAAAACATTATGACCCTTGCCTGAGTGTATCACCGGATATACCTC |
| 135 | GTGTCTGGCTCAGAGCATAAAGCTAGGGTGAGAATGGACCGTCATTAGAAGC |
| 136 | ATTCCCAGCAAAGAATTAGCAACATCAATAGAATAGTTGGCCTGAATCTT |
| 137 | TTGACCATAGCATTAACATCCAATTAAATTAAACGCTTACTAGCGATGTTAT |
| 138 | ATAACCTGAAAAGGTGGCATCAATGAGAGATCCCGAGGTTAGGTATTAGAGG |
| 139 | TGGAGCAACTATCAGGTCATTGCCATACAGTATAAGATCCGATACCATAGCG |
| 140 | ACAAAGAATAGGTCTGAGAGACTAAACAATAAGCTGGAACAAGTCTGTTATG |
| 141 | ATTTTAGTAGAAGAGTCAATAGTGTACCAAGTCCATATAAGGAAGCAGAGGC |
| 142 | TAATGGTAAACATAGCGATAGCATTCATTTCCATGACTTACGATGAGTGG |
| 143 | TAAGGCGTGCTATTAATTAATTTTGATGAAACAGGTATTCACTGTGGTGTTC |
| 144 | CATAATTATGAGTGAATAACCTTGTTACATTTCTCAGTAGACAATCTTCGCT |
| 145 | AAGGCGATCGCGTCTGGCCTTCCTGCATTAAACATACTACAGTCCATGTGCG |
| 146 | TTACGCCAAACATTAAATGTGAGAATTGTAAACTCACTTCCGTCAATTCG |
| 147 | GGAAGGGCGGATTCTCCGTGGGAAAAAAACAGGGCTATCAATAGACTCCTCG |
| 148 | GCCATTCGCCGTAATGGGATAGGTATGTACCCTATACCTGTGTCACTCGTAG |
| 149 | CACCGCTTGGCGCATCGTAACCGACGGTAAGATCCGCTATTCTATTCCGA |
| 150 | ACAGTACTAACGTCAGATGAATTGAGAGTCTCATACGTTCCGGAGATGTA |
| 151 | TGGTCAGTAAGGTTATCTAAAATACGTAGATTGGTGAATAGGGGTGATTGAT |
| 152 | GAACCTCACAACTAATAGATTAGATTTGCAACGCTTAGATGTAACACAGA |

| Location ID | Extended staple sequence with barcoded handle, 5' to 3' |
|---|---|
| 153 | GCAAATGATACATTTGAGGATTTAATGGAAGGCTTCGGTATTTGGTCATCGA |
| 154 | GCCTGCAAACAAACAATTCGACAAATCCTGATATTGACCTACCACATGAGTA |
| 155 | AAAACAGATGCCCGAACGTTATTGATTATCCATTGTAGGCCTTGGAGAAT |
| 156 | AAAAATACGTAACATTATCATTTTACCAGAAGGCGTACCTAAACATAGGCTC |
| 157 | ACCCTCAAAATCAGCTCATTTTCCATCAAACATGTGATACCTTGACACAT |
| 158 | TAATGTGTTATTTTGTTAAAATTCGTAGCCAGTTGCATCACTTCATCAGGTA |
| 159 | AAAGGCCGTATAAGCAAATATTTACGAGTAACTTCAAGCACATGACCTTATG |
| 160 | TGATATTAATCAGAAAAGCCCCCAAACGGCCGTATTGTCATCCGTCACAG |
| 161 | TGCCGGAGCTAGCATGTCAATCATCACGTTGGTGCTACTGTTTCCTTCATGG |
| 162 | TACAAAGGACAAGAGAATCGATGATGCATCTGGAGTTACCTTTTCGAGTGTC |
| 163 | CAGGAAGAAGGGGACGACGACAGTTGAAAGGATAGGTCTGTTGTAGCAACAA |
| 164 | CGGATTCGAGAAATAAAGAAATTGTCTTTAGGTATAGAGGAGCCAACCACTA |
| 165 | TACAAAATCTACCATATCAAAATTAGCCGTCAGATGGTATTGGGAATGGATG |
| 166 | CAATTACTTATACTTCTGAATAGAAGTATTCGGTTGAGAAATACCGTCTT |
| 167 | AAACATCAGCAATTCATCAATATACTCGTATTTCACAACAGTGTTGAGCTAT |
| 168 | AACAATTTTTATCATCATATTCCTAATTTTAATCACCTAGAGTCTTGGATGT |
| 169 | CTGATTGCGACGGCCAGTGCCAAGTGTGCTGCGGTTCATATCTCGACCTCTT |
| 170 | GTTTTTCTTCGACTCTAGAGGATTTCGCTACTCATGGAATTACGTCGATC |
| 171 | AGAGGCGGCGAATTCGTAATCATGAACTGTTGATATGGTGCAGGAGTATTGT |
| 172 | GCTGCATTTGTGTGAAATTGTTATGGCAAAGCGCATTACATCATGGTCCTAG |
| 173 | CCCGCTTTACAACATACGAGCCGTTTCCGGCAACTTCGAGGAGAATCCAT |
| 174 | ATTGAGGTGGCAAATCAACAGTATCGGCCTGGTCGGTCTTATTCGACATG |
| 175 | TTGCAACATGACGCTCAATCGTCTTCAATATCGGTTGTTGTTGTACACACAC |
| 176 | GGTAATATATTGGCAGATTCACCCCTTGCTAACGTGGAGTTCCGTATATC |
| 177 | GAGTAGAAATAAAAGGGACATTCTAGCCAGCATAGGACCATGGTATCTTAGG |
| 178 | CTTCTTTGAACCCTTCTGACCTGATTAACACCGGATTGTTACTCCGAGTAGG |
| 179 | GTCCATCAGGCACAGACAATATTAGAAGATGAATCAAGCTGCAATAGTGT |
| 180 | TAATCAGTTCTTTAATGCGCGAACTCGCCATTCACAATTACGACGCATTAGG |
| 181 | AATAATTTAAGTTGGGTAACGCACGACGTTTAGAACTGCTGTTGTGTTGT |
| 182 | CTTTCATCGCTGGCGAAAGGGGGACTTGCATGCAATTCCAGTAACGGCATAG |
| 183 | AACCCGTCGATCGGTGCGGGCCTCCCCCGGGTTCTGATGTTGGACTTGTGTA |
| 184 | GGATTGACCATTCAGGCTGCGCGTCATAGCGTATCCACTTACACGGTTCT |
| 185 | TGTAGATGCTGGTGCCGGAAACCACCGCTCACAACTATCACCTTCTGTGTGA |
| 186 | CCAGTTTGTCGCACTCCAGCCAGCGAAGCATAGATAAGAGCGCCACTTATGT |
| 187 | CTAACTCAAGCCTGGGGTGCCTAAGAAATACCGGTCTACTATCATGTGGCTT |
| 188 | AGCACTAAAATATCAAACCCTCAAGAAATGGATATTGGACACTAAGCTCGTT |
| 189 | ATAGATAAAAAATCTAAAGCATCAAGTCACACCCTTCTCAATTTGCTTGTCA |
| 190 | AGACTTTCAGTGCCACGCTGAGGGCCAACATCTGTATTCCAACACTGGAG |

| Location ID | Extended staple sequence with barcoded handle, 5’ to 3’ |
|---|---|
| 191 | AAATCCTTGGTGAGGCGGTCAGTAAAGCGTAACCAACTTCTCTTAGTGTGCT |
| 192 | AAGTTTGACGAACGAACCACCAGCTTTGAATGGAGGACTAGAAAGTCGTTGT |
| 193 | CCACGCTGGTTTGCCCCAGCAGGGGCAACAGTCTCGGTGAATAAGTCACAC |
| 194 | CTGTTTGATGGTGGTTCCGAAATCAGGGTGGTGAACCTATTCGGCTGTAG |
| 195 | TCCCTTATAAATCAAAAGAATAGCGCGCGGGGAAGGATTGGTACCGGATAAT |
| 196 | GGGTTGAGTGTTGTTCCAGTTTGTCGTGCCAGTACGTGGTTCACTAACCAT |
| 197 | GTCCACTATTAAAGAACGTGGACCTCACTGTGACGGTAGAGTACATTCGT |
| 198 | TACATTTGGAAAAACGCTCATGTGAGTGAGTAGTGTCTTGTAGGTCATCC |
| 199 | CGCTGGCAAGTGTAGCGGTCACGGCCAGCCACTGACCTGTATTGTCCACTG |
| 200 | AACCACCACACCCGCCGCGCTTACCTTGCTCATCTAGCAATCTACGCACG |
| 201 | CTACAGGGCGCGTACTATGGTTGCACTTGCCTTGGATCTGAAAAGAATGTGC |
| 202 | GCACGTATAACGTGCTTTCCTCGTAGCAATACAATACGCGACTCTGCTATT |
| 203 | AGAGCGGGAGCTAAACAGGAGGCAGAGTCTTCTGAGGATCTTCCTACCAG |
| 204 | GGGATTTTAGACAGGAACGGTACGTGTTTTTAAACTCGCACATGTGTCTAAG |
| 205 | GTAAAACCCTTCACCGCCTGGCCAAGCGGTGTGACCGACAGTTACTCATG |
| 206 | CCTGCAGGTTTCACCAGTGAGACGCGAAAATCGATACATTGCGTACGACATT |
| 207 | ACCGAGCTTTTGCGTATTGGGCGCCGGCAAAAGTATGACACCGAGCAATTCT |
| 208 | TGTTTCCAATGAATCGGCCAACCCGAGATACAGCTCTAATATCGAACGGT |
| 209 | AATTCCACCCAGTCGGGAAACCTGGAACAAGAGCTATGACAACCGCAGTATA |
| 210 | AAGTGTAACATTAATTGCGTTGCGTCCAACGTGCATTGTTGTGTAGACAAGT |
| 211 | CAAAGGGCGAAAAACCGTCTATCACGCTAGGGGAACTGATTGAACAACGGTC |
| 212 | TTATTTACCCAGAACAATATTACCCTGCGCGTGTGGAACATTCAATGACAGG |
| 213 | GACCAGTAGAACTCAAACTATCGGATGCGCCGCGAATCAATGTTCAAGTGGT |
| 214 | GAGATAGATTAGTAATAACATCTTTGACGAGATCTGTTCACGAAGTCTCC |
| 215 | GAATACGTCGCAAATTAACCGTTGTTAGAATCTTGCATGGTAGATCTTCTCC |
| 216 | GCTATTAGGAGGCCACCGAGTAAACGATTAAACCAGATACAACACGTTCAAT |

**Supplementary Table 3.** Staples extended with barcoded handles for recruiting barcoded recording primers. These are used in experiments described in Fig. 4. The location IDs correspond to the numeric position labels in Fig. S6.

| Location ID | Barcoded recording primer type ‘a’ | Barcoded recording primer type ‘a*’ |
|---|---|---|
| 1 | GTCTTGAGCAAATAGCAGGTGACAAGTCTATACCAGCGTAGGAGAAATAAATAAAT | TCCATCTTGTCTGTTAGCAAGCTGAGTCTATACCAGCGTAGGAGATTTATTTATTT |
| 2 | GTCTTGAGCAAATAGCAGGTGACATTACTGGAGGAGTGCAATAGAAATAAATAAAT | TCCATCTTGTCTGTTAGCAAGCTGTTACTGGAGGAGTGCAATAGATTTATTTATTT |
| 3 | GTCTTGAGCAAATAGCAGGTGACAGTCTAACGCTAGTGGTCTAAAAATAAATAAAT | TCCATCTTGTCTGTTAGCAAGCTGGTCTAACGCTAGTGGTCTAAATTTATTTATTT |
| 4 | GTCTTGAGCAAATAGCAGGTGACAAGTCAGTCATTCTTGTAGCTAAATAAATAAAT | TCCATCTTGTCTGTTAGCAAGCTGAGTCAGTCATTCTTGTAGCTATTTATTTATTT |
| 5 | GTCTTGAGCAAATAGCAGGTGACATATCCATCGTCAATAAGCCTAAATAAATAAAT | TCCATCTTGTCTGTTAGCAAGCTGTATCCATCGTCAATAAGCCTATTTATTTATTT |
| 6 | GTCTTGAGCAAATAGCAGGTGACATCATTACACGTAACCTTGCGAAATAAATAAAT | TCCATCTTGTCTGTTAGCAAGCTGTCATTACACGTAACCTTGCGATTTATTTATTT |
| 7 | GTCTTGAGCAAATAGCAGGTGACAGGTAGTTGTCCACCTAAGAGAAATAAATAAAT | TCCATCTTGTCTGTTAGCAAGCTGGGTAGTTGTCCACCTAAGAGATTTATTTATTT |
| 8 | GTCTTGAGCAAATAGCAGGTGACAACAATGCAGGATGAGATGAGAAATAAATAAAT | TCCATCTTGTCTGTTAGCAAGCTGACAATGCAGGATGAGATGAGATTTATTTATTT |
| 9 | GTCTTGAGCAAATAGCAGGTGACAGGACTGATATACCTCTGGATAAATAAATAAAT | TCCATCTTGTCTGTTAGCAAGCTGGGACTGATATACCTCTGGATATTTATTTATTT |
| 10 | GTCTTGAGCAAATAGCAGGTGACAGATGTCACCTACGAAGAGAGAAATAAATAAAT | TCCATCTTGTCTGTTAGCAAGCTGGATGTCACCTACGAAGAGAGATTTATTTATTT |
| 11 | GTCTTGAGCAAATAGCAGGTGACATTCTTGTTCGAACGCTATCGAAATAAATAAAT | TCCATCTTGTCTGTTAGCAAGCTGTTCTTGTTCGAACGCTATCGATTTATTTATTT |
| 12 | GTCTTGAGCAAATAGCAGGTGACATATGATCCTGCAGAAGTACCAAATAAATAAAT | TCCATCTTGTCTGTTAGCAAGCTGTATGATCCTGCAGAAGTACCATTTATTTATTT |
| 13 | GTCTTGAGCAAATAGCAGGTGACACTATCAAGCGAATTCAAGCCAAATAAATAAAT | TCCATCTTGTCTGTTAGCAAGCTGCTATCAAGCGAATTCAAGCCATTTATTTATTT |
| 14 | GTCTTGAGCAAATAGCAGGTGACATAAGCGGAATACGCTAATTCAAATAAATAAAT | TCCATCTTGTCTGTTAGCAAGCTGTAAGCGGAATACGCTAATTCATTTATTTATTT |
| 15 | GTCTTGAGCAAATAGCAGGTGACAACCTCATCTTGTTAGTCTCAAAATAAATAAAT | TCCATCTTGTCTGTTAGCAAGCTGACCTCATCTTGTTAGTCTCAATTTATTTATTT |
| 16 | GTCTTGAGCAAATAGCAGGTGACAAACCGGAGAACAATATCTGGAAATAAATAAAT | TCCATCTTGTCTGTTAGCAAGCTGAACCGGAGAACAATATCTGGATTTATTTATTT |
| 17 | GTCTTGAGCAAATAGCAGGTGACAGAGGACATTATGCAACTAGGAAATAAATAAAT | TCCATCTTGTCTGTTAGCAAGCTGGAGGACATTATGCAACTAGGATTTATTTATTT |
| 18 | GTCTTGAGCAAATAGCAGGTGACATGAACCTCTAACGGTGTAACAAATAAATAAAT | TCCATCTTGTCTGTTAGCAAGCTGTGAACCTCTAACGGTGTAACATTTATTTATTT |
| 19 | GTCTTGAGCAAATAGCAGGTGACAAACCAACTGATAGGATTGTGAAATAAATAAAT | TCCATCTTGTCTGTTAGCAAGCTGAACCAACTGATAGGATTGTGATTTATTTATTT |
| 20 | GTCTTGAGCAAATAGCAGGTGACACGAACTGAATGCCATTAGTTAAATAAATAAAT | TCCATCTTGTCTGTTAGCAAGCTGCGAACTGAATGCCATTAGTTATTTATTTATTT |
| 21 | GTCTTGAGCAAATAGCAGGTGACATGCCACGTAAGTCTCTATTCAAATAAATAAAT | TCCATCTTGTCTGTTAGCAAGCTGTGCCACGTAAGTCTCTATTCATTTATTTATTT |
| 22 | GTCTTGAGCAAATAGCAGGTGACAACTATGCACAAACTTCACCTAAATAAATAAAT | TCCATCTTGTCTGTTAGCAAGCTGACTATGCACAAACTTCACCTATTTATTTATTT |
| 23 | GTCTTGAGCAAATAGCAGGTGACACACTATACGCTCCGAGACTAAAATAAATAAAT | TCCATCTTGTCTGTTAGCAAGCTGCACTATACGCTCCGAGACTAATTTATTTATTT |
| 24 | GTCTTGAGCAAATAGCAGGTGACAGAACTACCTTGTGAACCATTAAATAAATAAAT | TCCATCTTGTCTGTTAGCAAGCTGGAACTACCTTGTGAACCATTATTTATTTATTT |
| 25 | GTCTTGAGCAAATAGCAGGTGACAACAATGGTCTACCTAGACTCAAATAAATAAAT | TCCATCTTGTCTGTTAGCAAGCTGACAATGGTCTACCTAGACTCATTTATTTATTT |
| 26 | GTCTTGAGCAAATAGCAGGTGACATTGCTTCGTAACAGGTTGAAAAATAAATAAAT | TCCATCTTGTCTGTTAGCAAGCTGTTGCTTCGTAACAGGTTGAAATTTATTTATTT |
| 27 | GTCTTGAGCAAATAGCAGGTGACAGAATTGGAACGACAGAGTTCAAATAAATAAAT | TCCATCTTGTCTGTTAGCAAGCTGGAATTGGAACGACAGAGTTCATTTATTTATTT |
| 28 | GTCTTGAGCAAATAGCAGGTGACAGCAGCTTCAAAATGCACTACAAATAAATAAAT | TCCATCTTGTCTGTTAGCAAGCTGGCAGCTTCAAAATGCACTACATTTATTTATTT |
| 29 | GTCTTGAGCAAATAGCAGGTGACATTAGTGAGTCGTGAGGTTATAAATAAATAAAT | TCCATCTTGTCTGTTAGCAAGCTGTTAGTGAGTCGTGAGGTTATATTTATTTATTT |

| Location ID | Barcoded recording primer type 'a' | Barcoded recording primer type 'a*' |
|---|---|---|
| 30 | GTCTTGAGCAAATAGCAGGTGACACATC CTTGTTATCCGGTTGTAAATAAATAAAT | TCCATCTTGTCTGTTAGCAAGCTGCATC CTTGTTATCCGGTTGTATTTATTTATTT |
| 31 | GTCTTGAGCAAATAGCAGGTGACACGAA TGGTTACTTACGGTCCAAATAAATAAAT | TCCATCTTGTCTGTTAGCAAGCTGCGAA TGGTTACTTACGGTCCATTTATTTATTT |
| 32 | GTCTTGAGCAAATAGCAGGTGACACTAG ATGGTACTCGTCACACAAATAAATAAAT | TCCATCTTGTCTGTTAGCAAGCTGCTAG ATGGTACTCGTCACACATTTATTTATTT |
| 33 | GTCTTGAGCAAATAGCAGGTGACAGGTA CGTGCAATGACTCTACAAATAAATAAAT | TCCATCTTGTCTGTTAGCAAGCTGGGTA CGTGCAATGACTCTACATTTATTTATTT |
| 34 | GTCTTGAGCAAATAGCAGGTGACAATCG CAAGAAGTAACACGTAAAATAAATAAAT | TCCATCTTGTCTGTTAGCAAGCTGATCG CAAGAAGTAACACGTAATTTATTTATTT |
| 35 | GTCTTGAGCAAATAGCAGGTGACACTAA CTGTACTATCCGCCTGAAATAAATAAAT | TCCATCTTGTCTGTTAGCAAGCTGCTAA CTGTACTATCCGCCTGATTTATTTATTT |
| 36 | GTCTTGAGCAAATAGCAGGTGACAAGTA AGAGGAGTGCCTTATTAAATAAATAAAT | TCCATCTTGTCTGTTAGCAAGCTGAGTA AGAGGAGTGCCTTATTATTTATTTATTT |
| 37 | GTCTTGAGCAAATAGCAGGTGACATTAA GTGACCTGGAGGATATAAATAAATAAAT | TCCATCTTGTCTGTTAGCAAGCTGTTAA GTGACCTGGAGGATATATTTATTTATTT |
| 38 | GTCTTGAGCAAATAGCAGGTGACACCGG ATTGTAAAGTGTCTCAAAATAAATAAAT | TCCATCTTGTCTGTTAGCAAGCTGCCGG ATTGTAAAGTGTCTCAATTTATTTATTT |
| 39 | GTCTTGAGCAAATAGCAGGTGACACGAT TACCTCATATGCCTCTAAATAAATAAAT | TCCATCTTGTCTGTTAGCAAGCTGCGAT TACCTCATATGCCTCTATTTATTTATTT |
| 40 | GTCTTGAGCAAATAGCAGGTGACAAAGT TGCTAGTGGTGATCTAAAATAAATAAAT | TCCATCTTGTCTGTTAGCAAGCTGAAGT TGCTAGTGGTGATCTAATTTATTTATTT |
| 41 | GTCTTGAGCAAATAGCAGGTGACACTTC CTATTGATCCGAGAGGAAATAAATAAAT | TCCATCTTGTCTGTTAGCAAGCTGCTTC CTATTGATCCGAGAGGATTTATTTATTT |
| 42 | GTCTTGAGCAAATAGCAGGTGACAAATA CTTGCCAACACACTTCAAATAAATAAAT | TCCATCTTGTCTGTTAGCAAGCTGAATA CTTGCCAACACACTTCATTTATTTATTT |
| 43 | GTCTTGAGCAAATAGCAGGTGACAACCT ATCAACACACCGTTAAAAATAAATAAAT | TCCATCTTGTCTGTTAGCAAGCTGACCT ATCAACACACCGTTAAATTTATTTATTT |
| 44 | GTCTTGAGCAAATAGCAGGTGACACCTA GTGTCCAGACAATGAGAAATAAATAAAT | TCCATCTTGTCTGTTAGCAAGCTGCCTA GTGTCCAGACAATGAGATTTATTTATTT |
| 45 | GTCTTGAGCAAATAGCAGGTGACACTCA GTGGATGCGATGAATTAAATAAATAAAT | TCCATCTTGTCTGTTAGCAAGCTGCTCA GTGGATGCGATGAATTATTTATTTATTT |
| 46 | GTCTTGAGCAAATAGCAGGTGACACCAG TCTCTTAAGATGCCAAAAATAAATAAAT | TCCATCTTGTCTGTTAGCAAGCTGCCAG TCTCTTAAGATGCCAAATTTATTTATTT |
| 47 | GTCTTGAGCAAATAGCAGGTGACATCGA GTATCAACATGGAACCAAATAAATAAAT | TCCATCTTGTCTGTTAGCAAGCTGTCGA GTATCAACATGGAACCATTTATTTATTT |
| 48 | GTCTTGAGCAAATAGCAGGTGACAAGGA TAAGGACTTATGTGCGAAATAAATAAAT | TCCATCTTGTCTGTTAGCAAGCTGAGGA TAAGGACTTATGTGCGATTTATTTATTT |
| 49 | GTCTTGAGCAAATAGCAGGTGACATCAT CCAGGTTAATGACGACAAATAAATAAAT | TCCATCTTGTCTGTTAGCAAGCTGTCAT CCAGGTTAATGACGACATTTATTTATTT |
| 50 | GTCTTGAGCAAATAGCAGGTGACATAGT TATCGCTACTGTGGAAAAATAAATAAAT | TCCATCTTGTCTGTTAGCAAGCTGTAGT TATCGCTACTGTGGAAATTTATTTATTT |
| 51 | GTCTTGAGCAAATAGCAGGTGACATTAC ACCAACGTGCTGTAAGAAATAAATAAAT | TCCATCTTGTCTGTTAGCAAGCTGTTAC ACCAACGTGCTGTAAGATTTATTTATTT |
| 52 | GTCTTGAGCAAATAGCAGGTGACACCAC GGTCTACGATCATAAGAAATAAATAAAT | TCCATCTTGTCTGTTAGCAAGCTGCCAC GGTCTACGATCATAAGATTTATTTATTT |
| 53 | GTCTTGAGCAAATAGCAGGTGACAAACA ATCGTGTGGAATACTCAAATAAATAAAT | TCCATCTTGTCTGTTAGCAAGCTGAACA ATCGTGTGGAATACTCATTTATTTATTT |
| 54 | GTCTTGAGCAAATAGCAGGTGACACGAG AAGAGAGTCGTTCCTAAAATAAATAAAT | TCCATCTTGTCTGTTAGCAAGCTGCGAG AAGAGAGTCGTTCCTAATTTATTTATTT |
| 55 | GTCTTGAGCAAATAGCAGGTGACAATCG ACAACTCTATTGCATGAAATAAATAAAT | TCCATCTTGTCTGTTAGCAAGCTGATCG ACAACTCTATTGCATGATTTATTTATTT |
| 56 | GTCTTGAGCAAATAGCAGGTGACAAGCC AATTACACAAGGTAGTAAATAAATAAAT | TCCATCTTGTCTGTTAGCAAGCTGAGCC AATTACACAAGGTAGTATTTATTTATTT |
| 57 | GTCTTGAGCAAATAGCAGGTGACAGTCG TCGTTGTCGACATTAGAAATAAATAAAT | TCCATCTTGTCTGTTAGCAAGCTGGTCG TCGTTGTCGACATTAGATTTATTTATTT |
| 58 | GTCTTGAGCAAATAGCAGGTGACACGAC TAGGAGTAGGTTAGGAAAATAAATAAAT | TCCATCTTGTCTGTTAGCAAGCTGCGAC TAGGAGTAGGTTAGGAATTTATTTATTT |

| Location ID | Barcoded recording primer type 'a' | Barcoded recording primer type 'a*' |
|---|---|---|
| 59 | GTCTTGAGCAAATAGCAGGTGACATTGA<br>ATCGCAAATCGACTCAAAATAAATAAAT | TCCATCTTGTCTGTTAGCAAGCTGTTGA<br>ATCGCAAATCGACTCAATTTATTTATTT |
| 60 | GTCTTGAGCAAATAGCAGGTGACAACAA<br>CCAGTATGGTATTCGTAAATAAATAAAT | TCCATCTTGTCTGTTAGCAAGCTGACAA<br>CCAGTATGGTATTCGTATTTATTTATTT |
| 61 | GTCTTGAGCAAATAGCAGGTGACATGGA<br>TTCAAGGCATAGTACAAAATAAATAAAT | TCCATCTTGTCTGTTAGCAAGCTGTGGA<br>TTCAAGGCATAGTACAATTTATTTATTT |
| 62 | GTCTTGAGCAAATAGCAGGTGACACATC<br>TCACGGCTTAGTTACCAAATAAATAAAT | TCCATCTTGTCTGTTAGCAAGCTGCATC<br>TCACGGCTTAGTTACCATTTATTTATTT |
| 63 | GTCTTGAGCAAATAGCAGGTGACAGTTA<br>GGAATGAGCCAAGGAAAAATAAATAAAT | TCCATCTTGTCTGTTAGCAAGCTGGTTA<br>GGAATGAGCCAAGGAAATTTATTTATTT |
| 64 | GTCTTGAGCAAATAGCAGGTGACACGTC<br>GGATGTAAGATAGAGCAAATAAATAAAT | TCCATCTTGTCTGTTAGCAAGCTGCGTC<br>GGATGTAAGATAGAGCATTTATTTATTT |
| 65 | GTCTTGAGCAAATAGCAGGTGACACACC<br>AATAACGTGAACTGTGAAATAAATAAAT | TCCATCTTGTCTGTTAGCAAGCTGCACC<br>AATAACGTGAACTGTGATTTATTTATTT |
| 66 | GTCTTGAGCAAATAGCAGGTGACACACG<br>TTCAGAACATTCTCTCAAATAAATAAAT | TCCATCTTGTCTGTTAGCAAGCTGCACG<br>TTCAGAACATTCTCTCATTTATTTATTT |
| 67 | GTCTTGAGCAAATAGCAGGTGACATAGT<br>CGCTCTATTGTGAACCAAATAAATAAAT | TCCATCTTGTCTGTTAGCAAGCTGTAGT<br>CGCTCTATTGTGAACCATTTATTTATTT |
| 68 | GTCTTGAGCAAATAGCAGGTGACACTTG<br>TTGAGATACCTCGACAAAATAAATAAAT | TCCATCTTGTCTGTTAGCAAGCTGCTTG<br>TTGAGATACCTCGACAATTTATTTATTT |
| 69 | GTCTTGAGCAAATAGCAGGTGACATTCT<br>GGTATGTTCAACAAGCAAATAAATAAAT | TCCATCTTGTCTGTTAGCAAGCTGTTCT<br>GGTATGTTCAACAAGCATTTATTTATTT |
| 70 | GTCTTGAGCAAATAGCAGGTGACAAAGC<br>TCAGTTTGTTCAGAGGAAATAAATAAAT | TCCATCTTGTCTGTTAGCAAGCTGAAGC<br>TCAGTTTGTTCAGAGGATTTATTTATTT |
| 71 | GTCTTGAGCAAATAGCAGGTGACACCGG<br>TTATTACCTACCTGTTAAATAAATAAAT | TCCATCTTGTCTGTTAGCAAGCTGCCGG<br>TTATTACCTACCTGTTATTTATTTATTT |
| 72 | GTCTTGAGCAAATAGCAGGTGACACGAC<br>TTATTGAGTCATTGCCAAATAAATAAAT | TCCATCTTGTCTGTTAGCAAGCTGCGAC<br>TTATTGAGTCATTGCCATTTATTTATTT |
| 73 | GTCTTGAGCAAATAGCAGGTGACAGATA<br>CGTCTATTGAACACCGAAATAAATAAAT | TCCATCTTGTCTGTTAGCAAGCTGGATA<br>CGTCTATTGAACACCGATTTATTTATTT |
| 74 | GTCTTGAGCAAATAGCAGGTGACAATCT<br>GAGTACCGAATAGTCCAAATAAATAAAT | TCCATCTTGTCTGTTAGCAAGCTGATCT<br>GAGTACCGAATAGTCCATTTATTTATTT |
| 75 | GTCTTGAGCAAATAGCAGGTGACAGTGT<br>TGATGGACGTAGTTGAAAATAAATAAAT | TCCATCTTGTCTGTTAGCAAGCTGGTGT<br>TGATGGACGTAGTTGAATTTATTTATTT |
| 76 | GTCTTGAGCAAATAGCAGGTGACAAACA<br>GAGTGATTGTCAGTTCAAATAAATAAAT | TCCATCTTGTCTGTTAGCAAGCTGAACA<br>GAGTGATTGTCAGTTCATTTATTTATTT |
| 77 | GTCTTGAGCAAATAGCAGGTGACACTAT<br>ATGAGGGGTTGCTCTCAAATAAATAAAT | TCCATCTTGTCTGTTAGCAAGCTGCTAT<br>ATGAGGGGTTGCTCTCATTTATTTATTT |
| 78 | GTCTTGAGCAAATAGCAGGTGACACACT<br>AGAACCCTGCTTATGCAAATAAATAAAT | TCCATCTTGTCTGTTAGCAAGCTGCACT<br>AGAACCCTGCTTATGCATTTATTTATTT |
| 79 | GTCTTGAGCAAATAGCAGGTGACATCAG<br>GTTCTAGATGGTCTTCAAATAAATAAAT | TCCATCTTGTCTGTTAGCAAGCTGTCAG<br>GTTCTAGATGGTCTTCATTTATTTATTT |
| 80 | GTCTTGAGCAAATAGCAGGTGACAGCAA<br>GACTATCGCCTAGTTCAAATAAATAAAT | TCCATCTTGTCTGTTAGCAAGCTGGCAA<br>GACTATCGCCTAGTTCATTTATTTATTT |
| 81 | GTCTTGAGCAAATAGCAGGTGACAAGTG<br>GTGATTACATTAGCCAAAATAAATAAAT | TCCATCTTGTCTGTTAGCAAGCTGAGTG<br>GTGATTACATTAGCCAATTTATTTATTT |
| 82 | GTCTTGAGCAAATAGCAGGTGACAAGGA<br>TTGAACGAAGCTGTTAAAATAAATAAAT | TCCATCTTGTCTGTTAGCAAGCTGAGGA<br>TTGAACGAAGCTGTTAATTTATTTATTT |
| 83 | GTCTTGAGCAAATAGCAGGTGACAATTG<br>TGTTGCGAGTACGATAAAATAAATAAAT | TCCATCTTGTCTGTTAGCAAGCTGATTG<br>TGTTGCGAGTACGATAATTTATTTATTT |
| 84 | GTCTTGAGCAAATAGCAGGTGACAGTAC<br>GAATAGACACTCGCTGAAATAAATAAAT | TCCATCTTGTCTGTTAGCAAGCTGGTAC<br>GAATAGACACTCGCTGATTTATTTATTT |
| 85 | GTCTTGAGCAAATAGCAGGTGACAAGAA<br>CATTGGTTCGGATTGAAAATAAATAAAT | TCCATCTTGTCTGTTAGCAAGCTGAGAA<br>CATTGGTTCGGATTGAATTTATTTATTT |
| 86 | GTCTTGAGCAAATAGCAGGTGACAGTTG<br>TACGCTTTACGTTGTGAAATAAATAAAT | TCCATCTTGTCTGTTAGCAAGCTGGTTG<br>TACGCTTTACGTTGTGATTTATTTATTT |
| 87 | GTCTTGAGCAAATAGCAGGTGACATAGC<br>ACGATTAGATCGGAATAAATAAATAAAT | TCCATCTTGTCTGTTAGCAAGCTGTAGC<br>ACGATTAGATCGGAATATTTATTTATTT |

| Location ID | Barcoded recording primer type 'a' | Barcoded recording primer type 'a*' |
|---|---|---|
| 88 | GTCTTGAGCAAATAGCAGGTGACACTCA<br>AGCTCATAGTACCTGAAAATAAATAAAT | TCCATCTTGTCTGTTAGCAAGCTGCTCA<br>AGCTCATAGTACCTGAATTTATTTATTT |
| 89 | GTCTTGAGCAAATAGCAGGTGACACATG<br>TAAGACGACTTACGTGAAATAAATAAAT | TCCATCTTGTCTGTTAGCAAGCTGCATG<br>TAAGACGACTTACGTGATTTATTTATTT |
| 90 | GTCTTGAGCAAATAGCAGGTGACATTCG<br>CAATGGATAAGTCTCCAAATAAATAAAT | TCCATCTTGTCTGTTAGCAAGCTGTTCG<br>CAATGGATAAGTCTCCATTTATTTATTT |
| 91 | GTCTTGAGCAAATAGCAGGTGACAGACA<br>CAATACACACCACTCAAAATAAATAAAT | TCCATCTTGTCTGTTAGCAAGCTGGACA<br>CAATACACACCACTCAATTTATTTATTT |
| 92 | GTCTTGAGCAAATAGCAGGTGACACTAT<br>CGAGAAAAGTACGCCTAAATAAATAAAT | TCCATCTTGTCTGTTAGCAAGCTGCTAT<br>CGAGAAAAGTACGCCTATTTATTTATTT |
| 93 | GTCTTGAGCAAATAGCAGGTGACATCAC<br>TGGTTACCTTCAATCAAAATAAATAAAT | TCCATCTTGTCTGTTAGCAAGCTGTCAC<br>TGGTTACCTTCAATCAATTTATTTATTT |
| 94 | GTCTTGAGCAAATAGCAGGTGACAAGTT<br>GCATGTGGATTCACTAAAATAAATAAAT | TCCATCTTGTCTGTTAGCAAGCTGAGTT<br>GCATGTGGATTCACTAATTTATTTATTT |
| 95 | GTCTTGAGCAAATAGCAGGTGACACACT<br>AGATTGCTCCAGTACAAAATAAATAAAT | TCCATCTTGTCTGTTAGCAAGCTGCACT<br>AGATTGCTCCAGTACAATTTATTTATTT |
| 96 | GTCTTGAGCAAATAGCAGGTGACAGGTT<br>ATGTGGATAGCTTGAGAAATAAATAAAT | TCCATCTTGTCTGTTAGCAAGCTGGGTT<br>ATGTGGATAGCTTGAGATTTATTTATTT |
| 97 | GTCTTGAGCAAATAGCAGGTGACATTGT<br>CTCGAAAGACGTAAGTAAATAAATAAAT | TCCATCTTGTCTGTTAGCAAGCTGTTGT<br>CTCGAAAGACGTAAGTATTTATTTATTT |
| 98 | GTCTTGAGCAAATAGCAGGTGACAACCA<br>CCAATTGATGATTGAGAAATAAATAAAT | TCCATCTTGTCTGTTAGCAAGCTGACCA<br>CCAATTGATGATTGAGATTTATTTATTT |
| 99 | GTCTTGAGCAAATAGCAGGTGACAATTA<br>CACTCCGTCCAAGTAAAAATAAATAAAT | TCCATCTTGTCTGTTAGCAAGCTGATTA<br>CACTCCGTCCAAGTAAATTTATTTATTT |
| 100 | GTCTTGAGCAAATAGCAGGTGACAACGG<br>TACTGTCTAAGATTGCAAATAAATAAAT | TCCATCTTGTCTGTTAGCAAGCTGACGG<br>TACTGTCTAAGATTGCATTTATTTATTT |
| 101 | GTCTTGAGCAAATAGCAGGTGACAAGAA<br>CTCACAATTCTGATGGAAATAAATAAAT | TCCATCTTGTCTGTTAGCAAGCTGAGAA<br>CTCACAATTCTGATGGATTTATTTATTT |
| 102 | GTCTTGAGCAAATAGCAGGTGACATATA<br>AGCACCATGAGGACTAAAATAAATAAAT | TCCATCTTGTCTGTTAGCAAGCTGTATA<br>AGCACCATGAGGACTAATTTATTTATTT |
| 103 | GTCTTGAGCAAATAGCAGGTGACACACT<br>ACGTCCTATACCTCCAAAATAAATAAAT | TCCATCTTGTCTGTTAGCAAGCTGCACT<br>ACGTCCTATACCTCCAATTTATTTATTT |
| 104 | GTCTTGAGCAAATAGCAGGTGACAGCTT<br>ACACTAGAAGCCGAAGAAATAAATAAAT | TCCATCTTGTCTGTTAGCAAGCTGGCTT<br>ACACTAGAAGCCGAAGATTTATTTATTT |
| 105 | GTCTTGAGCAAATAGCAGGTGACAACAT<br>TGAGAGCATTGTACCAAAATAAATAAAT | TCCATCTTGTCTGTTAGCAAGCTGACAT<br>TGAGAGCATTGTACCAATTTATTTATTT |
| 106 | GTCTTGAGCAAATAGCAGGTGACACACC<br>TCGTAGTGCGTTATAGAAATAAATAAAT | TCCATCTTGTCTGTTAGCAAGCTGCACC<br>TCGTAGTGCGTTATAGATTTATTTATTT |
| 107 | GTCTTGAGCAAATAGCAGGTGACACGAT<br>GTAGCAACGAATCATCAAATAAATAAAT | TCCATCTTGTCTGTTAGCAAGCTGCGAT<br>GTAGCAACGAATCATCATTTATTTATTT |
| 108 | GTCTTGAGCAAATAGCAGGTGACACGAG<br>GTGTAATAAGTTCGCAAAATAAATAAAT | TCCATCTTGTCTGTTAGCAAGCTGCGAG<br>GTGTAATAAGTTCGCAATTTATTTATTT |
| 109 | GTCTTGAGCAAATAGCAGGTGACAGTCA<br>TCAAGACGGTTATCGGAAATAAATAAAT | TCCATCTTGTCTGTTAGCAAGCTGGTCA<br>TCAAGACGGTTATCGGATTTATTTATTT |
| 110 | GTCTTGAGCAAATAGCAGGTGACAATCC<br>TTGCAGATAACGTAGGAAATAAATAAAT | TCCATCTTGTCTGTTAGCAAGCTGATCC<br>TTGCAGATAACGTAGGATTTATTTATTT |
| 111 | GTCTTGAGCAAATAGCAGGTGACAAACT<br>GATCTGCGCTAATGAAAAATAAATAAAT | TCCATCTTGTCTGTTAGCAAGCTGAACT<br>GATCTGCGCTAATGAAATTTATTTATTT |
| 112 | GTCTTGAGCAAATAGCAGGTGACAGAGG<br>TCTGGAAGTAACCTTCAAATAAATAAAT | TCCATCTTGTCTGTTAGCAAGCTGGAGG<br>TCTGGAAGTAACCTTCATTTATTTATTT |
| 113 | GTCTTGAGCAAATAGCAGGTGACAACCA<br>AGTATCCACTGACATAAAATAAATAAAT | TCCATCTTGTCTGTTAGCAAGCTGACCA<br>AGTATCCACTGACATAATTTATTTATTT |
| 114 | GTCTTGAGCAAATAGCAGGTGACAACGA<br>ACCTAAAGCGTGTTAAAAATAAATAAAT | TCCATCTTGTCTGTTAGCAAGCTGACGA<br>ACCTAAAGCGTGTTAAATTTATTTATTT |
| 115 | GTCTTGAGCAAATAGCAGGTGACAATAC<br>TACGAGAACTTCCGAGAAATAAATAAAT | TCCATCTTGTCTGTTAGCAAGCTGATAC<br>TACGAGAACTTCCGAGATTTATTTATTT |
| 116 | GTCTTGAGCAAATAGCAGGTGACAGTGG<br>ATCGAATGTTCTGACTAAATAAATAAAT | TCCATCTTGTCTGTTAGCAAGCTGGTGG<br>ATCGAATGTTCTGACTATTTATTTATTT |

| Location ID | Barcoded recording primer type 'a' | Barcoded recording primer type 'a*' |
|---|---|---|
| 117 | GTCTTGAGCAAATAGCAGGTGACAAATG TCCTTGATGTGACTCGAAATAAATAAAT | TCCATCTTGTCTGTTAGCAAGCTGAATG TCCTTGATGTGACTCGATTTATTTATTT |
| 118 | GTCTTGAGCAAATAGCAGGTGACACGGT AGGTTCGAACTTGTACAAATAAATAAAT | TCCATCTTGTCTGTTAGCAAGCTGCGGT AGGTTCGAACTTGTACATTTATTTATTT |
| 119 | GTCTTGAGCAAATAGCAGGTGACAGAGT TCAGCAGTATCGTAGCAAATAAATAAAT | TCCATCTTGTCTGTTAGCAAGCTGGAGT TCAGCAGTATCGTAGCATTTATTTATTT |
| 120 | GTCTTGAGCAAATAGCAGGTGACATAGA CAGACTATACGTGTCGAAATAAATAAAT | TCCATCTTGTCTGTTAGCAAGCTGTAGA CAGACTATACGTGTCGATTTATTTATTT |
| 121 | GTCTTGAGCAAATAGCAGGTGACAAAGA CCTCAAAACGGATGTCAAATAAATAAAT | TCCATCTTGTCTGTTAGCAAGCTGAAGA CCTCAAAACGGATGTCATTTATTTATTT |
| 122 | GTCTTGAGCAAATAGCAGGTGACAAGTA AGATCCCTAATACGCTAAATAAATAAAT | TCCATCTTGTCTGTTAGCAAGCTGAGTA AGATCCCTAATACGCTATTTATTTATTT |
| 123 | GTCTTGAGCAAATAGCAGGTGACATCTA GCCAAGTACGCTTAGAAAATAAATAAAT | TCCATCTTGTCTGTTAGCAAGCTGTCTA GCCAAGTACGCTTAGAATTTATTTATTT |
| 124 | GTCTTGAGCAAATAGCAGGTGACAATAT CAGTCCAGGAACAGTCAAATAAATAAAT | TCCATCTTGTCTGTTAGCAAGCTGATAT CAGTCCAGGAACAGTCATTTATTTATTT |
| 125 | GTCTTGAGCAAATAGCAGGTGACATGTA GTGTTCCTTCTATCCGAAATAAATAAAT | TCCATCTTGTCTGTTAGCAAGCTGTGTA GTGTTCCTTCTATCCGATTTATTTATTT |
| 126 | GTCTTGAGCAAATAGCAGGTGACACTAA CCACTAGGACTAGCTCAAATAAATAAAT | TCCATCTTGTCTGTTAGCAAGCTGCTAA CCACTAGGACTAGCTCATTTATTTATTT |
| 127 | GTCTTGAGCAAATAGCAGGTGACACGCA TTGATAATCCGCTACAAAATAAATAAAT | TCCATCTTGTCTGTTAGCAAGCTGCGCA TTGATAATCCGCTACAATTTATTTATTT |
| 128 | GTCTTGAGCAAATAGCAGGTGACAAATC ATCGACGAACAGCTAAAAATAAATAAAT | TCCATCTTGTCTGTTAGCAAGCTGAATC ATCGACGAACAGCTAAATTTATTTATTT |
| 129 | GTCTTGAGCAAATAGCAGGTGACAAGGA TACTACGAAGAATGGTAAATAAATAAAT | TCCATCTTGTCTGTTAGCAAGCTGAGGA TACTACGAAGAATGGTATTTATTTATTT |
| 130 | GTCTTGAGCAAATAGCAGGTGACATCGT AAGGAAGTAGACTGAAAAATAAATAAAT | TCCATCTTGTCTGTTAGCAAGCTGTCGT AAGGAAGTAGACTGAAATTTATTTATTT |
| 131 | GTCTTGAGCAAATAGCAGGTGACAAACG TACTAGATGGTGACAGAAATAAATAAAT | TCCATCTTGTCTGTTAGCAAGCTGAACG TACTAGATGGTGACAGATTTATTTATTT |
| 132 | GTCTTGAGCAAATAGCAGGTGACAAACC GTATTCCAGGTTACAAAAATAAATAAAT | TCCATCTTGTCTGTTAGCAAGCTGAACC GTATTCCAGGTTACAAATTTATTTATTT |
| 133 | GTCTTGAGCAAATAGCAGGTGACAAACA CAAGAGAGAATACGCAAAATAAATAAAT | TCCATCTTGTCTGTTAGCAAGCTGAACA CAAGAGAGAATACGCAATTTATTTATTT |
| 134 | GTCTTGAGCAAATAGCAGGTGACAGAGG TATATCCGGTGATACAAAATAAATAAAT | TCCATCTTGTCTGTTAGCAAGCTGGAGG TATATCCGGTGATACAATTTATTTATTT |
| 135 | GTCTTGAGCAAATAGCAGGTGACAGCTT CTAATGACGGTCCATTAAATAAATAAAT | TCCATCTTGTCTGTTAGCAAGCTGGCTT CTAATGACGGTCCATTATTTATTTATTT |
| 136 | GTCTTGAGCAAATAGCAGGTGACAAAGA TTCAGGCCAACTATTCAAATAAATAAAT | TCCATCTTGTCTGTTAGCAAGCTGAAGA TTCAGGCCAACTATTCATTTATTTATTT |
| 137 | GTCTTGAGCAAATAGCAGGTGACAATAA CATCGCTAGTAAGCGTAAATAAATAAAT | TCCATCTTGTCTGTTAGCAAGCTGATAA CATCGCTAGTAAGCGTATTTATTTATTT |
| 138 | GTCTTGAGCAAATAGCAGGTGACACCTC TAATACCTAACCTCGGAAATAAATAAAT | TCCATCTTGTCTGTTAGCAAGCTGCCTC TAATACCTAACCTCGGATTTATTTATTT |
| 139 | GTCTTGAGCAAATAGCAGGTGACACGCT ATGGTATCGGATCTTAAAATAAATAAAT | TCCATCTTGTCTGTTAGCAAGCTGCGCT ATGGTATCGGATCTTAATTTATTTATTT |
| 140 | GTCTTGAGCAAATAGCAGGTGACACATA ACAGACTTGTTCCAGCAAATAAATAAAT | TCCATCTTGTCTGTTAGCAAGCTGCATA ACAGACTTGTTCCAGCATTTATTTATTT |
| 141 | GTCTTGAGCAAATAGCAGGTGACAGCCT CTGCTTCCTTATATGGAAATAAATAAAT | TCCATCTTGTCTGTTAGCAAGCTGGCCT CTGCTTCCTTATATGGATTTATTTATTT |
| 142 | GTCTTGAGCAAATAGCAGGTGACACCAC TCATCGTAAGTCATGGAAATAAATAAAT | TCCATCTTGTCTGTTAGCAAGCTGCCAC TCATCGTAAGTCATGGATTTATTTATTT |
| 143 | GTCTTGAGCAAATAGCAGGTGACAGAAC ACCACAGTGAATACCTAAATAAATAAAT | TCCATCTTGTCTGTTAGCAAGCTGGAAC ACCACAGTGAATACCTATTTATTTATTT |
| 144 | GTCTTGAGCAAATAGCAGGTGACAAGCG AAGATTGTCTACTGAGAAATAAATAAAT | TCCATCTTGTCTGTTAGCAAGCTGAGCG AAGATTGTCTACTGAGATTTATTTATTT |
| 145 | GTCTTGAGCAAATAGCAGGTGACACGCA CATGGACTGTAGTATGAAATAAATAAAT | TCCATCTTGTCTGTTAGCAAGCTGCGCA CATGGACTGTAGTATGATTTATTTATTT |

| Location ID | Barcoded recording primer type 'a' | Barcoded recording primer type 'a*' |
|---|---|---|
| 146 | GTCTTGAGCAAATAGCAGGTGACACGAATTGACGGAAGTGAGTTAAATAAATAAAT | TCCATCTTGTCTGTTAGCAAGCTGCGAATTGACGGAAGTGAGTTATTTATTTATTT |
| 147 | GTCTTGAGCAAATAGCAGGTGACACGAGGAGTCTATTGATAGCCAAATAAATAAAT | TCCATCTTGTCTGTTAGCAAGCTGCGAGGAGTCTATTGATAGCCATTTATTTATTT |
| 148 | GTCTTGAGCAAATAGCAGGTGACACTACGAGTGACACAGGTATAAAATAAATAAAT | TCCATCTTGTCTGTTAGCAAGCTGCTACGAGTGACACAGGTATAATTTATTTATTT |
| 149 | GTCTTGAGCAAATAGCAGGTGACATCGGAATAGAATAGCGGATCAAATAAATAAAT | TCCATCTTGTCTGTTAGCAAGCTGTCGGAATAGAATAGCGGATCATTTATTTATTT |
| 150 | GTCTTGAGCAAATAGCAGGTGACATACATCTCCGGAACGTATGAAAATAAATAAAT | TCCATCTTGTCTGTTAGCAAGCTGTACATCTCCGGAACGTATGAATTTATTTATTT |
| 151 | GTCTTGAGCAAATAGCAGGTGACAATCAATCACCCCTATTCACCAAATAAATAAAT | TCCATCTTGTCTGTTAGCAAGCTGATCAATCACCCCTATTCACCATTTATTTATTT |
| 152 | GTCTTGAGCAAATAGCAGGTGACATCTGTGTTACATCTAAGCGTAAATAAATAAAT | TCCATCTTGTCTGTTAGCAAGCTGTCTGTGTTACATCTAAGCGTATTTATTTATTT |
| 153 | GTCTTGAGCAAATAGCAGGTGACATCGATGACCAAATACCGAAGAAATAAATAAAT | TCCATCTTGTCTGTTAGCAAGCTGTCGATGACCAAATACCGAAGATTTATTTATTT |
| 154 | GTCTTGAGCAAATAGCAGGTGACATACTCATGTGGTAGGTCAATAAATAAATAAAT | TCCATCTTGTCTGTTAGCAAGCTGTACTCATGTGGTAGGTCAATATTTATTTATTT |
| 155 | GTCTTGAGCAAATAGCAGGTGACAATTCTCCAAGGCCTACAATGAAATAAATAAAT | TCCATCTTGTCTGTTAGCAAGCTGATTCTCCAAGGCCTACAATGATTTATTTATTT |
| 156 | GTCTTGAGCAAATAGCAGGTGACAGAGCCTATGTTTAGGTACGCAAATAAATAAAT | TCCATCTTGTCTGTTAGCAAGCTGGAGCCTATGTTTAGGTACGCATTTATTTATTT |
| 157 | GTCTTGAGCAAATAGCAGGTGACAATGTGTCAAGGTATCACATGAAATAAATAAAT | TCCATCTTGTCTGTTAGCAAGCTGATGTGTCAAGGTATCACATGATTTATTTATTT |
| 158 | GTCTTGAGCAAATAGCAGGTGACATACCTGATGAAGTGATGCAAAAATAAATAAAT | TCCATCTTGTCTGTTAGCAAGCTGTACCTGATGAAGTGATGCAAATTTATTTATTT |
| 159 | GTCTTGAGCAAATAGCAGGTGACACATAAGGTCATGTGCTTGAAAAATAAATAAAT | TCCATCTTGTCTGTTAGCAAGCTGCATAAGGTCATGTGCTTGAAATTTATTTATTT |
| 160 | GTCTTGAGCAAATAGCAGGTGACACTGTGACGGATGACAATACGAAATAAATAAAT | TCCATCTTGTCTGTTAGCAAGCTGCTGTGACGGATGACAATACGATTTATTTATTT |
| 161 | GTCTTGAGCAAATAGCAGGTGACACCATGAAGGAAACAGTAGCAAAATAAATAAAT | TCCATCTTGTCTGTTAGCAAGCTGCCATGAAGGAAACAGTAGCAATTTATTTATTT |
| 162 | GTCTTGAGCAAATAGCAGGTGACAGACACTCGAAAAGGTAACTCAAATAAATAAAT | TCCATCTTGTCTGTTAGCAAGCTGGACACTCGAAAAGGTAACTCATTTATTTATTT |
| 163 | GTCTTGAGCAAATAGCAGGTGACATTGTTGCTACAACAGACCTAAAATAAATAAAT | TCCATCTTGTCTGTTAGCAAGCTGTTGTTGCTACAACAGACCTAATTTATTTATTT |
| 164 | GTCTTGAGCAAATAGCAGGTGACATAGTGGTTGGCTCCTCTATAAAATAAATAAAT | TCCATCTTGTCTGTTAGCAAGCTGTAGTGGTTGGCTCCTCTATAATTTATTTATTT |
| 165 | GTCTTGAGCAAATAGCAGGTGACACATCCATTCCCAATACCATCAAATAAATAAAT | TCCATCTTGTCTGTTAGCAAGCTGCATCCATTCCCAATACCATCATTTATTTATTT |
| 166 | GTCTTGAGCAAATAGCAGGTGACAAAGACGGTATTTCTCAACCGAAATAAATAAAT | TCCATCTTGTCTGTTAGCAAGCTGAAGACGGTATTTCTCAACCGATTTATTTATTT |
| 167 | GTCTTGAGCAAATAGCAGGTGACAATAGCTCAACACTGTTGTGAAAATAAATAAAT | TCCATCTTGTCTGTTAGCAAGCTGATAGCTCAACACTGTTGTGAATTTATTTATTT |
| 168 | GTCTTGAGCAAATAGCAGGTGACAACATCCAAGACTCTAGGTGAAAATAAATAAAT | TCCATCTTGTCTGTTAGCAAGCTGACATCCAAGACTCTAGGTGAATTTATTTATTT |
| 169 | GTCTTGAGCAAATAGCAGGTGACAAAGAGGTCGAGATATGAACCAAATAAATAAAT | TCCATCTTGTCTGTTAGCAAGCTGAAGAGGTCGAGATATGAACCATTTATTTATTT |
| 170 | GTCTTGAGCAAATAGCAGGTGACAGATCGACGTAATTCCATGAGAAATAAATAAAT | TCCATCTTGTCTGTTAGCAAGCTGGATCGACGTAATTCCATGAGATTTATTTATTT |
| 171 | GTCTTGAGCAAATAGCAGGTGACAACAATACTCCTGCACCATATAAATAAATAAAT | TCCATCTTGTCTGTTAGCAAGCTGACAATACTCCTGCACCATATATTTATTTATTT |
| 172 | GTCTTGAGCAAATAGCAGGTGACACTAGGACCATGATGTAATGCAAATAAATAAAT | TCCATCTTGTCTGTTAGCAAGCTGCTAGGACCATGATGTAATGCATTTATTTATTT |
| 173 | GTCTTGAGCAAATAGCAGGTGACAATGGATTCTCCTCGAAGTTGAAATAAATAAAT | TCCATCTTGTCTGTTAGCAAGCTGATGGATTCTCCTCGAAGTTGATTTATTTATTT |
| 174 | GTCTTGAGCAAATAGCAGGTGACACATGTCGAATAAGACCGACCAAATAAATAAAT | TCCATCTTGTCTGTTAGCAAGCTGCATGTCGAATAAGACCGACCATTTATTTATTT |

| Location ID | Barcoded recording primer type 'a' | Barcoded recording primer type 'a*' |
|---|---|---|
| 175 | GTCTTGAGCAAATAGCAGGTGACAGTGTGTGTACAACAACAACCAAATAAATAAAT | TCCATCTTGTCTGTTAGCAAGCTGGTGTGTGTACAACAACAACCATTTATTTATTT |
| 176 | GTCTTGAGCAAATAGCAGGTGACAGATATACGGAACTCCACGTTAAATAAATAAAT | TCCATCTTGTCTGTTAGCAAGCTGGATATACGGAACTCCACGTTATTTATTTATTT |
| 177 | GTCTTGAGCAAATAGCAGGTGACACCTAAGATACCATGGTCCTAAAATAAATAAAT | TCCATCTTGTCTGTTAGCAAGCTGCCTAAGATACCATGGTCCTAATTTATTTATTT |
| 178 | GTCTTGAGCAAATAGCAGGTGACACCTACTCGGAGTAACAATCCAAATAAATAAAT | TCCATCTTGTCTGTTAGCAAGCTGCCTACTCGGAGTAACAATCCATTTATTTATTT |
| 179 | GTCTTGAGCAAATAGCAGGTGACAACACTATTGCAGCTTGATTCAAATAAATAAAT | TCCATCTTGTCTGTTAGCAAGCTGACACTATTGCAGCTTGATTCATTTATTTATTT |
| 180 | GTCTTGAGCAAATAGCAGGTGACACCTAATGCGTCGTAATTGTGAAATAAATAAAT | TCCATCTTGTCTGTTAGCAAGCTGCCTAATGCGTCGTAATTGTGATTTATTTATTT |
| 181 | GTCTTGAGCAAATAGCAGGTGACAACAACACAACAGCAGTTCTAAAATAAATAAAT | TCCATCTTGTCTGTTAGCAAGCTGACAACACAACAGCAGTTCTAATTTATTTATTT |
| 182 | GTCTTGAGCAAATAGCAGGTGACACTATGCCGTTACTGGAATTGAAATAAATAAAT | TCCATCTTGTCTGTTAGCAAGCTGCTATGCCGTTACTGGAATTGATTTATTTATTT |
| 183 | GTCTTGAGCAAATAGCAGGTGACATACACAAGTCCAACATCAGAAAATAAATAAAT | TCCATCTTGTCTGTTAGCAAGCTGTACACAAGTCCAACATCAGAATTTATTTATTT |
| 184 | GTCTTGAGCAAATAGCAGGTGACAAGAACCGTGTAAGTGGATACAAATAAATAAAT | TCCATCTTGTCTGTTAGCAAGCTGAGAACCGTGTAAGTGGATACATTTATTTATTT |
| 185 | GTCTTGAGCAAATAGCAGGTGACATCACACAGAAGGTGATAGTTAAATAAATAAAT | TCCATCTTGTCTGTTAGCAAGCTGTCACACAGAAGGTGATAGTTATTTATTTATTT |
| 186 | GTCTTGAGCAAATAGCAGGTGACAACATAAGTGGCGCTCTTATCAAATAAATAAAT | TCCATCTTGTCTGTTAGCAAGCTGACATAAGTGGCGCTCTTATCATTTATTTATTT |
| 187 | GTCTTGAGCAAATAGCAGGTGACAAAGCCACATGATAGTAGACCAAATAAATAAAT | TCCATCTTGTCTGTTAGCAAGCTGAAGCCACATGATAGTAGACCATTTATTTATTT |
| 188 | GTCTTGAGCAAATAGCAGGTGACAAACGAGCTTAGTGTCCAATAAAATAAATAAAT | TCCATCTTGTCTGTTAGCAAGCTGAACGAGCTTAGTGTCCAATAATTTATTTATTT |
| 189 | GTCTTGAGCAAATAGCAGGTGACATGACAAGCAAATTGAGAAGGAAATAAATAAAT | TCCATCTTGTCTGTTAGCAAGCTGTGACAAGCAAATTGAGAAGGATTTATTTATTT |
| 190 | GTCTTGAGCAAATAGCAGGTGACACTCCAGTGTTGGAATACAGAAAATAAATAAAT | TCCATCTTGTCTGTTAGCAAGCTGCTCCAGTGTTGGAATACAGAATTTATTTATTT |
| 191 | GTCTTGAGCAAATAGCAGGTGACAAGCACACTAAGAGAAGTTGGAAATAAATAAAT | TCCATCTTGTCTGTTAGCAAGCTGAGCACACTAAGAGAAGTTGGATTTATTTATTT |
| 192 | GTCTTGAGCAAATAGCAGGTGACAACAACGACTTTCTAGTCCTCAAATAAATAAAT | TCCATCTTGTCTGTTAGCAAGCTGACAACGACTTTCTAGTCCTCATTTATTTATTT |
| 193 | GTCTTGAGCAAATAGCAGGTGACAGTGTGACTTATTCACCGAGAAAATAAATAAAT | TCCATCTTGTCTGTTAGCAAGCTGGTGTGACTTATTCACCGAGAATTTATTTATTT |
| 194 | GTCTTGAGCAAATAGCAGGTGACACTACAGCCGAATAGGTTCACAAATAAATAAAT | TCCATCTTGTCTGTTAGCAAGCTGCTACAGCCGAATAGGTTCACATTTATTTATTT |
| 195 | GTCTTGAGCAAATAGCAGGTGACAATTATCCGGTACCAATCCTTAAATAAATAAAT | TCCATCTTGTCTGTTAGCAAGCTGATTATCCGGTACCAATCCTTATTTATTTATTT |
| 196 | GTCTTGAGCAAATAGCAGGTGACAATGGTTAGTGAACCACGTACAAATAAATAAAT | TCCATCTTGTCTGTTAGCAAGCTGATGGTTAGTGAACCACGTACATTTATTTATTT |
| 197 | GTCTTGAGCAAATAGCAGGTGACAACGAATGTACTCTACCGTCAAAATAAATAAAT | TCCATCTTGTCTGTTAGCAAGCTGACGAATGTACTCTACCGTCAATTTATTTATTT |
| 198 | GTCTTGAGCAAATAGCAGGTGACAGGATGACCTACAAGACACTAAAATAAATAAAT | TCCATCTTGTCTGTTAGCAAGCTGGGATGACCTACAAGACACTAATTTATTTATTT |
| 199 | GTCTTGAGCAAATAGCAGGTGACACAGTGGACAATACAGGTCAGAAATAAATAAAT | TCCATCTTGTCTGTTAGCAAGCTGCAGTGGACAATACAGGTCAGATTTATTTATTT |
| 200 | GTCTTGAGCAAATAGCAGGTGACACGTGCGTAGATTGCTAGATGAAATAAATAAAT | TCCATCTTGTCTGTTAGCAAGCTGCGTGCGTAGATTGCTAGATGATTTATTTATTT |
| 201 | GTCTTGAGCAAATAGCAGGTGACAGCACATTCTTTTCAGATCCAAAATAAATAAAT | TCCATCTTGTCTGTTAGCAAGCTGGCACATTCTTTTCAGATCCAATTTATTTATTT |
| 202 | GTCTTGAGCAAATAGCAGGTGACAAATAGCAGAGTCGCGTATTGAAATAAATAAAT | TCCATCTTGTCTGTTAGCAAGCTGAATAGCAGAGTCGCGTATTGATTTATTTATTT |
| 203 | GTCTTGAGCAAATAGCAGGTGACACTGGTAGGAAGATCCTCAGAAAATAAATAAAT | TCCATCTTGTCTGTTAGCAAGCTGCTGGTAGGAAGATCCTCAGAATTTATTTATTT |

| Location ID | Barcoded recording primer type ‘a’ | Barcoded recording primer type ‘a*’ |
|---|---|---|
| 204 | GTCTTGAGCAAATAGCAGGTGACACTTAGACACATGTGCGAGTTAAATAAATAAAT | TCCATCTTGTCTGTTAGCAAGCTGCTTAGACACATGTGCGAGTTATTTATTTATTT |
| 205 | GTCTTGAGCAAATAGCAGGTGACACATGAGTAACTGTCGGTCACAAATAAATAAAT | TCCATCTTGTCTGTTAGCAAGCTGCATGAGTAACTGTCGGTCACATTTATTTATTT |
| 206 | GTCTTGAGCAAATAGCAGGTGACAAATGTCGTACGCAATGTATCAAATAAATAAAT | TCCATCTTGTCTGTTAGCAAGCTGAATGTCGTACGCAATGTATCATTTATTTATTT |
| 207 | GTCTTGAGCAAATAGCAGGTGACAAGAATTGCTCGGTGTCATACAAATAAATAAAT | TCCATCTTGTCTGTTAGCAAGCTGAGAATTGCTCGGTGTCATACATTTATTTATTT |
| 208 | GTCTTGAGCAAATAGCAGGTGACAACCGTTCGATATTAGAGCTGAAATAAATAAAT | TCCATCTTGTCTGTTAGCAAGCTGACCGTTCGATATTAGAGCTGATTTATTTATTT |
| 209 | GTCTTGAGCAAATAGCAGGTGACATATACTGCGGTTGTCATAGCAAATAAATAAAT | TCCATCTTGTCTGTTAGCAAGCTGTATACTGCGGTTGTCATAGCATTTATTTATTT |
| 210 | GTCTTGAGCAAATAGCAGGTGACAACTTGTCTACACAACAATGCAAATAAATAAAT | TCCATCTTGTCTGTTAGCAAGCTGACTTGTCTACACAACAATGCATTTATTTATTT |
| 211 | GTCTTGAGCAAATAGCAGGTGACAGACCGTTGTTCAATCAGTTCAAATAAATAAAT | TCCATCTTGTCTGTTAGCAAGCTGGACCGTTGTTCAATCAGTTCATTTATTTATTT |
| 212 | GTCTTGAGCAAATAGCAGGTGACACCTGTCATTGAATGTTCCACAAATAAATAAAT | TCCATCTTGTCTGTTAGCAAGCTGCCTGTCATTGAATGTTCCACATTTATTTATTT |
| 213 | GTCTTGAGCAAATAGCAGGTGACAACCACTTGAACATTGATTCGAAATAAATAAAT | TCCATCTTGTCTGTTAGCAAGCTGACCACTTGAACATTGATTCGATTTATTTATTT |
| 214 | GTCTTGAGCAAATAGCAGGTGACAGGAGACTTCGTGAACAGATCAAATAAATAAAT | TCCATCTTGTCTGTTAGCAAGCTGGGAGACTTCGTGAACAGATCATTTATTTATTT |
| 215 | GTCTTGAGCAAATAGCAGGTGACAGGAGAAGATCTACCATGCAAAAATAAATAAAT | TCCATCTTGTCTGTTAGCAAGCTGGGAGAAGATCTACCATGCAAATTTATTTATTT |
| 216 | GTCTTGAGCAAATAGCAGGTGACAATTGAACGTGTTGTATCTGGAAATAAATAAAT | TCCATCTTGTCTGTTAGCAAGCTGATTGAACGTGTTGTATCTGGATTTATTTATTT |

**Supplementary Table 4.** Barcoded recording primers for pattern reconstruction experiments in Fig. 4.

| Direction | Barcode ID | Inner Primers (5' to 3') | Outer Primers (5' to 3') |
|---|---|---|---|
| Forward | 1 | AAGAAAGTTGTCGGTGTCTTTGTG GTCTTGAGCAAATAGCAGGTGACA | AAGAAAGTTGTCGGTGTCTTTGTG |
| Forward | 2 | TCGATTCCGTTTGTAGTCGTCTGT GTCTTGAGCAAATAGCAGGTGACA | TCGATTCCGTTTGTAGTCGTCTGT |
| Forward | 3 | GAGTCTTGTGTCCCAGTTACCAGG GTCTTGAGCAAATAGCAGGTGACA | GAGTCTTGTGTCCCAGTTACCAGG |
| Forward | 4 | TTCGGATTCTATCGTGTTTCCCTA GTCTTGAGCAAATAGCAGGTGACA | TTCGGATTCTATCGTGTTTCCCTA |
| Forward | 5 | CTTGTCCAGGGTTTGTGTAACCTT GTCTTGAGCAAATAGCAGGTGACA | CTTGTCCAGGGTTTGTGTAACCTT |
| Forward | 6 | TTCTCGCAAAGGCAGAAAGTAGTC GTCTTGAGCAAATAGCAGGTGACA | TTCTCGCAAAGGCAGAAAGTAGTC |
| Forward | 7 | GTGTTACCGTGGGAATGAATCCTT GTCTTGAGCAAATAGCAGGTGACA | GTGTTACCGTGGGAATGAATCCTT |
| Forward | 8 | TTCAGGGAACAAACCAAGTTACGT GTCTTGAGCAAATAGCAGGTGACA | TTCAGGGAACAAACCAAGTTACGT |
| Forward | 9 | AACTAGGCACAGCGAGTCTTGGTT GTCTTGAGCAAATAGCAGGTGACA | AACTAGGCACAGCGAGTCTTGGTT |
| Forward | 10 | AAGCGTTGAAACCTTTGTCCTCTC GTCTTGAGCAAATAGCAGGTGACA | AAGCGTTGAAACCTTTGTCCTCTC |
| Forward | 11 | GTTTCATCTATCGGAGGGAATGGA GTCTTGAGCAAATAGCAGGTGACA | GTTTCATCTATCGGAGGGAATGGA |
| Forward | 12 | CAGGTAGAAAGAAGCAGAATCGGA GTCTTGAGCAAATAGCAGGTGACA | CAGGTAGAAAGAAGCAGAATCGGA |
| Forward | 13 | AGAACGACTTCCATACTCGTGTGA GTCTTGAGCAAATAGCAGGTGACA | AGAACGACTTCCATACTCGTGTGA |
| Forward | 14 | AACGAGTCTCTTGGGACCCATAGA GTCTTGAGCAAATAGCAGGTGACA | AACGAGTCTCTTGGGACCCATAGA |
| Forward | 15 | AGGTCTACCTCGCTAACACCACTG GTCTTGAGCAAATAGCAGGTGACA | AGGTCTACCTCGCTAACACCACTG |
| Forward | 16 | CGTCAACTGACAGTGGTTCGTACT GTCTTGAGCAAATAGCAGGTGACA | CGTCAACTGACAGTGGTTCGTACT |
| Forward | 17 | ACCCTCCAGGAAAGTACCTCTGAT GTCTTGAGCAAATAGCAGGTGACA | ACCCTCCAGGAAAGTACCTCTGAT |
| Forward | 18 | CCAAACCCAACAACCTAGATAGGC GTCTTGAGCAAATAGCAGGTGACA | CCAAACCCAACAACCTAGATAGGC |
| Forward | 19 | GTTCCTCGTGCAGTGTCAAGAGAT GTCTTGAGCAAATAGCAGGTGACA | GTTCCTCGTGCAGTGTCAAGAGAT |
| Forward | 20 | TTGCGTCCTGTTACGAGAACTCAT GTCTTGAGCAAATAGCAGGTGACA | TTGCGTCCTGTTACGAGAACTCAT |
| Forward | 21 | GAGCCTCTCATTGTCCGTTCTCTA GTCTTGAGCAAATAGCAGGTGACA | GAGCCTCTCATTGTCCGTTCTCTA |
| Forward | 22 | ACCACTGCCATGTATCAAAGTACG GTCTTGAGCAAATAGCAGGTGACA | ACCACTGCCATGTATCAAAGTACG |
| Forward | 23 | CTTACTACCCAGTGAACCTCCTCG GTCTTGAGCAAATAGCAGGTGACA | CTTACTACCCAGTGAACCTCCTCG |
| Forward | 24 | GCATAGTTCTGCATGATGGGTTAG GTCTTGAGCAAATAGCAGGTGACA | GCATAGTTCTGCATGATGGGTTAG |
| Reverse | 25 | GTAAGTTGGGTATGCAACGCAATG TCCATCTTGTCTGTTAGCAAGCTG | GTAAGTTGGGTATGCAACGCAATG |
| Reverse | 26 | CATACAGCGACTACGCATTCTCAT TCCATCTTGTCTGTTAGCAAGCTG | CATACAGCGACTACGCATTCTCAT |
| Reverse | 27 | CGACGGTTAGATTCACCTCTTACA TCCATCTTGTCTGTTAGCAAGCTG | CGACGGTTAGATTCACCTCTTACA |
| Reverse | 28 | TGAAACCTAAGAAGGCACCGTATC TCCATCTTGTCTGTTAGCAAGCTG | TGAAACCTAAGAAGGCACCGTATC |
| Reverse | 29 | CTAGACACCTTGGGTTGACAGACC TCCATCTTGTCTGTTAGCAAGCTG | CTAGACACCTTGGGTTGACAGACC |

| Direction | Barcode ID | Inner Primers (5' to 3') | Outer Primers (5' to 3') |
|---|---|---|---|
| Reverse | 30 | TCAGTGAGGATCTACTTCGACCCA TCCATCTTGTCTGTTAGCAAGCTG | TCAGTGAGGATCTACTTCGACCCA |
| Reverse | 31 | TGCGTACAGCAATCAGTTACATTG TCCATCTTGTCTGTTAGCAAGCTG | TGCGTACAGCAATCAGTTACATTG |
| Reverse | 32 | CCAGTAGAAGTCCGACAACGTCAT TCCATCTTGTCTGTTAGCAAGCTG | CCAGTAGAAGTCCGACAACGTCAT |
| Reverse | 33 | CAGACTTGGTACGGTTGGGTAACT TCCATCTTGTCTGTTAGCAAGCTG | CAGACTTGGTACGGTTGGGTAACT |
| Reverse | 34 | GGACGAAGAACTCAAGTCAAAGGC TCCATCTTGTCTGTTAGCAAGCTG | GGACGAAGAACTCAAGTCAAAGGC |
| Reverse | 35 | CTACTTACGAAGCTGAGGGACTGC TCCATCTTGTCTGTTAGCAAGCTG | CTACTTACGAAGCTGAGGGACTGC |
| Reverse | 36 | ATGTCCCAGTTAGAGGAGGAAACA TCCATCTTGTCTGTTAGCAAGCTG | ATGTCCCAGTTAGAGGAGGAAACA |
| Reverse | 37 | GCTTGCGATTGATGCTTAGTATCA TCCATCTTGTCTGTTAGCAAGCTG | GCTTGCGATTGATGCTTAGTATCA |
| Reverse | 38 | ACCACAGGAGGACGATACAGAGAA TCCATCTTGTCTGTTAGCAAGCTG | ACCACAGGAGGACGATACAGAGAA |
| Reverse | 39 | CCACAGTGTCAACTAGAGCCTCTC TCCATCTTGTCTGTTAGCAAGCTG | CCACAGTGTCAACTAGAGCCTCTC |
| Reverse | 40 | TAGTTTGGATGACCAAGGATAGCC TCCATCTTGTCTGTTAGCAAGCTG | TAGTTTGGATGACCAAGGATAGCC |
| Reverse | 41 | GGAGTTCGTCCAGAGAAGTACACG TCCATCTTGTCTGTTAGCAAGCTG | GGAGTTCGTCCAGAGAAGTACACG |
| Reverse | 42 | CTACGTGTAAGGCATACCTGCCAG TCCATCTTGTCTGTTAGCAAGCTG | CTACGTGTAAGGCATACCTGCCAG |
| Reverse | 43 | CTTTCGTTGTTGACTCGACGGTAG TCCATCTTGTCTGTTAGCAAGCTG | CTTTCGTTGTTGACTCGACGGTAG |
| Reverse | 44 | AGTAGAAAGGGTTCCTTCCCACTC TCCATCTTGTCTGTTAGCAAGCTG | AGTAGAAAGGGTTCCTTCCCACTC |
| Reverse | 45 | GATCCAACAGAGATGCCTTCAGTG TCCATCTTGTCTGTTAGCAAGCTG | GATCCAACAGAGATGCCTTCAGTG |
| Reverse | 46 | GCTGTGTTCCACTTCATTCTCCTG TCCATCTTGTCTGTTAGCAAGCTG | GCTGTGTTCCACTTCATTCTCCTG |
| Reverse | 47 | GTGCAACTTTCCCACAGGTAGTTC TCCATCTTGTCTGTTAGCAAGCTG | GTGCAACTTTCCCACAGGTAGTTC |
| Reverse | 48 | CATCTGGAACGTGGTACACCTGTA TCCATCTTGTCTGTTAGCAAGCTG | CATCTGGAACGTGGTACACCTGTA |

**Supplementary Table 5.** Inner and outer PCR primers for DNA nanoscope experiments in Fig. 4 and Fig. S4. Inner and outer primers in the same row are designed to be used together to PCR barcoded recording primers. Any 'forward' inner and outer primer pair can be used with any 'reverse' inner and outer primer pair to generate a unique sequencing library. Multiple such distinct libraries can be combined and run on the same sequencing chip. The sequence of the outer primer is used to demultiplex libraries.

**References**

27. S. M. Douglas, A. H. Marblestone, S. Teerapittayanon, A. Vazquez, G. M. Church, W. M. Shih, Rapid prototyping of 3D DNA-origami shapes with caDNAno. *Nucleic Acids Res.* **37**, 5001–5006 (2009).
28. P. W. K. Rothemund, in *IEEE/ACM International Conference on Computer-Aided Design, Digest of Technical Papers, ICCAD* (2005), vol. 2005, pp. 470–477.
